# A mechanistic framework for the recognition of chemically diverse brassinosteroids by BRI1-family receptor kinases

**DOI:** 10.1101/2025.08.08.669299

**Authors:** Alberto Caregnato, Houming Chen, Miroslav Kvasnica, Ulrich Hohmann, Jana Oklestkova, Karoll Ferrer, Larissa Broger, Ludwig A. Hothorn, Miroslav Strnad, Michael Hothorn

**Affiliations:** Structural Plant Biology Laboratory, Department of Plant Science, University of Geneva, 1211 Geneva, Switzerland; Laboratory of Growth Regulators, Faculty of Science, Palacky University & Institute of Experimental Botany, The Czech Academy of Sciences, CZ-77900, Olomouc, Czech Republic; Institute of Molecular Biotechnology of the Austrian Academy of Sciences (IMBA) & Research Institute of Molecular Pathology (IMP), Vienna BioCenter (VBC), 1030, Vienna, Austria; Retired from Leibniz University Hannover, Germany, present address: Im Grund 12, 31867 Lauenau, Germany

## Abstract

Brassinosteroids (BRs) are chemically diverse plant steroid hormones produced via a branched biosynthetic pathway. The potent BR brassinolide is sensed by the membrane receptor kinase BRIl and a SERK co-receptor, but the physiological functions of other abundant BRs remain to be characterized. Here, we present quantitative binding kinetics for four *Arabidopsis thaliana* BR receptors and fifteen BRs, which define the key chemical features required for high affinity receptor binding, ligand positioning, and co-receptor recognition. BRIl, BRLl, and BRL3 share overlapping ligand preferences, whereas BRL2 preferentially binds C_28_ BRs with moderate affinity. Structural analyses of BR-bound BRIl and BRL3 ectodomains, combined with extensive in vitro and in vivo mutagenesis studies reveal a high structural plasticity of the hormone binding pocket. Functional assays using structure-based BR agonists and antagonists uncover that BR receptor - co-receptor signaling complexes can recognize chemically diverse BRs, introducing an additional, intriguing layer of BR signaling regulation.

## Introduction

Brassinosteroids (BRs) are a class of lipophilic, polyhydroxylated steroid hormones first isolated from rapeseed pollen and later recognized for their growth-promoting roles in plants^l^. BRs are synthesized via three main biosynthetic branches, producing C_27_, C_28_ or C_29_ compounds. While all share a steroidal ring core, they differ in the structure of their aliphatic side chains^2^ (Fig. la). The C_27_ branch, derived from cholesterol, yields 28-norbrassinolide^3,4^, whereas the C_29_ branch, from sitosterol, produces 28-homocastasterone and 28-homobrassinolide (Fig. lb). The C_28_ pathway originates from campesterol and leads to brassinolide, one of the most bioactive BRs^2^ (Fig. lb). The C_28_ pathway includes two routes: early and late C_22_ oxidation (Fig. lb). In the early pathway, DWARF4 (DWF4), a cytochrome P450 monooxygenase, converts campesterol to (22*S*)-22-hydroxycampesterol^3,5,6^. Further steps involve the P450 enzymes CPD^7,8^ and ROT3^9^, and the steroid-5a-reductase DET2^10-12^, producing 6-deoxocastasterone (Fig. lb). The enzymes BR6oxl and BR6ox2 then convert this intermediate to castasterone^l3^, and BR6ox2 can further catalyze formation of a seven-membered lactone ring from the C_6_ ketone, yielding brassinolide^l3^ (Fig. lb).

**Fig. 1.**
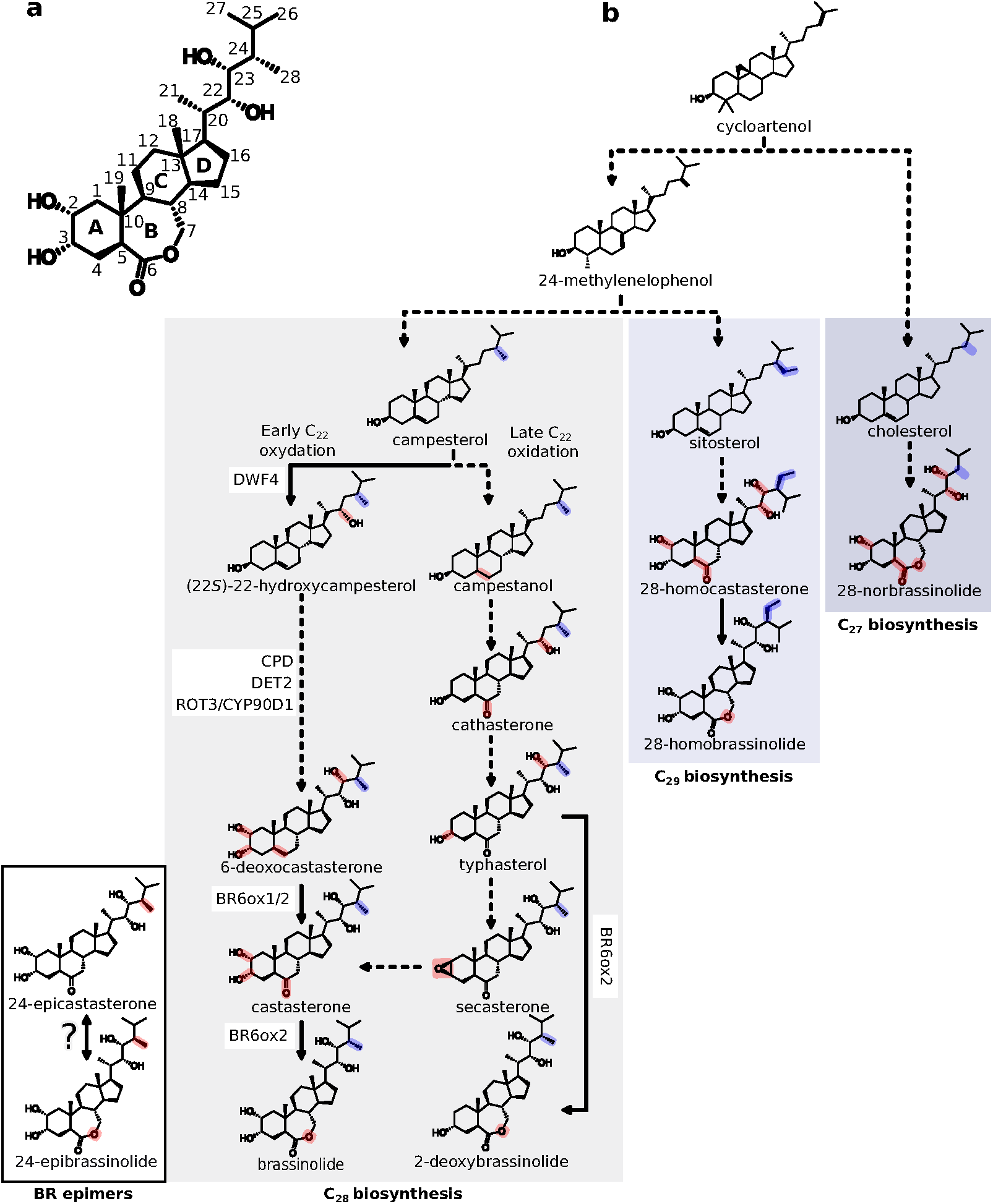
I The branched architecture of the BR biosynthesis pathway. **a**, Chemical structure of brassinolide with labeled steroidal rings (letters) and carbon atoms (numbers). **b**, Schematic representation of the BR biosynthesis pathway, highlighting the relative positions of the BRs discussed in this study. The C_28_, C_29_, and C_27_ branches are shaded from light to dark gray, respectively. Key chemical differences between branches are indicated in blue, while red marks chemical modifications along each biosynthetic route. Solid lines denote single-step enzymatic reactions; dotted lines indicate multi-step conversions. Known enzymes involved in the canonical C_28_ pathway are shown alongside.

In the late C_22_ oxidation pathway, campestanol is processed into cathasterone and typhasterol^l4^ (Fig. lb). BR6ox2 can convert typhasterol into 2-deoxybrassinolide^l5^, or alternatively, typhasterol may be converted to castasterone via secasterone, a 2,3-epoxy intermediate^l6^ (Fig. lb). The different branches of BR biosynthesis are likely interconnected^2^. BR homeostasis is regulated by several catabolic enzymes, including the P450 enzyme BASl^l7^, the UDP-glycosyltransferase UGT73C5^l8^, and sulfotransferases^l9^.

BR biosynthesis occurs, at least in part, within the endoplasmic reticulum^20,21^. Local BR transport is facilitated via plasmodesmata^22^. The ATP-binding cassette transporters ABCBl and ABCB19 are responsible for translocating BRs across the plasma membrane from the cytosol into the apoplast^23^, where BR perception takes place^24^.

Brassinosteroids (BRs) are perceived by the plasma membrane-localized receptor kinases BRASSINOSTEROID INSENSITIVE l (BRIl)^24–28^, BRLl^29–31^ and BRL3^29^, which share a conserved extracellular leucine-rich repeat (LRR) ligand binding domain. BRIl is broadly expressed, and *bril* loss-of-function mutants display a characteristic dwarf phenotype^25^. BRLl and BRL3 are present in vascular tissues and can rescue the growth defects of the weak *bril-301* allele^32^, when expressed from the *BRIl* promoter^29^. Expression of the sequence-related BRL2 could not rescue the *bril-301* mutant phenotype^29^, or a *bril brll brl3* triple mutant^33^. BRL2 is also known as VASCULAR HIGHWAYl (VHl) and has been implicated in leaf vein patterning^34^.BR_ligand_recognition_MH13.odt BRs bind to a steroid-binding pocket formed by the LRR core and a ∼70 amino acid island domain within BRIl or BRLl^26,27,31^. BR binding enables BRIl to recruit SOMATIC EMBRYOGENESIS RECEPTOR KINASEs (SERKs) to form an active signaling complex^26,35,36^. In the absence of BRs, SERKs are sequestered by BIR pseudokinases^37,38^. Assembly of the BRIl-BR-SERK complex activates the cytoplasmic kinase domains^39^, triggering a downstream signaling cascade that leads to dephosphorylation of the BR transcription factors BESl and BZRl^40,41^.

Bioassays with synthetic, exogenously applied brassinosteroids (BRs) have been performed across various plant species. In Arabidopsis, nanomolar levels of brassinolide promote hypocotyl elongation in both wild-type and *det2* mutant seedlings^42^. Brassinolide or 24-epibrassinolide applied at high nanomolar to micromolar concentrations inhibits root growth^43^, whereas low concentrations - high picomolar to low nanomolar - of 24-epibrassinolide or 24-epicastasterone stimulate root growth^44^.

Initial BR-binding assays used microsomal fractions from transgenic Arabidopsis expressing BRIl-GFP and [3H]-brassinolide, yielding estimated dissociation constants (K_d_) of ∼10-15 nM for brassinolide and 40-75 nM for castasterone^24^. BRLl and BRL3 bound brassinolide with K_d_ values of ∼5 nM and ∼50 nM, respectively, while BRL2 showed no detectable binding^29^. Grating-coupled interferometry (GCI) quantified bBR_ligand_recognition_MH12.odtinding of unlabeled brassinolide to BRIl with a K_d_ of ∼10 nM^28^. Brassinolide-bound BRIl enables the binding of the SERK3/BAKl LRR ectodomain, with K_d_ values of ∼0.2-l µM, as determined by GCI and isothermal titration calorimetry^28,36^. Mutation of Gly644---Asp (*bril-6*/*bril-119*)^45^ in the BR island domain^26^ reduces BR binding in vitro^28,46^. The corresponding *bril-6* allele exhibits intermediate growth defects^45^. Mutation of neighboring Gly643---Glu in the gain-of-function BRIl*^sudl^* allele^47^, enhances the structural stability of the island domain^26^. Notably, five simultaneous point mutations in the BR-binding pocket were required to completely abolish brassinolide binding in vitro, and to mimic the severe growth phenotype of *bril* null alleles in vivo^46^.

The natural occurrence of different BRs has been characterized using high-performance liquid chromatography coupled with gas chromatography-mass spectrometry. In Arabidopsis shoots, castasterone, 6-deoxocastasterone, and typhasterol were identified^ll,48,49^. Like in other species, brassinolide accumulates more prominently in Arabidopsis seeds^50^.

Brassinolide, the most bioactive BR^2^, is widely considered the primary signaling ligand for BRIl, BRLl, and BRL3. Consequently, prior structural and biochemical investigations have focused on brassinolide^24,26–29,31,36,46^. Given the chemical diversity of BRs from the C_27_, C_28_, and C_29_ biosynthetic branches that accumulate in plants, we set out to systematically characterize their binding kinetics to the ectodomains of the *Arabidopsis thaliana* BR receptors BRIl, BRLl, BRL2, and BRL3. By integrating these measurements with structural and mutational analyses of the BRIl binding pocket - both in vitro and in planta - as well as targeted BR bioassays, we define the ligand-binding specificities of each receptor and uncover the chemical features in BRs required for receptor binding and co-receptor recruitment to form an active signaling complex.

## Results

### BRIl, BRLl and BRL3 bind chemically diverse BRs with high affinity

To investigate the ligand-binding specificity of Arabidopsis BR receptors, we obtained the LRR ectodomains of BRIl, BRLl, and BRL3 by secreted expression in baculovirus-infected insect cells, as described^26,28,46^. Quantitative GCI kinetic binding assays confirmed^28,29^ that BRIl, BRLl and BRL3 bind brassinolide with nanomolar affinity (Fig. 2a, Extended Data Figs. 1,2, Extended Data Table l). We then determined the crystal structure of the BRL3 ectodomain in complex with brassinolide at 2.45 A resolution (Fig. 2b and Supplementary Fig. 1 and Supplementary Table l). BRIl^26,27^, BRLl^31^ and BRL3 share a highly conserved ligand binding pocket (root mean square deviation, r.m.s.d. is ∼0.7 A comparing 180 corresponding C_a_ atoms, Supplementary Fig. 1a), formed by conserved hydrophobic residues along the inner LRR surface and island domain (Fig. 2b). The brassinolide C_23_ hydroxyl forms a hydrogen bond with a main chain amino group in the island domain of all three receptors (Fig. 2b and Supplementary Fig. 1a). Few receptor-specific polar contacts with the ligand are observed. In BRIl, Tyr597 forms hydrogen bonds with the brassinolide C_22_ and C_23_ hydroxyls via a water molecule (Fig. 2b), whereas in BRLl and BRL3, Tyr597 is replaced by phenylalanine. In the BRL3 complex, Arg588 (corresponding to Lys601 in BRIl) and Tyr627 form hydrogen bonds BR_ligand_recognition_MH13.odtwith the C_6_ hydroxyl group of brassinolide (Fig. 2b). BRL3 Gln666 (Phe681 in BRIl) coordinates the C_2_ hydroxyl via a water molecule (Fig. 2b). Thus, BRIl, BRLl, and BRL3 share a structurally conserved BR-binding pocket with limited receptor-specific features, and all bind brassinolide with high affinity.

**Fig. 2.**
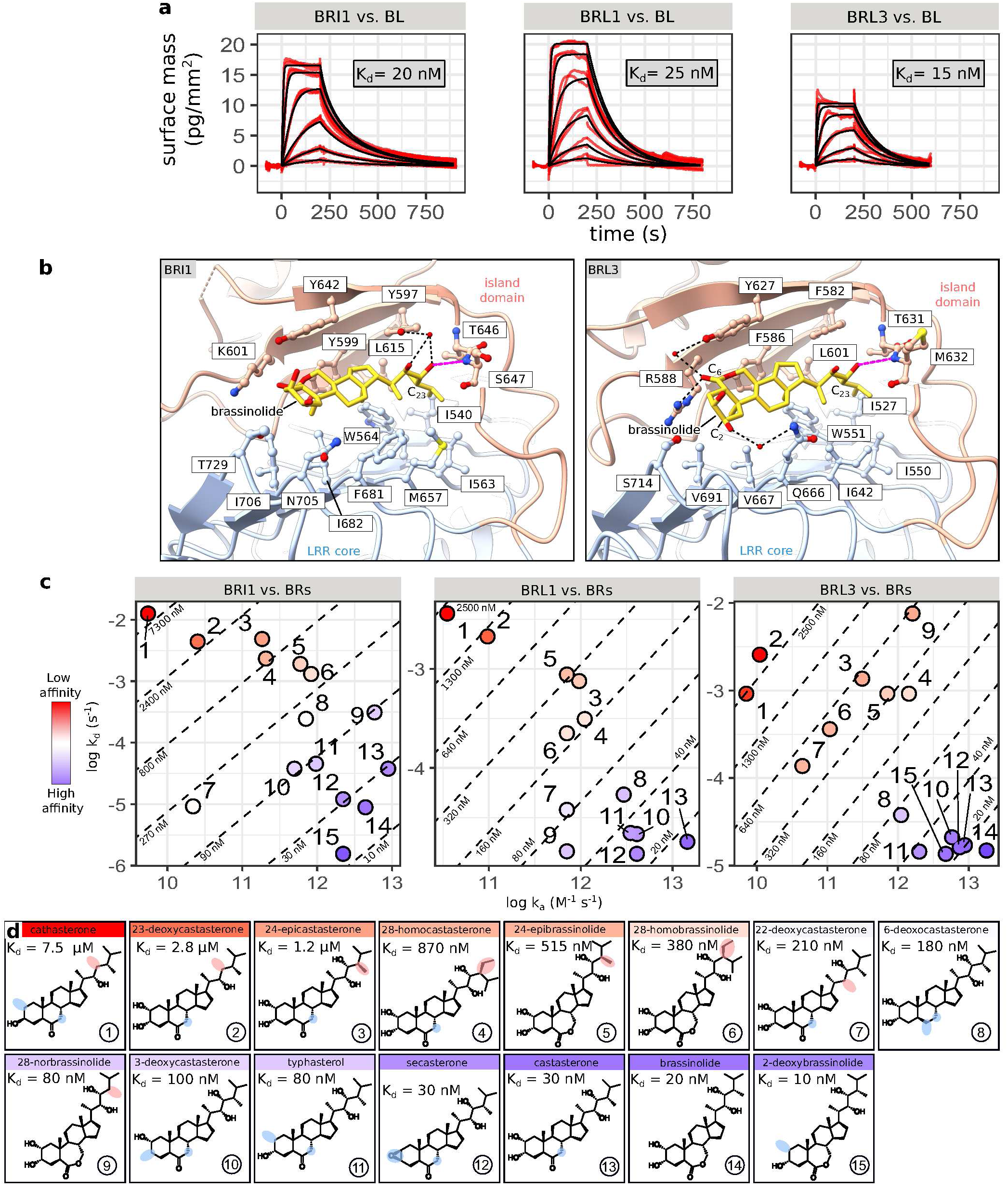
I BRil, BRLl and BRL3 bind chemically diverse BRs. **a,** GCI analysis of ligand-binding kinetics between BRIl, BRLl, BRL3 and brassinolide. Shown are sensorgrams (n=2, red) with global fits (black) and equilibrium dissociation constants (K_d_) indicated. **b,** BR-binding pockets of BRIl (left; PDB-ID: pdb_00003rj0) and BRL3 (right) shown as ribbon diagrams. LRRs in blue, island domains in red, brassinolide in yellow. Pocket residues and hydrogen bonds (dotted lines) shown; conserved C_23_-OH hydrogen bond highlighted in purple. **c,** Isoaffinity plots of association (k_a_) and dissociation (k_d_) rates for receptor - BR pairs. Each point (n = 2) reflects a global fit; diagonal lines denote K_d_. High- and low-affinity interactions are in blue and red, respectively. **d,** Structures of tested BRs, numbered as in **c**, with BRIl K_d_ values. Red highlights indicate modifications that reduce binding, while blue highlights indicate modifications that have a minimal effect.

Next, we quantified the interaction of BRIl, BRLl, and BRL3 with 12 natural BRs from the three biosynthetic branches, along with synthetic compounds 3-deoxycastasterone, 22-deoxycastasterone, and 23-deoxycastasterone (Fig. 2c,d). BRIl displayed binding constants ranging from low micromolar (red in k_a_/k_d_ plots, Fig. 2c) to low nanomolar (blue in Fig. 2c, Extended Data Figs. 1,2, Extended Data Table l). Low-affinity ligands include cathasterone (an early C_28_ precursor), 24-epibrassinolide and 24-epicastasterone (late C_28_ branch), and the C_29_ products 28-homocastasterone and 28-homobrassinolide (Fig. lb, Fig. 2c,d). 28-norbrassinolide (C_27_ branch), as well as brassinolide and castasterone (late C_28_ branch) represent high affinity ligands (Fig. lb and Fig. 2c,d). Additional high-affinity binders include typhasterol, secasterone, and 2-deoxybrassinolide (late C_22_ oxidation), as well as 6-deoxocastasterone (early C_22_ oxidation), all within the C_28_ branch (Fig. lb, Fig. 2c,d). BRLl and BRL3 share highly similar ligand binding specificity profiles with BRIl (Fig. 2c, Extended Data Figs. 1,2, Extended Data Table l), consistent with their structurally conserved BR-binding pockets (Fig. 2b, Supplementary Fig. 1) and their ability to rescue *bril-301* mutant phenotypes^29^.

We next sought to define the chemical features of BRs that contribute to high-affinity receptor binding. Side chain modifications - such as the C_24_ ethyl substitution in 28-homocastasterone or 28-homobrassinolide (Fig. la,b) - were associated with ∼10- to 30-fold and ∼10- to 35-fold reductions in binding affinity, respectively, compared to castasterone and brassinolide (Fig. 2c,d). Similarly, the brassinolide / castasterone enantiomers 24-epibrassinolide and 24-epicastasterone (Fig. la,b) showed ∼10- to 20-fold and ∼10- to 40-fold reduced affinities (Fig. 2c,d). Absence of a hydroxyl group at C_23_ - as in cathasterone - correlated with weak receptor binding (Fig. lb, Fig. 2c,d).

To further assess the role of side chain hydroxylation, we synthesized 23-deoxycastasterone (Supplementary Note l). 23-deoxycastasterone bound BRIl with a dissociation constant of ∼3 µM, representing a 100-fold reduction compared to castasterone (Fig. 2c,d). In contrast, 22- deoxycastasterone - lacking the C_22_ hydroxyl group present in castasterone and brassinolide (Fig. lb) - still bound BRIl with medium to high affinity (K_d_ ∼200 nM, Fig. 2c,d). These results are consistent with our structural data showing that the C_23_, but not the C_22_ hydroxyl group forms conserved hydrogen bonds with the island domains of BRIl, BRLl, and BRL3 (Fig. 2b, Supplementary Fig. 1a).

Ring modifications were generally well tolerated by the BRIl, BRLl, and BRL3 ectodomains (Fig. 2c,d). Lactonization of the B-ring had minimal impact on binding, as brassinolide and castasterone showed nearly identical dissociation constants (Fig. 2c,d). Although C_6_ oxidation is a key biosynthetic step towards brassinolide^l3,51^, 6-deoxocastasterone bound all three receptors with high affinity (Fig. 2c,d). Supporting this, the complex structures of BRIl, BRLl, and BRL3 show no conserved interactions with the C_6_ ketone^26,27,31^ (Fig. 2b, Supplementary Fig. 1a). The 2a and 3a hydroxyl groups in the A-ring also appear dispensable for receptor binding, as ligands lacking either - such as 2-deoxybrassinolide, 3-deoxycastasterone, typhasterol, and secasterone - bound BRIl, BRLl, and BRL3 with nanomolar affinity (Fig. 2c,d).

Overall, our quantitative binding data demonstrate that BR receptors recognize a broad range of structurally diverse BRs with high affinity. High-affinity receptor binding is driven by the configuration and size of the aliphatic side chain - most critically, the presence of a C_23_ hydroxyl group and the correct C_24_ stereochemistry.

### High structural plasticity of the hormone binding pocket enables BRIl/BRLs to sense different BRs

To clarify the structural basis of ligand-binding specificity in BR receptors, we co- crystallized the BRIl ectodomain with various BRs. Crystals grown with cathasterone, 24- epicastasterone, or 28-homocastasterone showed no interpretable difference electron density in the BR-binding pocket, while 6-deoxocastasterone yielded poorly diffracting crystals. A 2.l A structure of the BRIl - castasterone complex revealed a ligand conformation nearly identical to that of brassinolide (Fig. 3a,b, Supplementary Table l), consistent with its high binding affinity (Fig. 2c,d), and confirming that B-ring lactonization is not essential for receptor recognition. Similarly, a 2.7 A structure of the BRIl - typhasterol complex showed that the absence of a C_2_ hydroxyl group does not affect ligand coordination by the receptor (Fig. 3a,b). In contrast, the 2.4 A BRIl - 24- epibrassinolide structure exhibited weak electron density for the BR side chain, consistent with its reduced binding affinity (Fig. 2c,d, Fig. 3d). Finally, the 2.4 A BRIl - 28-homobrassinolide complex revealed that the receptor can in principle accommodate a C_24_ ethyl substitution in its binding pocket (Fig. 3a,b).

**Fig. 3.**
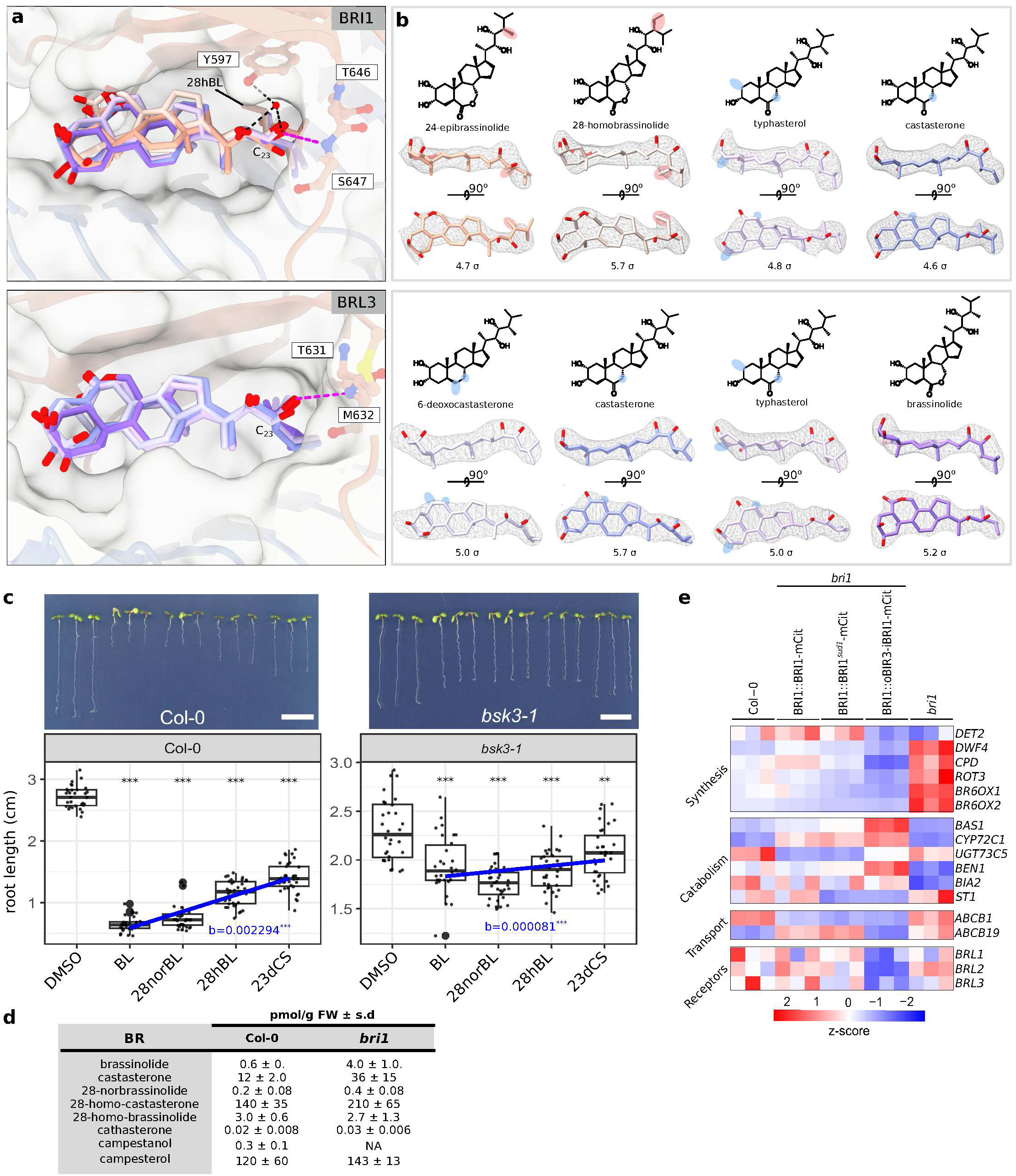
I The hormone binding pocket can accommodate structurally diverse BRs. **a,** Structural superposition of different BRIl - BR (top, r.m.s.d. ∼0.4 A comparing 186 corresponding C_a_ atoms) and BRL3 - BR complexes (bottom, ∼r.m.s.d. ∼0.3 A comparing 190 corresponding C_a_ atoms). The receptors are shown in combined ribbon diagram (LRR domain in light blue, island domain in light red) and molecular surface views. BRs are in bonds representation color-coded according to their receptor binding affinities, ranging from low (light red) to high affinity (purple; BRIl - brassinolide complex PDB-ID: pdb_00003rj0). Dashed lines highlight key polar interactions among the ligand and the protein receptor binding pockets. A conserved hydrogen bond between brassinolide and the island domains of BRIl, BRLl, and BRL3 is marked in magenta. **b,** Structures of BRs (in bonds representation, color-coded as in **a** with BRIl (top) or BRL3 (bottom) and including final mF_o_-DF_c_ omit electron density maps (gray mesh, contoured at the indicated o level). The respective chemical structures are shown above, with modifications relative to brassinolide marked in red (for changes that reduce binding affinity), or blue (neutral). **c,** Root growth inhibition assay of 7-d-old Col-0 wild-type and *bsk3-l*^79^ mutant seedlings (n=31-35) grown on ^l/2^MS plates supplemented with the indicated BR (brassinolide, BL; 28-norbrassinolide, 28norBL; 28-homobrassinolide, 28hBL; 23- deoxycastasterone, 23dCS, 0.l% [v/v] DMSO as control) at a final concentration of 100 nM (scale bar = l cm). Quantifications (n=30) are shown as box plots (centre line, median; box, interquartile range IQR; whiskers, lowest/highest data point within l.5 IQR of the lower/upper quartile, raw data shown a gray dots). Multiple comparison of the different treatments vs. the DMSO control were performed using a Dunnett^91^ test as implemented in the R package multcomp^92^ (\**p* < 0.05; \*\**p* < 0.01; \*\*\**p* < 0.001). The relationship between the estimated dissociation constant (K_d_) for each ligand and root length were tested using a Tukey trend test^93,94^. The blue line represent the linear trend with slope (b) and the associated test p-value (\**p* < 0.05; \*\**p* < 0.01; \*\*\**p* < 0.001). **d**, BR quantification in 10 d old Col-0 and *bril* (GABI_134E10^47^) seedlings (FW, fresh weight). **e**, RNA- seq derived expression profile heat map of BR biosynthesis, catabolism, transport and receptor genes in 10 d old seedlings in untreated BR signaling mutants and the Col-0 wild type. Expression values were transformed into Z-scores based on the normalized counts, three biological replicates are shown in three horizontal rows, the color scale indicates expression level (red, up-regulation, blue, down-regulation).

We next resolved the crystal structures of the BRL3 ectodomain bound to castasterone and typhasterol at 2.6 A and 2.3 A, respectively. In both structures, the ligands adopt conformations closely resembling that of brassinolide (Fig. 3a,b). A 3.2 A structure of the BRL3 - 6- deoxocastasterone complex shows that, despite lacking B-ring modifications, the ligand is accommodated in the binding pocket, albeit in a shifted conformation (Fig. 3a,b). Notably, in all BR receptor - ligand complexes analyzed, the BR C_23_ hydroxyl group consistently forms a conserved hydrogen bond with the main chain amino group of Ser647 in BRIl or Met632 in BRLl/BRL3 (Fig. 3a).

Several catabolic enzymes contribute to BR homeostasis, including BASl, which hydroxylates side chain C_26_^l7^, UGT73C5, which glucosylates the hydroxyl group at C_23_^l8^, and STl/ST4a, which sulfonylates the C_22_ hydroxyl^l9^. Structural analysis of our receptor-ligand complexes indicates that such tail modifications are not compatible with the BR-binding pocket (Supplementary Fig. 2). Together, the BRIl and BRL3 ligand-bound structures rationalize the BR binding kinetics data, emphasizing the importance of the BR side chain for high-affinity receptor binding, while showing that modifications to the steroid ring core are generally tolerated.

To evaluate the physiological relevance of our biochemical and structural experiments, we selected ligands representing high, medium, and low-affinity binders for root growth inhibition assays, as previously described^43^. Brassinolide (K_d_ ∼20 nM) and 28-norbrassinolide (K_d_ ∼80 nM) were tested as high-affinity binders, 28-homobrassinolide (K_d_ ∼380 nM) as medium-affinity, and 23-deoxycastasterone (K_d_ ∼2800 nM) as a low-affinity binder (Fig. 2d). Root growth inhibition assays were performed using each compound at a final concentration of 100 nM. A strong trend for increasing root length with increasing dissociation constant (K_d_) was observed in treated wild-type seedlings, and a weaker trend in the *bsk3-l* control (Fig. 3c, Extended Data Table 2), suggesting that in vivo BR bioactivities correlate well with their receptor binding affinity in vitro. Notably, even the low-affinity binder 23-deoxycastasterone significantly inhibited root growth of wild-type seedlings at a concentration of 100 nM (Fig. 3c).

We next correlated in vitro BR receptor binding affinities with BR levels in planta. We quantified BR levels in 10-d-old Arabidopsis plants using UHPLC-MS/MS (see Methods). In whole wild-type seedlings, brassinolide was detected at low levels (∼0.6 pmol/g fresh weight), while castasterone levels were ∼20-fold higher (Fig. 3d). 28-norbrassinolide was also present at low concentrations (∼0.2 pmol/g FW), whereas 28-homocastasterone was abundant (∼140 pmol/g FW) (Fig. 3d). Correlating these endogenous BR levels with their respective in vitro binding affinities (Fig. 2c,d), suggests that, in addition to brassinolide, other BRs - such as castasterone, 28- norbrassinolide, and 28-homocastasterone - may act as physiologically relevant ligands for BRIl, BRLl, and BRL3.

### Constitutive activation or inactivation of BRIl modulates BR homeostasis

We next compared BR levels in wild-type and *bril* null mutant seedlings^47^. In the *bril* mutant, brassinolide and castasterone levels were elevated by ∼7-fold and ∼3-fold, respectively, compared to wild type (Fig. 3d), as previously described^45^. This aligns with earlier findings that BR metabolic and catabolic gene expression is altered in *bril* loss-of-function mutants^52^, partly mediated by the BR-regulated transcription factor BZRl^40,53^. To further investigate, we performed RNA-seq on untreated 10-d-old Col-0 seedlings, a *bril* null mutant^47^, and in *bril* complementation lines expressing either a wild-type BRIl receptor^47^, the BRIl*^sudl^* gain-of-function allele^47,36^, or a constitutively active, ligand-independent oBIR3-iBRIl receptor chimera^54^. We found many BR biosynthesis genes to be mildly induced in *bril* mutant seedlings and to be repressed in the constitutively active oBIR3-iBRIl^54^ line (Fig. 3e and Extended Data Table 3). BR biosynthesis genes were up-regulated in *bril* and down-regulated in oBIR3-iBRIl seedlings (Fig. 3e, Extended Data Table 3). Conversely, several BR catabolic genes were repressed in *bril* and induced in oBIR3-iBRIl plants (Fig. 3e). ABCBl and ABCB19, which function as BR transporters required for signaling^23^, also showed differential expression in *bril* and oBIR3-iBRIl seedlings (Fig. 3e). Notably, *BRLl*, *BRL2*, and *BRL3* transcript levels were reduced in oBIR3-iBRIl and partially induced in *bril* (Fig. 3e). Expression of a wild-type BRIl receptor restored gene expression in *bril* mutants to wild-type levels (Fig. 3e). The gain-of-function allele BRIl*^sudl^* mimicked the effects of the constitutively active oBIR3-iBRIl receptor chimera, albeit with less pronounced effects (Fig. 3e). Together, these experiments indicate that BRIl signaling activity shapes BR homeostasis by modulating the expression of BR metabolic, catabolic, and transport genes.

### BRL2 is a medium-affinity BR receptor

The structural flexibility of the BR-binding pockets in BRIl, BRLl, and BRL3, along with altered BRL2 expression in BRIl signaling mutants (Fig. 3e), prompted us to re-examine BRL2’s ligand-binding properties. BRIl and BRL2 share ∼40% sequence identity in their extracellular domains. We expressed and purified the BRL2 ectodomain from insect cells and solved its apo crystal structure to 2.6 A resolution (see Methods). The N-glycosylated BRL2 adopts a superhelical architecture with 21 LRRs (compared to 25 in BRIl) with a more open conformation (Supplementary Fig. 3a). BRL2’s island domain is partially disordered and participates in a domain- swapped crystallographic dimer (Supplementary Fig. 3a).

The ligand-binding pocket of BRL2 exhibits a high degree of structural similarity with the BR-binding sites of BRIl, BRLl, and BRL3 (Fig. 4a, Supplementary Fig. 3b). Using GCI assays, we found that BRL2 can bind BRs, but it has a distinct ligand binding profile compared to BRIl, BRLl and BRL3 (Fig 4b,c, Fig. 2c,d). BRL2 bound brassinolide and castasterone with K_d_ values of ∼200 nM and ∼400 nM, respectively (Fig. 4b,c; Extended Data Fig. 3, Extended Data Table 4) - 10- 30-fold weaker than BRIl (Fig. 2c,d). Unlike BRIl, which binds typhasterol and 28-norbrassinolide with nanomolar affinity, BRL2 bound these ligands only weakly (Fig. 4b,c). BRL2 showed no detectable binding to 28-homocastasterone, 24-epicastasterone, 24-epibrassinolide, or to dehydroepiandrosterone (DHEA), triolone, lupeol, or betulin (Fig. 4b,c). Our experiments suggest that BRL2 functions as a medium-affinity receptor for C_28_ BRs (Fig. lb). In line with this, the BR- binding pocket of BRIl, BRLl/3 and BRL2 share a high degree of structural and sequence conservation. The only notable exception is Glu614 in BRL2, which corresponds to Met657 and Ile642 in BRIl and BRL3, respectively (Fig. 4a, Supplementary Fig. 3b).

**Fig. 4.**
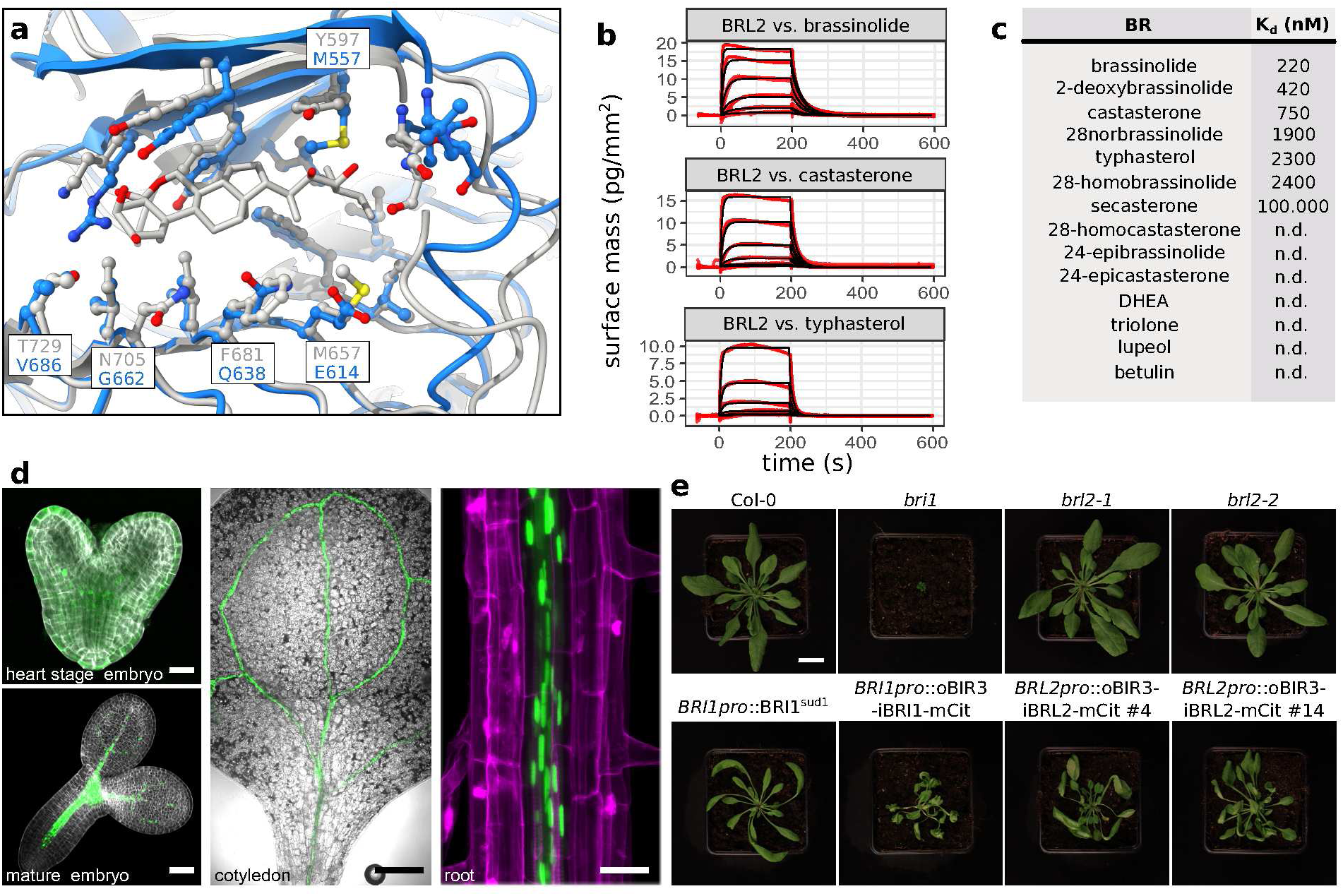
I BRL2 is a medium-affinity BR receptor. **a,** Structural superposition of the BRIl ectodomain (ribbon diagram, in gray; PDB-ID: pdb_00003rj0) with the AlphaFold2 model of the apo BRL2 ectodomain (in blue, AlphaFoldDB-ID: AF-Q9ZPS9-Fl, r.m.s.d. ∼ 0.8 A comparing 163 C_a_ atoms). Brassinolide is in bonds representation, interacting residue are shown as ball-and-sticks. **b,** GCI-derived binding kinetics of the BRL2 ectodomain vs. brassinolide, castasterone and typhasterol. Red traces represent experimental sensorgrams (n=2), and black lines show the global fits. **c,** Dissociation constant (K_d_) table of the BRL2 ectodomain vs. different BRs, non-BR steroids and triterpenpoids (n.d., no binding detected). **d,** Confocal images of a representative *BRL2prom*::NLS-3xmVenus reporter (in green) line in different stages: heart-stage embryo, mature embryo (cell wall stained with Renaissance SR2200, gray, scale bar = 20 µm), cotyledon (scale bar = 200 µm) and root from a 10-d-old seedling (cell wall stained with propidium iodide, magenta, scale bar = 30 µm). **e,** Rosette phenotypes of 5-week-old plants, germinated on ^l/2^MS for 2 weeks before transferring to soil for additional 3 weeks (scale bar = 2 cm).

We generated a *BRL2prom*::NLS-3xmVenus reporter line in Col-0 background and found *BRL2* to be expressed in vascular tissues across different developmental stages (Fig. 4d). To investigate its function, we analyzed the *brl2-l* loss-of-function mutant alongside a *bril* null mutant^47^. Unlike *bril*, *brl2-l* showed no dwarf phenotype when grown in soil (Fig. 4e). A second loss-of-function allele, *brl2-2*, created by CRISPR/Cas9 gene editing (Supplementary Fig. 3c), also showed no visible growth defects (Fig. 4e). Although prior work reported vein patterning defects inVHl/BRL2 over-expression^34^ and *brl2-l* loss-of-function^55^ lines, a quantitative analysis of leaf vein patterns in *brl2-l* did not reveal statistically significant differences compared to the Col-0 wild-type control (Supplementary Fig. 3d). *brl2-l* mutants also showed wild-type responses in hypocotyl growth assays (Supplementary Fig. 3e) and normal BESl dephosphorylation in response to brassinolide (Supplementary Fig. 3f).

We have previously reported that receptor chimeras, in which the extracellular and transmembrane domains of BIR3 (oBIR3) are fused to the cytoplasmic kinase domain of BRIl (iBRIl), trigger constitutive, ligand-independent BR responses^38,54^. To assess the signaling capacity of BRL2, we expressed a *BRL2prom*::oBIR3-iBRL2 chimera in Col-0 wild-type background. The resulting transgenics displayed elongated, slender petioles and curly leaves, similar to BRIl*^sudl^* gain- of-function mutants^24,47^ (Fig. 4e). However, their rosette phenotypes were milder than those of *BRIlprom*::oBIR3-iBRIl chimeras^54^ (Fig. 4e). Taken together, BRL2 is a vascular-localized, medium-affinity BR receptor that, like BRLl and BRL3^29,30^, does not exhibit strong loss-of-function phenotypes.

### A mutational map of the BR-binding pocket rationalizes the specificity and plasticity of BR receptors

Given the high structural conservation of the BR-binding site in BRIl, BRL2, BRLl and BRL3 (Supplementary Fig. 1b, Supplementary Fig. 4a), we examined the role of individual pocket- lining residues in BR perception. Based on our BRIl - BR complex structures (Fig. 3a), we mutated eleven residues in the LRR core and island domain (Fig. 5a) and measured brassinolide binding kinetics for each mutant ectodomain (Fig. 5b,c, Extended Data Fig. 4, and Extended Table 5). All mutants behaved as well-folded monomers in solution (Supplementary Fig. 4b).

**Fig. 5.**
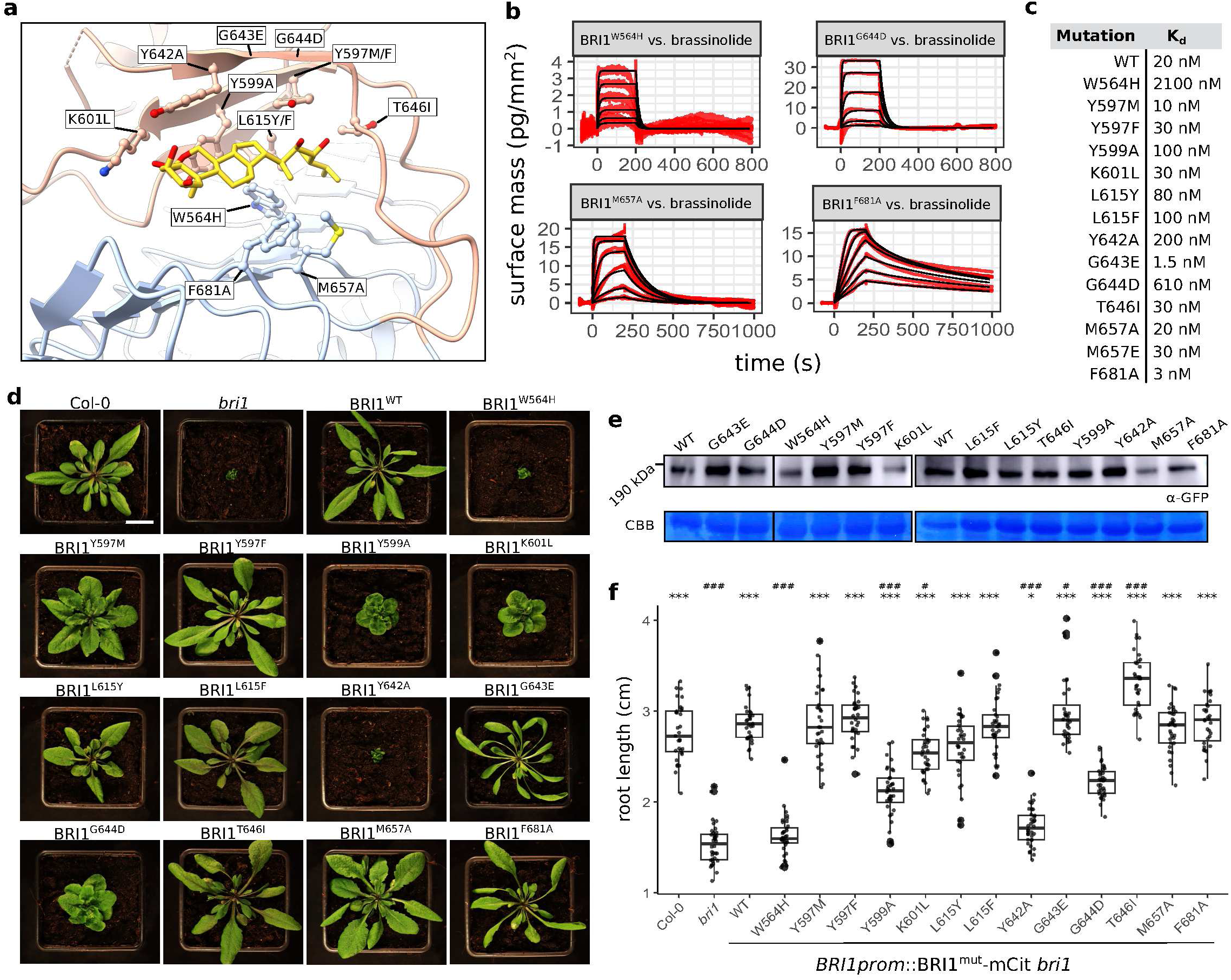
I Conserved aromatic residues in the hormone binding pocket are critical for BR perception. **a,** Ribbon diagram of BRIl (LRR domain in blue, island domain in red) in complex with brassinolide (in bonds representation, PDB-ID: pdb_00003rj0). Mutated residues are displayed as ball-and-sticks. **b,** GCI-derived binding kinetics for BRIl^W564H^, BRIl^G644D^, BRIl^M657A^ and BRIl^F681A^ vs. brassinolide. Red traces represent experimental data (n=2), black lines show the global fits. **c,** Dissociation constant (K_d_) table for BRIl point mutants vs. brassinolide. **d,** Rosette phenotypes of representative, 5-week-old transgenic T3 *BRIlprom*::BRIl-Citrine^mut^ complementation lines in *bril* null^47^ background compared to the Col-0 wild-type control (scale bar = 2 cm). **e,** Quantification of BRIl^mut^-mCitrine (theoretical molecular mass ∼ 158.6 kDa) protein levels by western blot from 10-d-old seedling protein extracts, immuno-blotted with an anti-GFP antibody. ReadyBlue-stained RuBisCO is shown below as input loading control. **f,** Root length phenotypes of 8-d-old seedlings (n=30) shown as box plots (centre line, median; box, interquartile range IQR; whiskers, lowest/highest data point within l.5 IQR of the lower/upper quartile, raw data shown a gray dots). Multiple comparison of the different treatments vs. the indicated controls were performed using a Dunnett^91^ test as implemented in the R package multcomp^92^ (multiple comparison against *bril*: \**p* < 0.05; \*\**p* < 0.01; \*\*\**p* < 0.001; against Col-0 # *p* < 0.05; ## *p* < 0.01; ### *p* < 0.001).

Consistent with the BR-binding site’s structural plasticity, mutation of the conserved Thr646, Tyr597, or Lys601 from the island domain had no detectable effect on brassinolide binding (Fig. 5a-c). Substitution of Tyr599 or Tyr642 with alanine moderately reduced binding affinity (K_d_ ∼100 nM and ∼200 nM, respectively), and replacing the invariant Leu615 with phenylalanine or tyrosine likewise had little effect on BR binding (Fig. 5a-c). In contrast, mutation of Trp564 --- His caused a ∼100-fold drop in BR binding (K_d_ ∼2 µM) (Fig. 5a-c). Notably, mutating Met657 (Glu614 in BRL2, see Supplementary Fig. 3b) to either alanine or glutamate did not affect brassinolide binding (Fig. 5a-c). Consistently, a mild growth phenotype has been previously reported for the BRIl^M657E^ mutant^46^. These findings underscoring the presence of a conserved BR-binding site in BRIl/BRLl/BRL3 and in BRL2 (Fig. 4a). As previously shown^46^, the *bril-6*^45^ mutation (Gly644---Asp) strongly reduces brassinolide binding (Fig. 5a-c). In contrast, the neighboring Gly643---Glu mutation in the BRIl*^sudl^* gain-of-function allele^47^ stabilizes the island domain^36^ and enhances brassinolide binding by ∼13-fold over wild type (Fig. 5a-c). Similarly, mutating LRR core Phe681 to alanine increases binding by ∼7-fold. These results further highlight the BR-binding site’s structural plasticity and identify Trp564 from the LRR core as essential for ligand recognition.

To assess the functional impact of these mutations, we expressed mutant BRIl receptors under the native BRIl promoter in a *bril* null background (Fig. 5d). T3 lines with wild-type-like BRIl expression (Fig. 5e and Supplementary Fig. 4c) and proper plasma membrane localization (Supplementary Fig. 4d) were selected. High-affinity mutants (Tyr597---Phe, Leu615---Phe, Thr646---Ile) restored wild-type rosette and root growth phenotypes (Fig. 5d-f), and BR responses (Supplementary Fig. 4e,f and Extended Data Table 6). Intermediate phenotypes were observed in lines with Tyr597---Met, Leu615---Tyr, or Met657---Ala, consistent with their moderate in vitro binding affinities (Fig. 5a-c). The Gly644---Asp mutant displayed medium-to-strong defects, while Trp564---His, which showed ∼100-fold reduced binding, phenocopied the *bril* null at both seedling and adult stages (Fig. 5d-f).

Conversely, Gly643---Glu and Phe681---Ala conferred gain-of-function rosette phenotypes, matching their enhanced binding affinities compared to wild type (Fig. 5a-d). Three mutants (Tyr599---Ala, Lys601---Leu, Tyr642---Ala) displayed severe growth phenotypes despite retaining strong brassinolide binding in vitro (Fig. 5a-f, see below). Taken together, we observe a strong correlation between the in vitro ligand binding affinities and in planta growth phenotypes of BRIl mutant receptors, yielding novel loss-of-function (Trp564-His) and gain-of-function (Phe681-Ala) alleles.

### BR antagonists target the interaction of BR receptors with SERK co-receptors

We previously showed that the BR-binding site comprises the LRR core and island domain of BRIl, and the LRR ectodomain of a SERK co-receptor^26,36,28^ (Fig. 6a). SERKl Phe61 stacks with the A/B rings of brassinolide, while His62 forms a conserved hydrogen bond with the C_2_ hydroxyl group of the ligand^36^ (Fig. 6a). Mutating the corresponding Phe60 and His61 in SERK3/BAKl to alanine abolishes both co-receptor interaction in vitro and BR signaling in planta^28^.

**Fig. 6.**
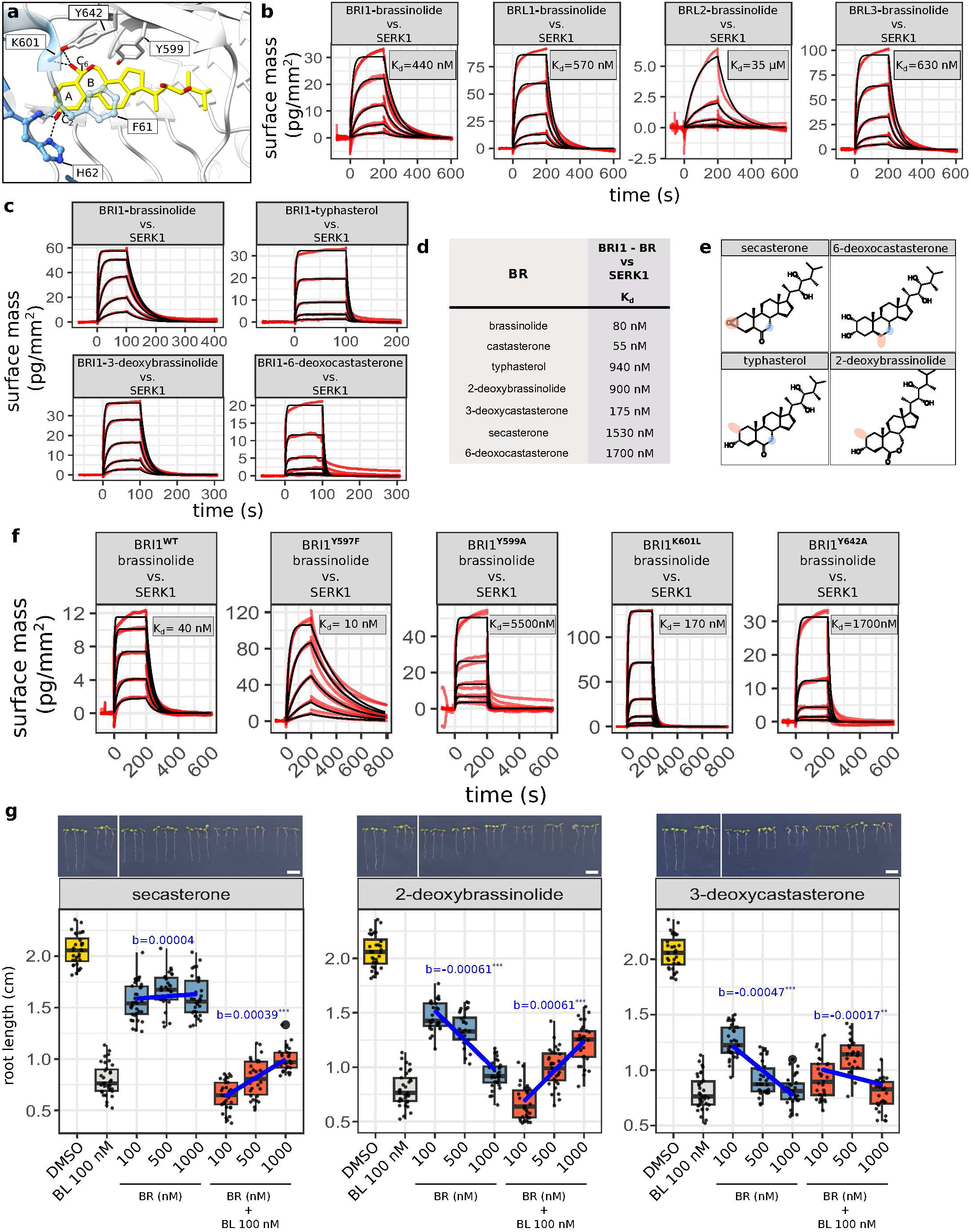
I BR C_2_ hydroxylation and C_6_ oxidation are critical for SERK co-receptor binding. **a,** Close-up view of the SERKl-bound (in blue) BRIl (in gray) - brassinolide (in bonds representation, in yellow) complex (PDB-ID: 000041sx). Receptor and co-receptor amino acids involved in BR- binding are highlighted as ball-and-sticks in gray and blue, respectively. Polar interactions between the brassinolide C_2_ hydroxyl group and His62, or the C_6_ oxygen with Tyr642 and Lys601 are shown as dashed lines (in black). **b,** GCI binding kinetics of amine-coupled and brassinolide-bound BRIl, BRLl, BRL2, and BRL3 ectodomains vs. SERKl. Shown are sensorgrams with red traces indicating experimental data and black lines representing global fits (n=l). Binding affinities (K_d_) are reported alongside. **c,** Representative GCI traces showing the interaction between SERKl and biotinylated, Avi-tagged BRIl ectodomain in the presence of either brassinolide, typhasterol, 3- deoxybrassinolide, or 6-deoxocastasterone. **d,** Table summaries of the receptor - BR - co-receptor interaction dissociation constants. **e,** Chemical structures of selected BRs tested in **c**,**d**. Structural modifications relative to brassinolide that reduce co-receptor binding are highlighted in red. Modifications that have little effect on complex formation are shown in blue. **f,** GCI binding kinetics of amine-coupled wild-type BRIl, BRIl^Y597F^, BRIl^Y599A^, BRIl^K601L^, and BRIl^Y642A^ ectodomains vs. SERKl in the presence of a high, saturating concentration of brassinolide. Shown are sensorgrams with red traces indicating experimental data and black lines representing global fits (n=2). Binding affinities (K_d_) are reported alongside. **g,** BR seedling root growth inhibition antagonist assay (n=30-35). Brassinolide (BL) at a fixed concentration of 100 nM was used in the different treatments, vs. 0.02% (v/v) DMSO as control (in yellow). The antagonist candidate compounds secasterone, 2-deoxybrassinolide and 3-deoxycastasterone were tested for their ability to inhibit seedling root growth by themselves (in blue), or to reverse BL-induced root growth inhibition (in orange). Shown are boxplots with centre lines (medians), box limits indicate the 25th and 75th percentiles, whiskers extend to the full data range. The relationship between antagonist concentration and root length were tested using a Tukey trend test^93,94^. The blue line represent the linear trend with slope b (negative values indicate agonistic behavior, positive values indicate antagonistic behavior) and the associated test p-value (\**p* < 0.05; \*\**p* < 0.01; \*\*\**p* < 0.001). Representative seedling images are shown above (scale bar = 0.5 cm). The DMSO and BL 100 nM controls are same in each panel, indicated by a white vertical line.

To test whether this activation mechanism is conserved across BR receptors, we immobilized BRIl, BRLl, BRL2, and BRL3 ectodomains on a GCI chip and examined SERKl binding in the presence of 100 nM brassinolide. BRLl and BRL3, like BRIl, bound SERKl with high affinity, while BRL2 displayed much weaker SERKl binding (K_d_ ∼35 µM), even in the presence of 2 µM brassinolide (Fig. 6b, Extended Data Fig. 5, Extended Data Table 7).

We next quantified the interaction of SERKl with BRIl in the presence of different BRs. We used an avi-tagged BRIl ectodomain to minimize steric hindrance between the GCI chip and the SERKl co-receptor (see Methods), and measured dissociation constants of ∼100 nM for SERKl association to either brassinolide- or castasterone-bound BRIl (Fig. 6c,d, Extended Data Fig. 5, Extended Data Table 7). However, SERKl binding was ∼15-fold weaker with typhasterol- vs. castasterone-bound BRIl (Fig. 6c,d), despite these ligands having similar receptor binding affinities (Fig. 2c,d). This reveals that the C_2_ hydroxyl coordinated by SERKl His62 is key for receptor - co- receptor complex formation (Fig. 6a,e). Similarly, 2-deoxybrassinolide reduced SERKl binding ∼10-fold, while 3-deoxycastasterone, lacking the C_3_ hydroxyl, still supported high-affinity SERKl binding (Fig. 6c-e). Secasterone - with a bulky 2,3-a-epoxy group - impaired complex formation, as did 6-deoxocastasterone, despite possessing both C_2_ and C_3_ hydroxyls (Fig. 6c-e), likely due to its distorted binding mode in the BR pocket (Fig. 3a,b).

Our data identify the BR C_2_ hydroxyl as essential for SERK recruitment. While C_6_ oxidation on the B-ring is not required for receptor binding, its coordination by BRIl island domain residues Tyr599, Tyr642, and Lys601 may help orient the ligand for optimal co-receptor binding (Fig. 6a,c- e). Indeed, SERKl binding is partially or severely compromised in the BRIl^Y599A^, BRIl^K601L^ and BRIl^Y642A^ mutants in the presence of high, saturating concentrations of brassinolide (Fig. 6f). This results rationalize the dwarf phenotypes of the BRIl Tyr599---Ala, Lys601---Leu and Tyr642---Ala mutants (Figs. 5d-f, 6a), despite their ability to bind brassinolide with high affinity (Fig. 5c).

As BRs that bind BRIl but block SERK association can act as receptor antagonists, we performed seedling root growth inhibition assays in the presence of 100 nM brassinolide and with different antagonist candidates from our biochemical screen (Fig. 6d). We found that secasterone applied on its own did not trigger root growth inhibition in a concentration-dependent manner (Fig. 6g, Extended Data Table 8). However, applying increasing concentrations of secasterone significantly reversed brassinolide-induced root growth inhibition (Fig. 6g). 2-deoxybrassinolide acted as a weak BR agonist when applied on its own, but had a significant, dose-dependent antagonistic effect on brassinolide-induced root growth inhibition (Fig. 6g). In contrast, 3- deoxycastasterone acted as a weak BR agonist both in the absence and presence of brassinolide in the assay (Fig. 6g). We conclude from these experiments that the BRs lacking a C_2_ but not C_3_ hydroxyl group behave as receptor antagonist (Fig. 6a,c-e,g). A bulky head group, such as in secasterone is also effective in blocking receptor activation in vitro and in planta (Fig. 6a,c-e,g). We speculate that the agonistic effect of 2-deoxybrassinolide may be caused by this metabolite entering the biosynthetic pathway, generating other, bioactive BRs (Fig. 6g and Fig. lb). Taken together, BRs that block BRIl - SERKl complex formation (Fig. 6c-e) in vitro without compromising isolated receptor binding (Fig. 2c,d), can partially reverse brassinolide-induced root growth inhibition in a dose-dependent manner (Fig. 6g), underscoring the importance of A-ring features in BR receptor activation^36,28^.

## Discussion

Genetic and metabolite studies have revealed a complex BR biosynthetic network in plants^2,l4^. In this study, we provide quantitative binding kinetics for 15 distinct BRs from this network, evaluated against the four BR receptors in Arabidopsis. Like other plant hormones^56,57^, BRs act as molecular glues promoting the association of receptor and co-receptor. Our work now defines the chemical features required for high affinity receptor binding and co-receptor recruitment: Hydroxylation at C_23_ and C_24_ stereochemistry in the BR aliphatic tail are critical for receptor recognition (Fig. 2b-d) and BR bioactivity (Fig. 3d and Fig. 7). Consistent with this, C_23_ hydroxylase mutants display severe dwarfism^9,51^. Lack of a C_6_ ketone is tolerated by the isolated receptors (Fig. 2b-d) but impairs proper ligand positioning and SERK co-receptor recruitment (Fig. 6c-f and Fig. 7). The presence of a B-ring lactone is not required for high affinity receptor binding (Fig. 2b-d and Fig. 3a,b), in agreement with the moderate growth phenotype of the *br6ox2* mutant^l3,51^. Notably, C_2_ hydroxylation is essential for SERK recognition^28,36^ (Fig. 6c-e) but not for receptor binding (Fig. 2b-d and Fig. 3a,b), providing a mechanistic basis for known BR antagonists^58,59^ (Fig. 7). In planta, 2-deoxybrassinolide and secasterone act as receptor antagonists (Fig. 6e,g), supporting the idea that these abundant^48^ biosynthetic intermediates may modulate BR signaling by competing with agonists for receptor binding (Fig. lb, Fig. 6g). This could prevent formation of active BR signaling complexes, in a similar way to the known BIR receptor pseudo- kinases^37,38^.

**Fig 7.**
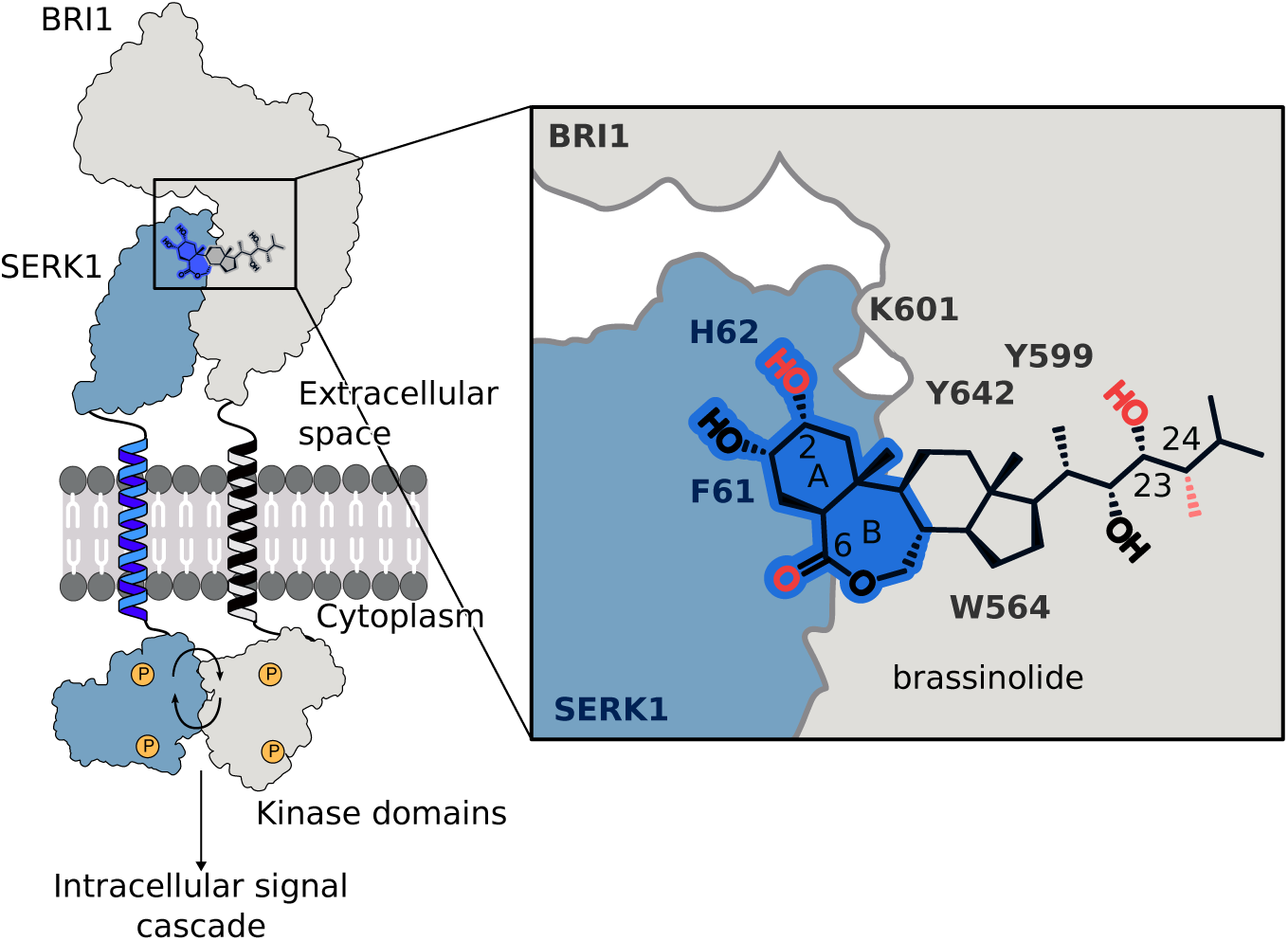
I Strong receptor binding, correct positioning within the BR binding site, and interaction with the co-receptor are essential for BR bioactivity. Schematic overview of brassinosteroid- induced receptor - co-receptor complex formation and activation. A BR ligand binds to the BRIl receptor ectodomain (in gray), creating a binding surface for a SERK co-receptor kinase (in blue). This brings the kinase domains of receptor and co-receptor in close proximity, leading to trans- phosphorylation and intracellular signaling. The BR-binding pocket is formed by the receptor (in gray) and co-receptor (in blue). Chemical features of the BR ligand (in black) involved in co- receptor binding are highlighted in dark-blue, key carbon atoms critical for receptor or co-receptor interaction are numbered, chemical features required for receptor and/or co-receptor binding are shown in red. The approximate positions of residues critically involved in BR binding, positioning and co-receptor binding are indicated.

Importantly, our quantitative BR binding assay identifies BRL2 as a functional BR receptor with a distinct ligand binding spectrum (Fig. 2c,d, Fig. 4b,c). BRL2 shares a conserved BR binding mechanism (Supplementary Fig. 3b) and weakly interacts with SERK co-receptors (Fig. 6b). The gain-of-function phenotype of oBIR3-iBRL2 receptor chimera (Fig. 4e) further supports the notion that BRL2 shares the SERK-based receptor activation mechanism^54^ with BRIl^28,35,36^, BRLl and BRL3 (Fig. 6b). We speculate that the inability of BRL2 to replace BRIl in *bril* mutants^29,33^ is caused by its distinct BR ligand binding spectrum (Fig. 4b,c), while its receptor activation mechanism and cytoplasmic signaling pathway appear to be conserved^60,61^. Taken together, BRL2, which emerged early on in land plant evolution, may represents an ancestral BR receptor^62^.

Mutational analysis of the BR-binding pocket in BRIl reveals that the receptor can tolerate various missense mutations (Fig. 5 and Fig. 7), in agreement with previous mutational studies^46,63^. We identified single point mutation causing a strong BR loss-of-function phenotype (Trp564His), two residues (Tyr599 and Tyr642) critically involved in the proper positioning of the ligand in the BR binding pocket, and a novel gain-of-function allele (Phe681Ala) (Fig. 5d-f and Fig. 7). The structural plasticity of the hormone binding pocket supports the receptor’s capacity to recognize chemically diverse BRs, while providing sufficient selectivity to discriminate between BRs, other phytosterols and triterpenoids. Notably, the BR-binding site is conserved among the four Arabidopsis receptors and their orthologs in crop species, implying that BR receptors in crops likely share a similar BR ligand binding spectrum (Supplementary Fig. 4a).

Based on binding profiles (Fig. 2c,d, Fig. 6c,d), bioactivity data, and in vivo BR levels^ll,48,50^ (Fig. 3d), we propose that brassinolide and castasterone (C_28_), 28-norbrassinolide (C_27_), and possibly 28-homocastasterone (C_29_) may act as physiological BR ligands in Arabidopsis. Feedback regulation of BR biosynthesis likely fine-tunes these BR levels (Fig. 3e). Given that BR biosynthetic enzymes have specific expression domains^51,66^, different BR ligands may be perceived by the broadly expressed BRIl receptor in specific tissues and organs. Such a broad ligand spectrum has been previously reported for other small molecule^64^ and peptide^65^ hormone receptors in plants. We would like to note that our in vitro binding assays with the isolated LRR domains probably underestimate the actual BR binding affinities of intact BRIl - SERK receptor complexes embedded in the plasma membrane, as plants can respond to BRs applied externally at picomolar to low nanomolar concentrations^44,51^.

Taken together, distinct chemical features in BRs required for (l) high affinity receptor binding, (2) proper ligand orientation in the receptor hormone binding pocket, and (3) high affinity co-receptor binding (Fig. 7) have now been defined and can be exploited for rational BR agonist and antagonist design.

## Online material and methods

### Protein expression and purification

The ectodomains from AtBRIl (https://www.uniprot.org UniProt-ID O22476, residues l- 788), AtBRLl (Q9ZWC8, l-770), AtBRL2 (Q9ZPS9, l-752), AtBRL3 (Q9LJF3, l-770) and AtSERKl (Q94AG2, 23-270) ectodomains were amplified from *Arabidopsis thaliana* cDNA (ecotype Col-0). AtBRIl, AtBRLl, AtBRL2 and AtBRL3 were cloned in vector pBB2 providing tobacco etch virus (TEV) protease cleavable C-terminal Twin-StrepII and 9xHis affinity tags. AtSERKl was cloned in the same pBB2 vector containing an N-terminal azurocidin signal peptide^67,28^. For grating-coupled interferometry assays, AtBRIl was also cloned in vector pBB2 containing a C-terminal Avitag followed by TEV cleavable C-terminal Twin-StrepII and 10xHis affinity tags.

All proteins were expressed in *Trichoplusia ni* (Tnao cells, Invitrogen) using the baculovirus system^68^. Tnao cells in SF900 medium (Gibco) were grown to a density of 2.0 × 10^6^ cells/mL and subsequently infected with 10 mL of P2 virus. Cells were then incubated for 24 h at 28 °C and for an additional 48 h at 21 °C while shaking at 110 rpm. Cells were then pelleted at 4’000 x g for 20 min and the supernatant containing the secreted protein was filtrated using 0.45 µm filters (Durapore Merck). The cleared medium was then loaded onto an Ni^2+^ metal affinity column (HisTrap Excel 5 mL, Cytiva) equilibrated in buffer A (50 mM KPi pH 7.8, 250mM NaCl, l mM - mercaptoethanol). The column was washed with 25 mL of buffer A and subsequently eluted onto a Strep-Tactin XT 4flow 5mL column (IBA) in buffer A supplemented with 250 mM imidazole (pH 8.0). The Strep-Tactin affinity column was further washed with 25 mL buffer (20 mM Tris pH 8.0, 250 mM NaCl and l mM EDTA), and the protein was eluted from the Strep-Tactin column with buffer C (10 mM Tris pH 8.0, 150 mM NaCl, l mM EDTA, 50 mM biotin, lxBTX, IBA). For protein crystallization, the affinity purified proteins were incubated overnight at 4 °C with TEV protease using a molar receptor:protease ration of ∼10:l. The tag free protein was separated from the tag and from TEV protease by a second Ni^2+^ affinity chromatography step. Finally, the untagged protein was applied to a HiLoad Superdex 200 16/200 pg column (Cytiva) equilibrated in 20 mM sodium citrate pH 5.0, 150 mM NaCl. Fractions containing the purified protein were concentrated and stored at -80 °C.

Receptor biotinylation for grating-coupled interferometry was achieved by incubating Avi- tagged BRIl with His-tagged BirA^69^. Briefly, BRIl at a final concentration of 20 µM was incubated for l h at 30 °C with BirA, biotin, ATP and MgCl_2_ at final concentrations of 85 µM, 150 µM, 2 mM and 5 mM respectively. The BirA enzyme was subsequently removed using a Ni^2+^ affinity chromatography step. Biotinylated BRIl was concentrated and loaded onto a HiLoad Superdex 200 16/200 pg column (Cytiva) equilibrated in 20 mM sodium citrate pH 5.0, 150 mM NaCl.

### Chemical synthesis and analysis of BRs

The chemical synthesis of typhasterol, 3-deoxycastasterone and 2-deoxybrassinolide are described in Supplementary Note l.

### Grating Coupled Interferometry (GCI)

GCI experiments were performed using the Creoptix WAVE system (Malvern Panalytical). Binding of BRs to the isolated BRIl, BRLl, BRL2 and AtBRL3 ectodomains was measured using 4PCH WAVEchips. Borate buffer (100 mM sodium borate, pH 9.0, l M NaCl, Xantec) injections were used to condition the chips. For the interaction screening between BRIl, BRLl, BRL2, BRL3 or BRIl^mut^ with BRs, the purified ectodomains were immobilized using standard amine-coupling. Chips were activated using a l:l mix of 400 mM N-(3-dimethylaminopropyl)-N’-ethylcarbodiimide hydrochloride and 100 mM N-hydroxysuccinimide (Xantec). Protein Ligands (in 10 mM sodium acetate, pH 5.0) were injected to the desired chip surface at the concentration indicated in Extended Data Tables l, 4 and 5. In order to minimize non-specific binding of analytes, a second injection of BSA (0.5% [w/v] in 10 mM sodium acetate, pH 5.0) was used to passivate the surface. Finally, the chip was then quenched by injecting l M ethanolamine. All kinetic analyses were performed at room temperature in citrate buffer (20 mM sodium citrate pH 5.0, 150 mM NaCl) supplemented with 0.05 % (v/v) Tween and 0.025 % (v/v) dimethyl sulfoxide to reduce unspecific binding of the respective BR analyte. Analyte concentrations were varied in l:3 dilution series (compare Extended Data Tables l, 4, 5). All experiments were done in duplicates, blank injections were used for double referencing and DMSO calibration was performed for bulk correction.

Binding of the co-receptor SERKl to brassinolide-bound wild-type and mutant receptors was done by amine-coupling the BRIl, BRLl, BRL2 and BRL3 ectodomains onto 4PCP WAVEchips and using the purified SERKl ectodomain as analyte (Extended Data Table 7) in 20 mM sodium citrate pH 5.0, 150 mM NaCl) supplemented with 0.05 % (v/v) Tween and 0.025 % (v/v) dimethyl sulfoxide and brassinolide to a final concentration of l µM. Binding of SERKl to BRIl in the presence of different BRs was performed by immobilizing Avi-tagged and biotinylated BRIl by neutravidin capturing. After chip activation, neutravidin (100 µg/mL in 10 mM sodium acetate, pH 5.0) was injected. To passivate and quench the chip surface, a further injection of 0.5% (w/v) BSA followed by an injection of l M ethanolamine were performed, respectively. Biotinylated BRIl (100 µg/mL in citrate buffer) was then injected to the desired surface concentration (Extended Data Table 7), followed by an injection of biotinylated 0.5% (w/v) BSA in order to passivate the surface. All the binding assays between BRIl and SERKl were done at room temperature in citrate buffer supplemented with 0.05% Tween and the respective BR at a final concentration of of l µM. SERKl injections were done in l:3 dilution series (Extended Data Table 7).

Data analysis was performed with the Creoptix WAVEcontrol software (version: 4.7.2) and data were fitted using one to one binding models or mass transport model, using bulk correction. Plotting and data representation were performed using R (https://www.r-project.org, version: 4.l.2).

### Crystallization and data collection

BRL3 (in 20 mM citric acid pH 5.0, 250 mM NaCl) at a concentration of ∼l mg/mL was incubated with the respective BR (10 mM stock solution in 100 % [v/v] dimethyl sulfoxide [DMSO]) to a final concentration of 50 µM for 4 h at room temperature and then concentrated to ∼6-7 mg/mL using a Amicon Ultra concentrator (Merck Millipore) with a molecular weight cut-off of 30 kDa. Triclinic crystals developed in hanging drops composed of l.0 µL of protein solution and l.0 µL of crystallization buffer (23-25 % [w/v] PEG 3,350, 0.2 M Li_2_SO_4_, 0.l M citric acid pH 4.0) suspended over l.0 mL of the latter as reservoir solution. Crystals were cryoprotected by serial transfer into crystallization buffer containing ethylene glycol to a final concentration of 25 % (v/v), and snap frozen in liquid N_2_. BRIl (in 20 mM citric acid pH 5.0, 250 mM NaCl) - BR (at a final concentration of 50 µM) complex crystals developed in the same crystallization condition. Crystals of apo BRL2 were obtained using micro-seeding protocols in hanging drops containing l.0 µL of protein solution (BRL2 at 6 mg/mL in 20 mM citric acid pH 5.0, 250 mM NaCl) and l.0 µL of crystallization buffer (23 % [w/v] PEG 3,350, 0.2 M MgC12 • 6H_2_O, 0.l M Tris pH 9.0) over l mL of reservoir solution. Tetragonal crystals were cryoprotected by serial transfer in crystallization buffer supplemented with 20% (v/v) glycerol. All datasets were collected at beam line X06DA of the Swiss Light Source, Villigen, Switzerland (Supplementary Table l). Data processing and scaling was done with XDS^70^ (version: June 2024).

### Crystallographic structure solution and refinement

The structures of the BRL3 - brassinolide complex and of apo BRL2 were solved by molecular replacement in PHASER^71^, with BRIl ectodomain structure^26^ (PDB-ID pdb_00003rj0) as search model. For BRL3, the solution revealed a dimer in the asymmetric unit, closely resembling the previously reported BRLl dimer^31^. The BRL2 structure was resolved at 2.6 A resolution in space group *P* 4_l_ 2_l_ 2, containing a single molecule per asymmetric unit (Supplementary Table l). Crystals of the BRIl-BR complex shared the same crystal form as the previously described apo BRIl^26^. Restraints for the BR ligand were generated from SMILES strings using phenix.elbow^72^. All structures were completed through iterative cycles of manual model building in COOT^73^ and restrained TLS refinement in phenix.refine^74^. Analysis of the refined structures with phenix.molprobity^75^ revealed good to excellent stereochemistry. Omit maps of the different BR ligands were generated with phenix.polder^76^. Data collection and refinement statistics are provided in Supplementary Table l. Structural figures were prepared with ChimeraX^77^, chemical structures were drawn with PubChem sketcher^78^.

### Plant material

*Arabidopsis thaliana* ecotype Col-0, T-DNA insertion mutants *bsk3-l* (SALK_096500C)^79^, *bril* null (GABI_134E10)^47^ and *brl2-l* (SALK_016024)^55^ were obtained from the Nottingham Arabidopsis Stock Center, alongside the previously reported *BRIlprom*::BRIl-mCitrine / *bril* and *BRIlprom*::BRIl*^sudl^*-mCitrine / *bril*^36^ and *BRIlprom*::oBIR3-iBRIl-mCitrine / *bril*^54^ transgenic lines. Genotyping primers for the mutants are listed in Supplementary Table 2.

### Molecular cloning and generation of transgenic lines

To generate BRL2 expression reporter lines, a 3.6 kb fragment upstream of the BRL2 ORF was cloned into the entry vector pUC57- L4_EcoRV-XbaI-BamHI_Rl via Gibson assembly to produce L4-*BRL2prom*-Rl^80^. Multi-site Gateway cloning (Thermo Fisher Scientific) with Gateway LR reaction (11791020; Thermo Fisher Scientific) was used to assemble the following fragments L4-BRL2prom-Rl, Ll-NLS-3xmVenus-L2^81^ and R2-tHSP18.2-L3^82^ into the destination vector pB7m34GW^83^, to generate *BRL2prom*::NLS-3xmVenus. The BRL2 chimera (*BRL2prom*::oBIR3- iBRL2-mCitrine) were generated using a construct in which the N-terminal part of AtBIR3 (residues l-246, containing the native signal peptide, extracellular domain, and transmembrane helix) and the C-terminal region of BRL2 (residues 779-1143, comprising the juxtamembrane and kinase domains) were joined using two-fragment Gibson assembly, as previously described^54^. For the BRIl complementary assays, the BRIl coding sequence lacking a stop codon was cloned into entry vector pDONR221^36^ using the Gateway BP reaction (11789020; Thermo Fisher Scientific). The single point mutations were introduced by the site-directed mutagenesis by overlap extension^84^ on the entry vector. A 1690 bp upstream fragment of BRIl (*BRIlprom*) and a mCitrine coding sequence with the stop codon was cloned into pDONR P4-PlR and pDONR P2R-P3, respectively, as previously reported^54^. The final binary vectors *BRIlprom*::BRIlmu-mCitrine were generated by the multi-site Gateway technology (Thermo Fisher Scientific) with destination vector pB7m34GW^83^. Cloning primers are list in Supplementary Table 3.

The *brl2-2* allele was generated using a CRISPR/Cas9^85^ system^86^. Two 20 bp sgRNA target sequences, designed using CRISPR-P 2.0^87^, were cloned into shuttle vectors pEF004/005 via mutagenesis PCR^88^. These cassettes were subsequently assembled into the pBay-Bar-CRmCherry vector using Gibson assembly and the final construct was transformed into Col-0 wild-type plants using the floral dip method^89^. Primary transformants (Tl) were selected on ^l/2^MS medium containing 15 mg/L phosphinothricin; Cas9-free mutant lines (T2/T3) were identified by screening for loss of the mCherry marker (Leica Stereo Microscope MSV269, Wetzlar, Germany).

All constructs were transferred into *Agrobacterium tumefaciens* strain GV3101 by electroporation and transgenic lines were generated by floral dip^89^. Transformants were selected on half-strength Murashige and Skoog (^l/2^MS) media containing l% (w/v) sucrose, 0.8% (w/v) agar and 15 mg/L phosphinothricin.

### BR root growth inhibition and competition assays

For BR-mediated root growth inhibition assays^43,44^, surface sterilized seeds were stratified at 4°C for 2 d and plated on ^l/2^MS media plates containing l% (w/v) sucrose and 0.8% (w/v) agar, supplemented with 100 nM of the indicated BR from 10 mM stock solutions prepared in 100 % (v/v) DMSO (Carl ROTH) or, with 0.01% (v/v) DMSO as control. The previously reported *bsk3-l* T-DNA insertion mutant (SALK_096500C)^79^ was used as negative control. For the BR antagonist competition assays, 100 nM brassinolide and 100, 500 or 1000 nM of antagonist candidate were added to ^l/2^MS plates containing l% (w/v) sucrose and 0.8% (w/v) agar. DMSO to a final concentration of 0.02% (v/v) was used as control treatment. In all cases, plates were moved to a Percival (SE-41AR3; CLF PlantClimatics, settings: 70% humidity, 16 h light period at 22°C with 250 µmol m^-2^ s^-l^ light intensity and 8 h dark period at 18°C), vertically placed for 7 d. The plates were scanned using a flat-bed scanner (Epson perfection V600 Photo, Epson). The primary root length of 30-35 individual seedlings was then determined using Fiji^90^.

### Statistical analysis

Multiple comparisons of different treatments vs. control(s) in one-way layouts were performed using a two-sided Dunnett^91^ test as implemented in the R package multcomp^92^. The relationship between root length and dissociation constant were tested using a Tukey trend test^93,94^, i.e. a multiple regression model approach using arithmetic, ordinal or logarithmic scores. We chose the linear regression model and represented the corresponding slope b (positive slopes were interpreted as antagonistic; negative slopes as agonistic) and associated p-values. Data were plotted in R^95^ (version 4.2.2), the associated raw data can be found in Extended Data Tables 2, 6 and 8.

### BR in vivo quantification

The extraction and quantification of endogenous BRs were carried out as previously described^96^. In brief, five biological replicates, each consisting of approximately 10 mg fresh weight (FW) of plant material, were extracted using ice-cold 60% (v/v) acetonitrile. To each sample, 25 pmol of deuterium-labelled BR internal standards (Olchemim LtD, Olomouc, Czech Republic) were added. After a 12h incubation at 4°C, samples were centrifuged, and the resulting supernatants were purified using 50 mg Discovery DPA-6S solid-phase extraction cartridges (Supelco, Bellefonte, PA, USA). The eluates were then evaporated to dryness and reconstituted in 40 µL of methanol. BR quantification was performed using ultra-high-performance liquid chromatography coupled with tandem mass spectrometry (UHPLC-MS/MS), employing the ACQUITY UPLC I-Class System (Waters, Milford, MA, USA) and the Xevo TQ-S triple quadrupole mass spectrometer (Waters MS Technologies, Manchester, UK), using the previously reported UHPLC-MS/MS parameters^96,97^.

### RNA-seq experiments

10-d-old Arabidopsis plants grown on ^l/2^MS media plates containing l% (w/v) sucrose and 0.8% (w/v) agar were harvested. 15 plants were pooled per individual biological replicate and all samples were ground. RNA was extracted using a RNeasy Plant Mini Kit (74904; Qiagen) and total RNA concentration was estimated using a Qubit™ 4 Fluorometer (Q33238; Thermo Fisher Scientific) and a 2100 Bioanalyzer system (Agilent Technologies). Library preparation and sequencing (100 bp single-end with Novaseq 6000 sequencing system; Illumina) was performed at the iGE3 Genomic Platform at the Faculty of Medicine, University of Geneva,, Switzerland (https://ige3.genomics.unige.ch/). Data analysis was conducted within the Galaxy EU platform (https://usegalaxy.eu/) using the following tools: quality control and adaptor trimming by MultiQC^98^, reads mapping to the reference genome (Arabidopsis TAIR10 reference genome from TAIR; https://www.arabidopsis.org/) by HISAT2^99^, transcript assembly and FPKM estimates by Cufflinks^100^, reads counting by FeatureCounts^101^ and differentially gene expression analysis by DESeq2^102^. Visualization was done using SRplot^103^ (https://www.bioinformatics.com.cn/).

### Confocal microscopy

To detect BRL2 promoter activity with *BRL2prom*::NLS-3xmVenus (in T2 generation) in the early vasculature of embryos, SCRI Renaissance 2200 staining was performed according to the protocol^86,104^. The embryos were dissected from the siliques in the flowering 5-week-old plants under a Leica S6D stereo microscope (Leica S6D, Wetzlar, Germany) and incubated in an SR2200 solution containing 0.l% (v/v) SR2200, l% (v/v) DMSO, and 0.5% (v/v) Triton X-100, 5% (v/v) glycerol, and 4% (v/v) paraformaldehyde in phosphate-buffered saline buffer (pH 8.0) for five min. Then, the embryos were washed once with H_2_O and mounted in 5% (v/v) glycerol for imaging. The cell wall signal stained by SR2200 was obtained using a 405 nm laser and a detection wavelength ranging from 415 to 475 nm. To image roots, propidium iodide (PI) staining was used to label the cell wall. The primary root of 10-d-old seedlings previously grown on ^l/2^MS plates containing l% (w/v) sucrose and 0.8% (w/v) agar was incubated in a 10 mg/L PI solution for 10 minutes. Then, it was washed three times with water and incubated in a 5% (v/v) glycerol solution as the imaging buffer. PI fluorescence was detected from 571 nm to 656 nm upon a 561 nm wavelength laser excitation. The mVenus signal was detected using a laser with a 514 nm wavelength laser for excitation and detectors with an emission range between 517 and 570 nm. To detect the mCitrine signal in the roots of T2 generation *BRIlprom*::BRIl^mut^-mCitrine complementation lines, primary root tips from 10-d-old seedlings growing on ^l/2^MS plates containing l% (w/v) sucrose and 0.8% (w/v) agar were sampled. The mCitrine signal was detected by a laser with 514 nm wavelength for excitation, detectors recorded emission between 517 nm and 570 nm. All imaging was performed with a LSM 780 NLO microscope (Zeiss) at the Photonic Bioimaging Center, University of Geneva, Switzerland and data were analyzed using ZEN 2.0 blue edition and Fiji^90^.

### Analysis of leaf vein patterning

Leaf vein patterns were analyzed as described^55^. Cotyledons from 2-week-old seedlings grown on ^l/2^MS plates containing l% (w/v) sucrose and 0.8% (w/v) agar were dissected and cleared in 95% ethanol, hydrated in an ethanol dilution series to water, and mounted on slides in 10% (v/v) glycerol for imaging on a Nikon binocular microscope (Nikon SMZ18).

### Hypocotyl growth assay

Surface-sterilized Col-0 wild-type and *brl2-l* mutant^55^ seeds were cold-stratified at 4 °C for 48 h and sown on ^l/2^MS medium containing 0.8% (w/v) agar. The growth medium was supplemented with either: l µM brassinazole (BRZ)^105^ prepared from a 10 mM stock solution in 100% DMSO, or 0.l% (v/v) DMSO as control. After a l h light treatment to promote germination, the plates were wrapped in aluminium foil and maintained in complete darkness at 22 °C for 5 d. Seedling where then scanned using a flat-bed scanner (Epson Perfection V600 Photo scanner, Epson).

### BESl and BRIl-mCitrine immunoblotting

Arabidopsis seeds (Col-0 wild type, *bril*^47^, *brl2-l*^55^ and T3 generation *BRIlprom*::BRIl^mut^- mCitrine complementation lines) were surface-sterilized and grown on ^l/2^MS plates containing l% (w/v) sucrose and 0.8% (w/v) agar for 10 or 14 d, respectively. Around 250 mg of tissue samples were collected for each line in l.5 ml tubes containing glass beads and immediately frozen with liquid nitrogen. The samples were ground with Silamat S6 (Ivolar Vivodent) and then mixed with extraction buffer (50 mM Tris-HCl pH 7.5, 150 mM NaCl, l mM EDTA, l% [v/v] Triton X-100, lx cOmplete EDTA-free protease inhibitor cocktail [11873580001; Roche], lx plant protease inhibitor cocktail [P9599; Sigma-Aldrich] and lx PhosStop [4906845001; Roche]). After protein solubilization for 30 min on ice, cell debris was removed by centrifugation (14,000 xg, 2x 15 min, 4°C) and the supernatant of each sample was transferred to a fresh tube. Protein concentration determination with Pierce BCA Protein Assay Kit (23225; Thermo Fisher Scientific). For BRIl- mCitrine fusion protein detection, an anti-GFP antibody coupled to horseradish peroxidase (anti- GFP-HRP, 130-091-833; Miltenyi Biotec) at a dilution of l:5,000 was applied. Alternatively, BRIl was detected using a primary BRIl antibody^106^ (l:5000 dilution) and a secondary Anti-Rabbit IgG- Peroxidase antibody produced in goat (l:10000 dilution, A6154; Sigma-Aldrich). BESl dephosphorylation levels were estimated using a primary anti-BESl antibody^41^ at a dilution of l:2000 and a secondary Anti-Rabbit IgG-Peroxidase antibody produced in goat (l:10000 dilution, A6154; Sigma-Aldrich). SuperSignal West Pico PLUS Chemiluminescent Substrate (34577; Thermo Fisher Scientific) was used for chemiluminescence detection on an Amersham Imager 680 (GE). ReadyBlue Protein Gel Stain (RSB-lL; Sigma-Aldrich) was used for the total protein input detection.

### Analytical size-exclusion chromatography

The oligomeric state of recombinantly expressed and purified wild-type and point-mutant BRIl ectodomains was characterized using analytical size-exclusion chromatography. 500 µL sample at l.0 mg/mL was loaded onto a Superdex 200 Increase 10/300 GL column (Cytiva) pre- equilibrated in 20 mM citrate buffer (pH 5.0), 150 mM NaCl. Protein elution at a flow rate of 0.7 mL/min was monitored by UV absorption at 280 nm.

### Quantitative real-time PCR

Expression of the BR marker genes CPD and DWF4 was estimated by qRT-PCR, using ∼ 30 individual 8-d-old seedlings per each T3 generation transgenic line. Samples were homogenized by Silamat S6 (Ivolar Vivodent) and RNA was extracted using a RNeasy Plant Mini Kit (74904; Qiagen). Subsequently, cDNA synthesis was performed with the SuperScript™ II Reverse Transcriptase (18064014; Invitrogen), following the manufacturer’s protocol. Quantitative Real- time PCR was conducted on a QuantStudio 5 thermo-cycler (Applied Biosystems) using SYBR™ Select Master Mix for CFX (4472942; Applied Biosystems). The relative expression (n=3) was quantified with the 2 -l)l)Ct method. ACT2 was used as the internal reference gene. qPCR Primers are included in Supplementary Table 4.

## Data availability

Raw RNA-seq reads have been deposited with the NCBI sequence read archive (SRA) with identifier PRJNA1299306 (https://www.ncbi.nlm.nih.gov/sra/PRJNA1299306). BRIl, BRL2 and BRL3 complex structures have been deposited with the Protein Data Bank (http://www.rcsb.org) with accession numbers pdb_00009S80 (BRL3 - brassinolide), pdb_00009S87 (BRIl - castasterone), pdb_00009S8S (BRIl - typhasterol), pdb_00009S8V (BRIl - 24-epibrassinolide), pdb_00009S8Z (BRIl - 28-homobrassinolide), pdb_00009S90 (BRL3 - castasterone), pdb_00009S96 (BRL3 - typhasterol), pdb_00009S9A (BRL3 - 6-deoxocastasterone), pdb_00009S9C (apo BRL2).

## Acknowledgments

We thank Jenny Russinova for providing us with the BESl antibody, Danuse Tarkowska for help with UHPLC-MS/MS analyses, Adam Pribylka for the measurement of HRMS spectra, Samuele Meschini for help with the root growth inhibition assays and Philippe Rieu, Christian Hardtke, Roman Ulm and Dorothea Fiedler for providing feedback on the manuscript. We acknowledge the Paul Scherrer Institute, Villigen, Switzerland for provision of synchrotron radiation beamtime at beamline X06DA of the SLS. This work was supported by Grant 310030_205201 from the Swiss National Science Foundation (to M.H.) and IGA grant IGA_PrF_2025_019 from Palacky University (to M.S.).

## Author contributions

A.C. and M.H. conceived the project. A.C expressed and purified proteins, performed GCI assays and crystallized proteins. H.C. generated transgenic lines with the help of L.B., performed plant phenotyping and BR-response assays, confocal microscopy, transcriptomics and western blotting. M.K. and K.F. synthesized BRs. U.H. crystallized the BRL3 - brassinolide complex. J.O. quantified endogenous BR levels. A.C., U.H. and M.H. solved and refined structures. A.C., H.C., M.K., J.O., L.H., M.S. and M.H. analyzed data. A.C. and M.H. wrote the manuscript, with input from the other authors. M.H. and M.S. acquired funding and M.H. supervised the project.

---

**Supplementary Fig. 1.**
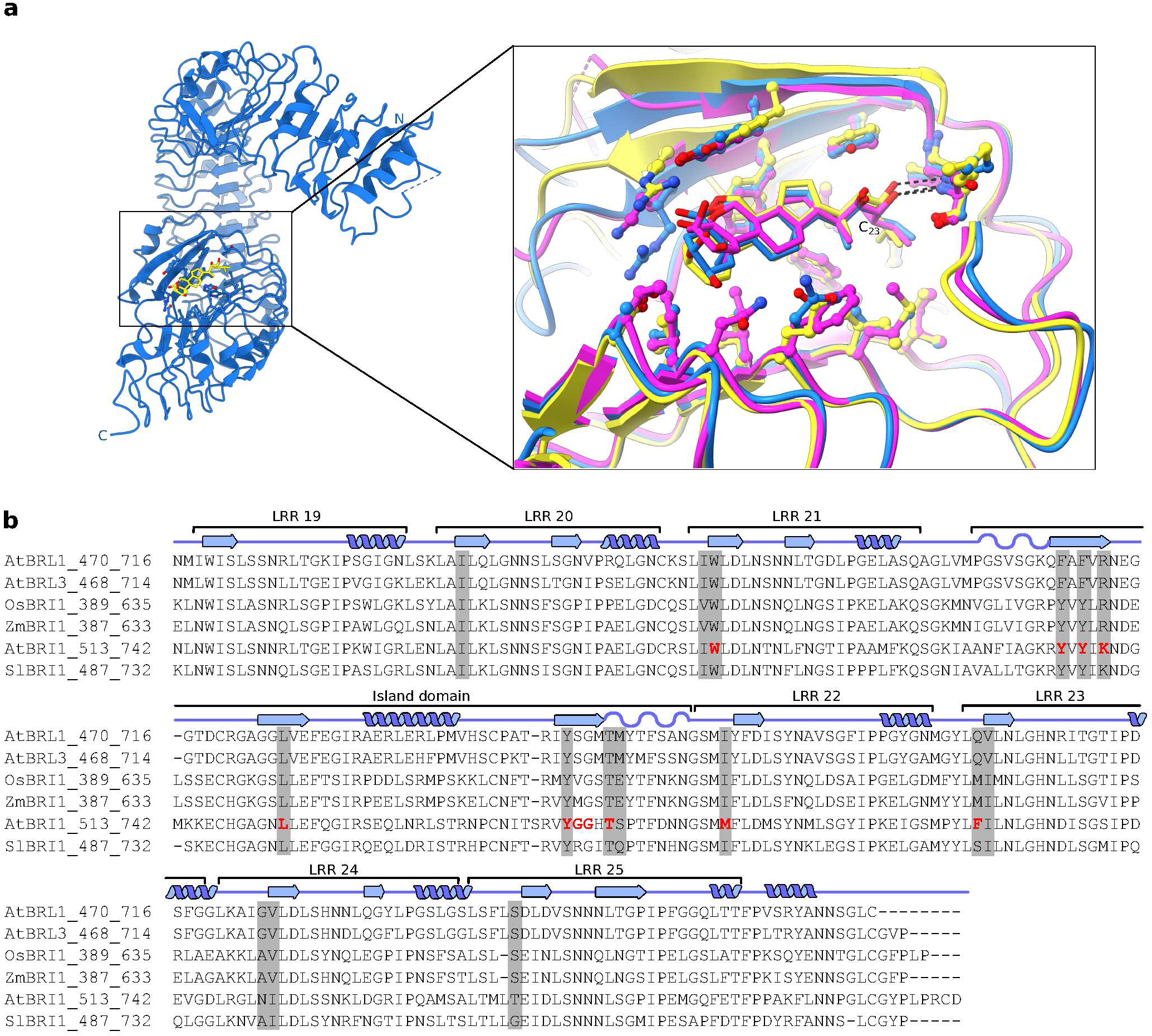
I BRil, BRLl and BRL3 share a structurally conserved BR-binding pocket. **a,** Right: Ribbon diagram of the BRL3 ectodomain bound to brassinolide (yellow, bonds representation). Left: Close-up of a structural superposition of the BR-binding pockets of BRL3 (blue), BRIl (purple; r.m.s.d. ∼ 0.7 A over 172 C_a_ atoms), and BRLl (yellow; r.m.s.d. ∼ 0.6 A over 166 C_a_ atoms). Key residues involved in brassinolide coordination are shown in ball-and-stick representation. The conserved hydrogen bond between the C_23_ hydroxyl group of brassinolide and the island domain backbone is indicated with a black dotted line. **b,** Multiple sequence alignment of BR-binding pocket regions from *Arabidopsis thaliana* BRIl, BRLl, and BRL3, along with orthologs from rice (OsBRIl, https://www.uniprot.org/ UniProt-ID Q942F3), maize (ZmBRIl, A0A3L6FIW6), and tomato (SlBRIl, F2XYF6), generated using MUSCLE^107^. Secondary structure elements based on AtBRIl and calculated with DSSP^108^ are shown above the alignment. BR- contacting residues (compare Fig. 2b) are highlighted in gray; residues mutated in this study (compare Fig. 4a) are shown in red.

**Supplementary Fig 2.**
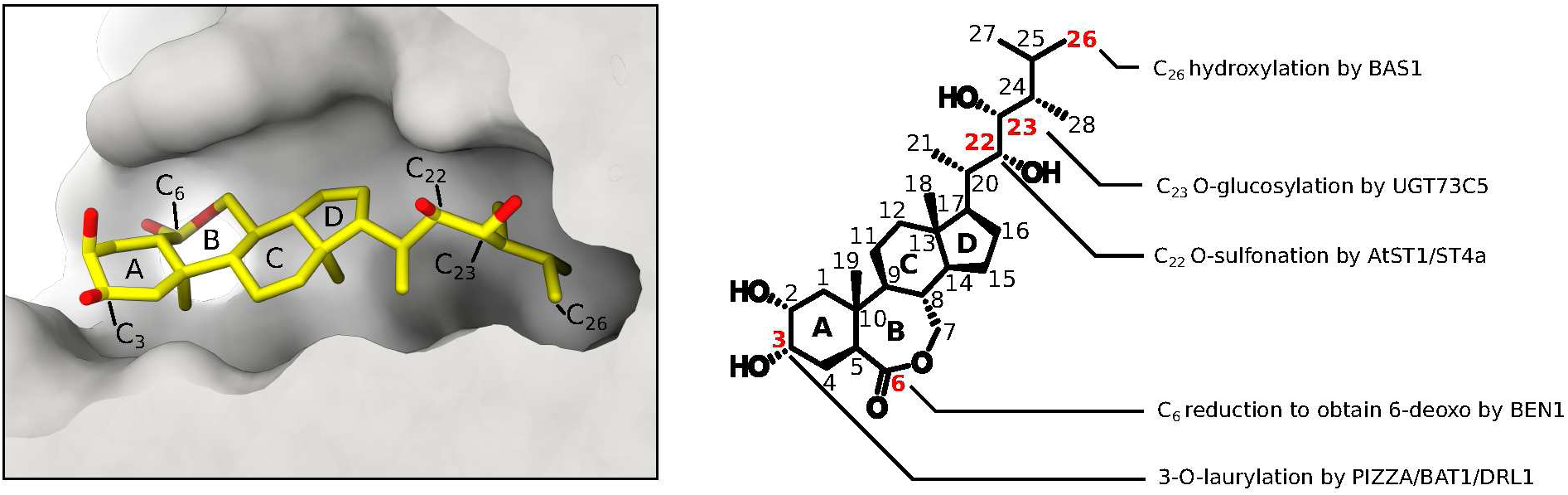
I BR catabolic enzyme modify BRs at positions critical for receptor and co-receptor binding. Left panel: Molecular surface view of the BRIl hormone binding pocket (in gray) occupied by brassinolide (in bonds representation, in yellow; PDB-ID: pdb_00003rj0). The A, B, C, and D rings that make up the hormone’s steroid core are labeled and carbon atoms targeted for catabolic modifications are highlighted. Right panel: Chemical structure of brassinolide with carbon numbering. Red, bold numbers indicate sites of BR catabolic reactions: C_26_ hydroxylation by BASl^l7^, C_23_ O-glucosylation by UGT73C5^l8^, C_22_ O-sulfonation by AtSTl/ST4a^l9^, C_6_ reduction by BENl^109^ and C_3_ O-laurylation by PIZZA/BATl/DRLl^110-112^.

**Supplementary Fig. 3.**
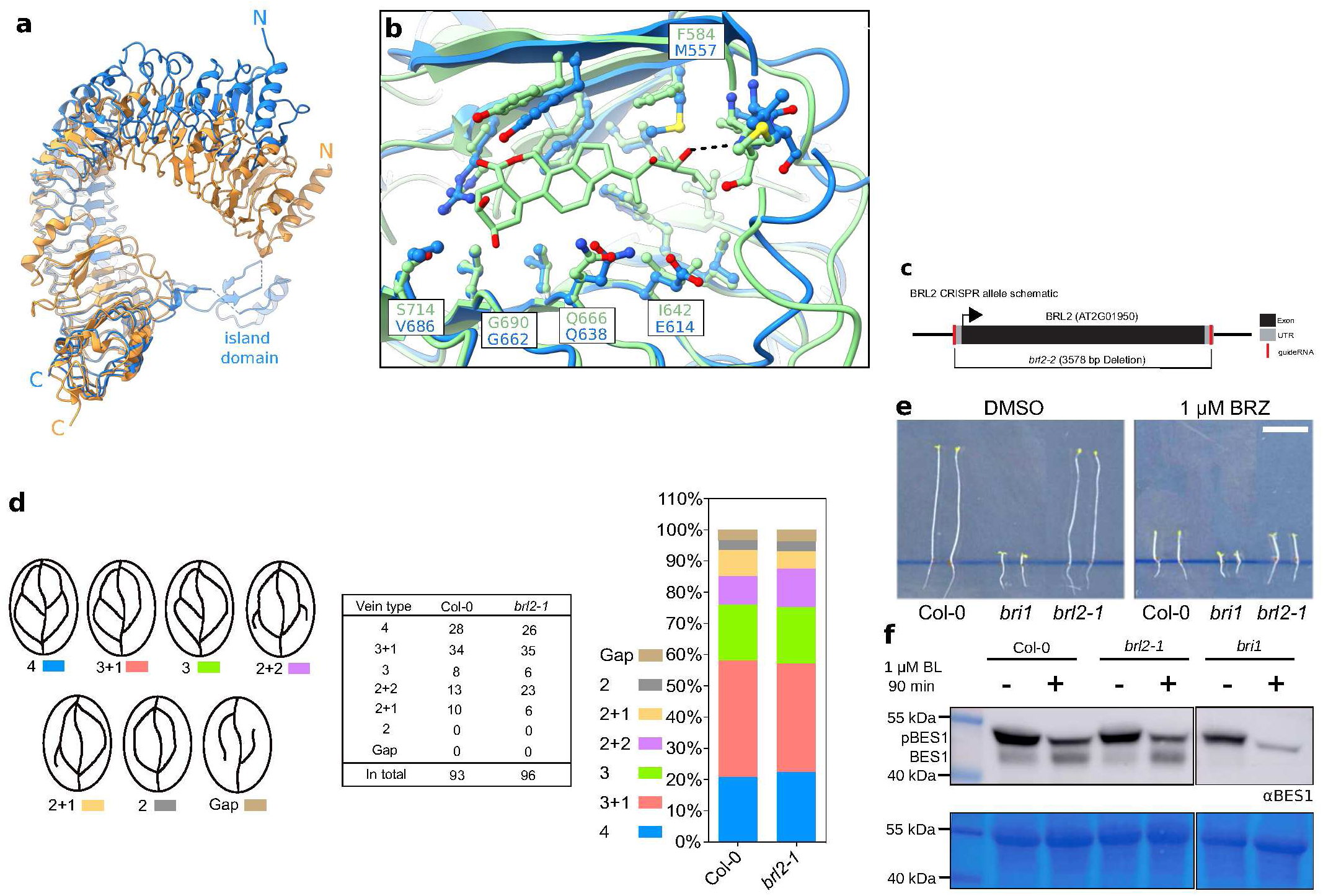
I BRL2 harbors a conserved BR-binding pocket but has not detectable function in BR growth signaling. **a,** Structural superposition of the BRIl ectodomain (ribbon diagram, in orange; PDB-ID: pdb_00003rj0) with the apo BRL2 crystal structure (in blue, r.m.s.d. ∼ 0.8 A comparing 314 C_a_ atoms). **b,** Close-up view of the BRL2 BR-binding pocket (in blue, AlphaFoldDB-ID: AF-Q9ZPS9-Fl) superimposed with BRL3 (right panel, in green, brassinolide in bonds representation, r.m.s.d. ∼ 0.7 A comparing 162 C_a_ atoms). Residues lining the BR binding site are shown in ball-and-stick representation, with corresponding residues in BRL2 shown in blue. Non-synonymous residues among BRIl, BRL2 and BRL3 are labeled. **c,** Schematic diagram of *brl2-2* CRISPR/Cas9 deletion allele. The BRL2 genomic locus is shown, including the translation start codon (black arrow), exon (black rectangle) and 5’ and 3- primer untranslated regions (UTRs, gray blocks). The putative binding sites of the designed guide RNAs are indicated in red. **d,** Cotyledon vein patterning analysis performed using 2-week-old Col-0 and *brl2-l* seedlings. Leaf patterns are categorized in seven groups, indicated 4 (4 distinct regions are formed by the intersection of the leaf veins, blue), 3+l (3 distinct regions formed with l free branch, beige), 3 (3 distinct regions formed, lemon green), 2+2 (2 distinct regions formed with 2 free branches, light purple), 2+l (2 distinct regions formed with l free branch, canary), 2 (2 distinct regions formed, gray) and gap (no distinct region formed, earthy yellow). **e,** Hypocotyl growth assay of dark-grown brassinazole (BRZ)^105^ treated vs untreated control seedlings. Seeds of Col-0 and *brl2-l* were germinated and grown on ^l/2^MS plates supplemented with l µM BRZ or 0.l% (v/v) DMSO (scale bar = l cm). f, BESl (theoretical molecular mass ∼ 39.2 kDa) western blot. 14-d-old Col-0, *brl2-l* and *bril* seedlings were grown on ^l/2^MS plates with l% (w/v) sucrose, transferred to ^l/2^MS liquid medium containing l% (w/v) sucrose and either l µM brassinolide (BL) or 0.01 % DMSO (as control) for 90 min, and probed with an anti-BESl antibody^41^ by western blotting.

**Supplementary Fig. 4.**
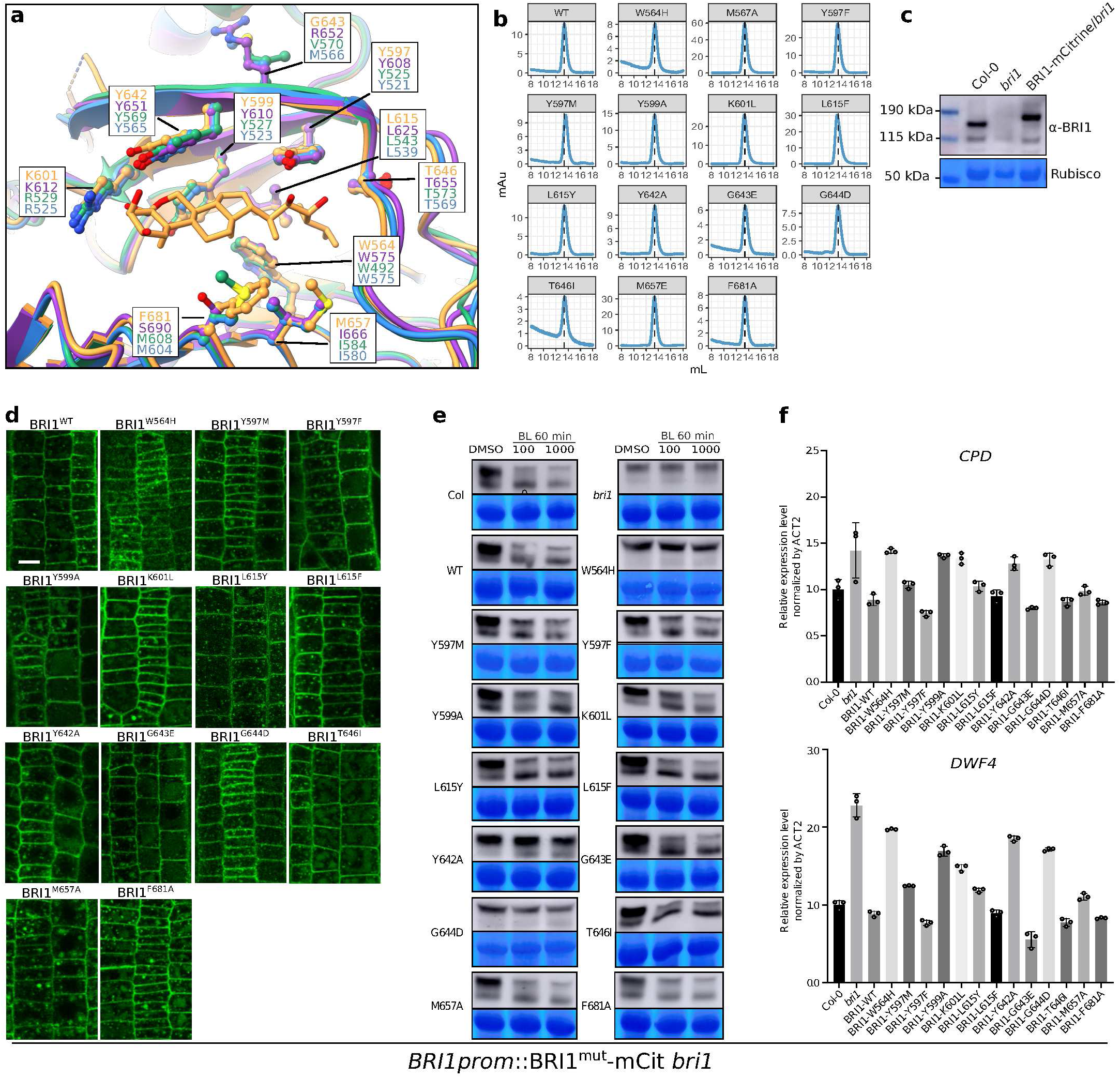
I Mutations of conserved residues in the BRil hormone binding pocket alter BR signaling without compromising the plasma-membrane localization of the mutant receptor. **a**, Structural superposition of the brassinolide-bound AtBRIl hormone binding pocket (*Arabidopsis thaliana*, in orange, PDB-ID: pdb_00003rj0) with the corresponding regions in ZmBRIl (*Zea mays*, in blue, AlphaFoldDB-ID: AF-A0A3L6FIW6-Fl, r.m.s.d ∼ 0.7 A between 175 C_a_ atoms), SlBRIl (*Solanum lycopersicum*, in purple, AlphaFoldDB-ID: AF-F2XYF6-Fl, r.m.s.d ∼ 0.6 A between 178 C_a_ atoms), and OsBRIl (*Oryza sativa*, in green, AlphaFoldDB-ID: AF-Q942F3- Fl, r.m.s.d ∼ 0.7 A between 170 C_a_ atoms). Mutated residues in AtBRIl are displayed as ball-and- stick models, the corresponding amino acid side-chains from ZmBRIl, SlBRIl, and OsBRIl are included. **b,** Analytical size-exclusion chromatography elution profiles for recombinant wild-type (WT) and point-mutant BRIl ectodomains used in the GCI assays. **c,** BRIl protein level in Col-0, *bril* and *BRIlprom*::BRIl-mCitrine / *bril* lines, detected with a primary anti-BRIl antibody^106^. ReadyBlue-stained RuBisCO is shown as loading control below. **d,** Confocal imaging of T2 generation *BRIlprom*::BRIl^mut^-mCitrine signal in the mersistematic zone of 10-d-old primary roots mounted for confocal imaging (scale bar = 20 µm). **e,** Brassinolide (BL)-stimulated BESl dephosphorylation. 14-d-old seedlings of Col-0 wild-type, *bril* and *BRIlprom*::BRIl^mut^-mCit T3 complementation lines were treated with 0.l% (v/v) DMSO or 100 nM BL and 1000 nM BL for 60 min before protein extraction and anti-BESl antibody^41^ immuno-blotting. The upper band represents phosphorylated BESl, the lower band (more) dephosphorylated BESl. ReadyBlue- stained RuBisCO is shown as loading control below. **f,** qRT-PCR analysis of the BR marker genes CPD and DWF4 in Col-0 wild-type, *bril* and *BRIlprom*::BRIl^mut^-mCit T3 complementation, with ACT2 used as internal reference gene. Each dot represents a biological replicate (n=3).

**Extended Data Fig. 1.**
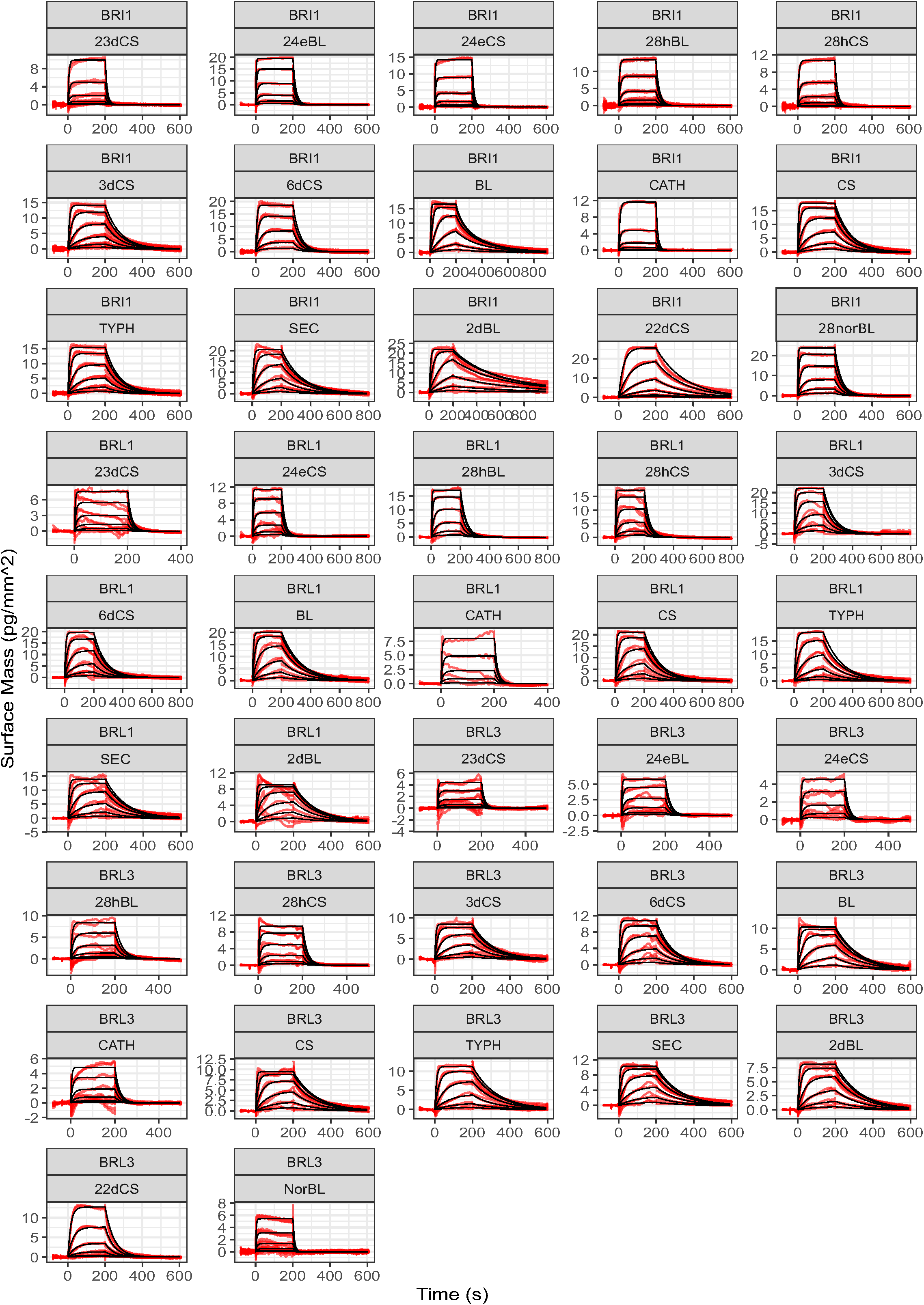
I Complete set of GCi sensorgrams for the experiments described in Fig. 2a, c, d. Grating Coupling Interferometry (GCI)-derived binding kinetics for BRIl, BRLl, and BRL3 in response to different BRs. Raw sensorgrams are shown with experimental traces in red, and global fits as black lines. CATH: cathasterone, 23dCS: 23-deoxycastasterone, 24eCS: 24-epicastasterone, 28hCS: 28-homocastasterone, 24eBL: 24-epibrassinolide, 28hBL: 28- homobrassinolide, 22dCS: 22-deoxycastasterone, 6dCS: 6-deoxocastasterone, NorBL: 28- norbrassinolide, 3dCS: 3-deoxycastasterone, TYPH: typhasterol, CS: castasterone, SEC: secasterone, 2dBL: 2-deoxybrassinolide, BL: brassinolide.

**Extended Data Fig. 2.**
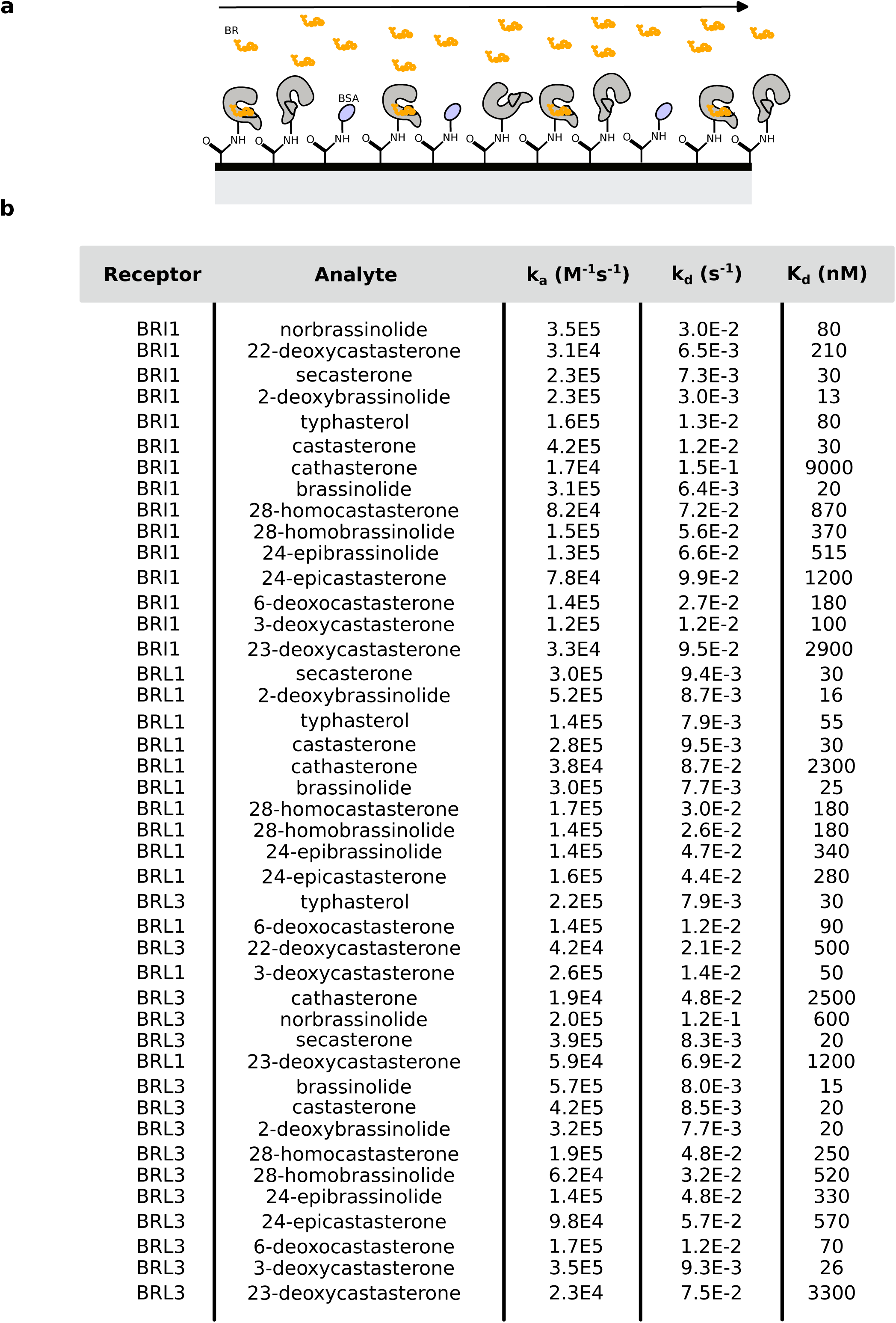
I Complete set of GCi kinetic parameters for the experiments described in Fig. 2a, c, d. **a,** Schematic illustration of the Grating Coupling Interferometry (GCI) setup used to screen receptor - brassinosteroid interactions. **b,** Kinetic parameters obtained for BRIl, BRLl, and BRL3 in response to the 15 BRs analyzed in this study.

**Extended Data Fig. 3.**
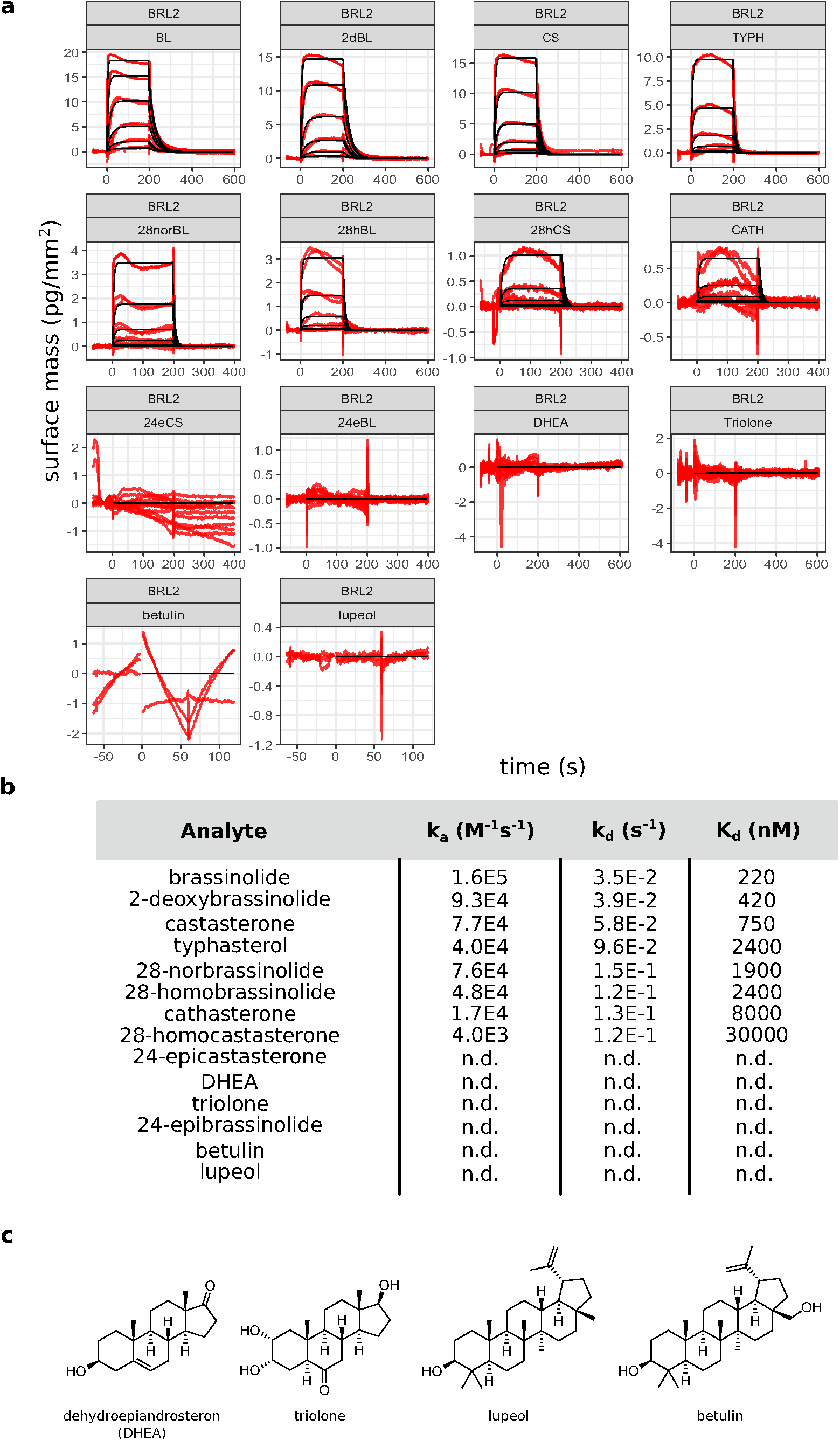
I Complete set of GCi sensorgrams and derived kinetic parameters for the experiments described in Fig. 4b, c. **a,** Grating Coupling Interferometry (GCI)-derived binding kinetics for BRL2 in response to different BRs and other plant steroids. Raw sensorgrams are shown with experimental traces in red (n=2), and global fits as black lines (n.d., no detectable binding). BL: brassinolide, 2dBL: 2-deoxybrassinolide, CS: castasterone, TYPH: typhasterol, 28norBL: 28-norbrassinolide, 28hBL: 28-homobrassinolide, 28hCS: 28-homocastasterone, CATH: cathasterone, 24eCS: 24-epicastasterone, 24eBL: 24-epibrassinolide, DHEA: dehydroepiandrosterone. **b,** Kinetic parameters obtained for BRL2 in response to the different BR and non-BR ligand candidates analyzed in this study. **c,** Chemical structures of dehydroepiandrosterone (DHEA), triolone, lupeol and betulin^ll3-ll6^.

**Extended Data Fig. 4.**
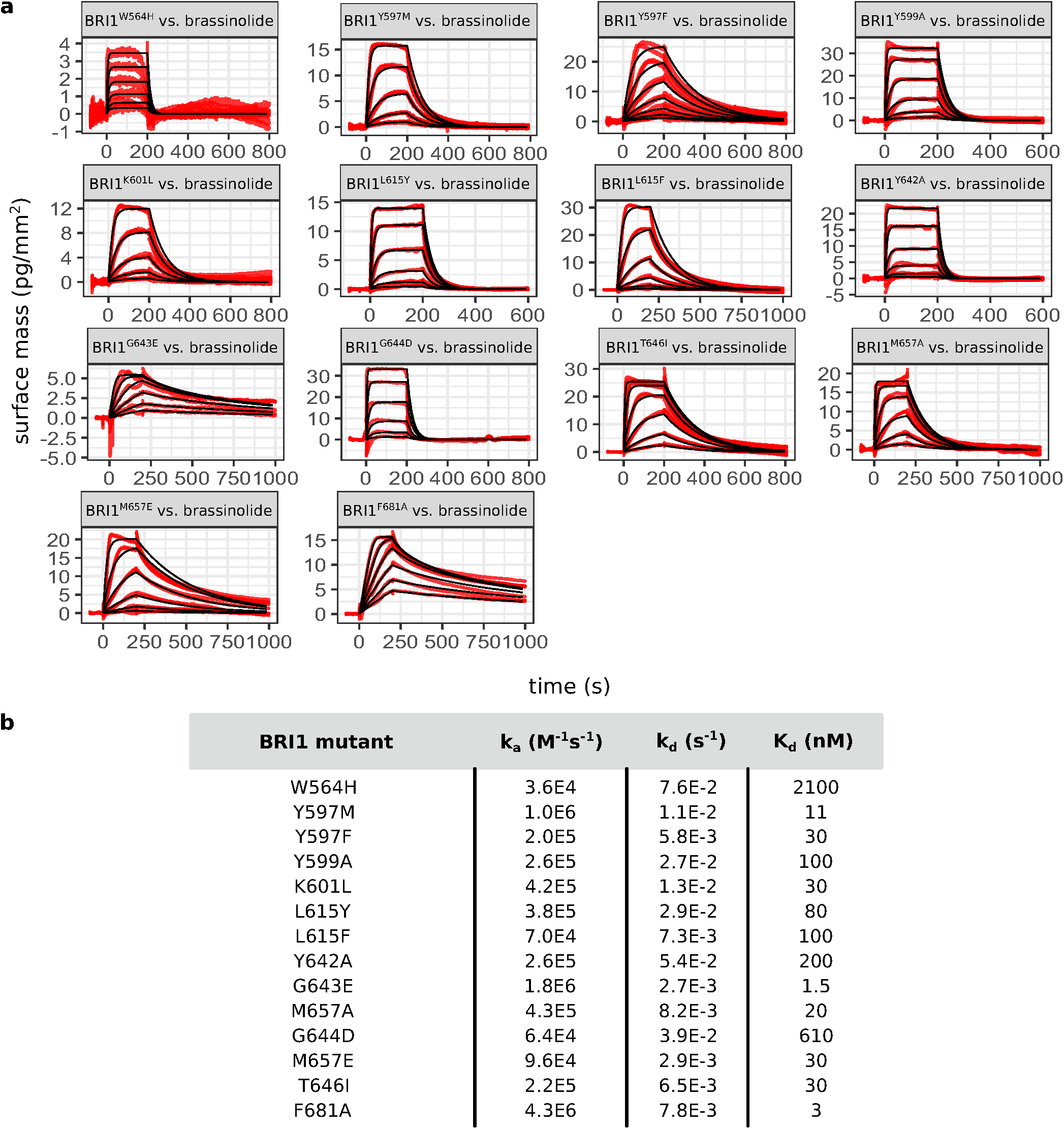
I Complete set of GCi sensorgrams and derived kinetic parameters for the experiments described in Fig. 5b,c. **a,** Grating Coupling Interferometry (GCI)-derived binding kinetics for different BRIl ectodomains carrying point mutations in the BR-binding pocket vs. brassinolide. Raw sensorgrams are shown with experimental traces in red, and global fits as black lines. **b,** Table summaries of kinetic parameters obtained for the experiments shown in **a**.

**Extended Data Fig. 5.**
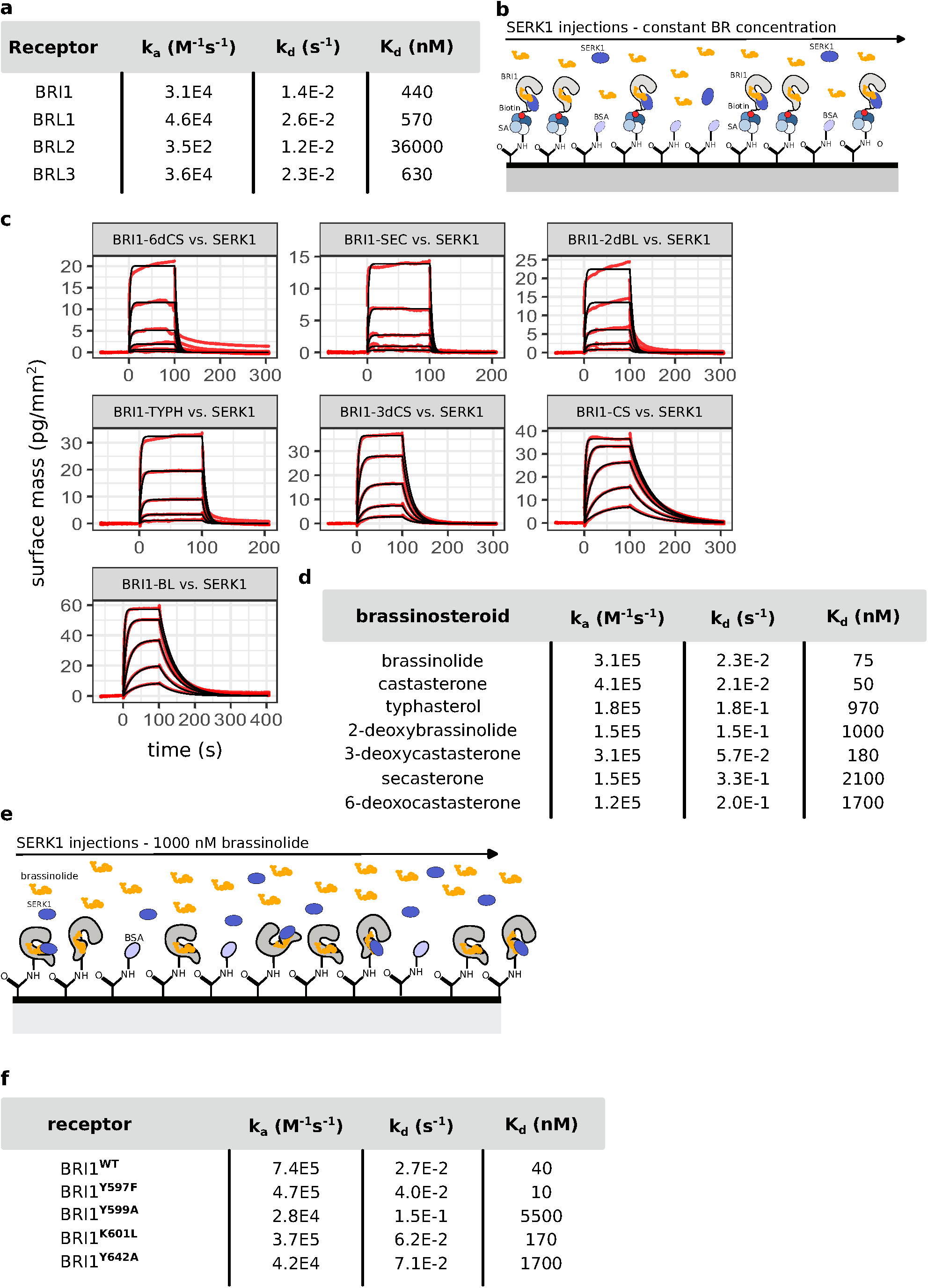
I Complete set of GCi sensorgrams and derived kinetic parameters for experiments shown in Fig. 6b-f. **a,** Table summaries of kinetic parameters obtained for the experiments shown in Fig. 6b. **b,** Schematic overview of the GCI receptor capturing and SERKl binding assay. For Avitag based coupling, streptavidin (SA, shown as a tetramer) was immobilized using the amine-coupling method on the GCI chip (black line). Next, the biotinylated avi-tagged BR receptor ectodomain was captured by streptavidin, followed by passivation and quenching of the surface with BSA (in light blue). Then, in the presence of a constant concentration of brassinolide in the analyte buffer (in yellow), SERKl (in dark blue) association and dissociation at different protein concentrations was performed. **c,** Grating Coupling Interferometry (GCI)-derived binding kinetics for the avi-tagged BRIl ectodomain vs. different antagonist candidate ligands including 6dCS: 6- deoxocastasterone, SEC: secasterone, TYPH: typhasterol, 3dCS: 3-deoxocastasterone, CS: castasterone, BL: brassinolide. Raw sensorgrams are shown with experimental traces in red, and global fits as black lines. **d,** Table summaries of kinetic parameters obtained for the experiments shown in Fig. 6c,d. **e**, Schematic overview of the GCI receptor capturing and SERKl binding assay to analyze the effect of BR binding site missense mutations in BRIl for SERKl co-receptor binding (related to Fig. 6f). BRIl^WT^, BRIl^Y597F^, BRIl^Y599A^, BRIl^K601L^ and BRIl^Y642A^ were immobilized by amine-coupling on the GCI chip (black line). SERKl binding to the receptors (in gray) was then quantified in the presence of a constant, high concentration of brassinolide (in yellow) in the running buffer and by injecting the co-receptor at different concentrations (in blue). **f,** Table summaries of kinetic parameters obtained for the experiments shown in Fig. 6f.

**Supplementary Table 1. I Crystallographic data collection and refinement statistics.**

**Supplementary Table 2. List of genotyping primers used in this study.**

**Supplementary Table 3. Cloning primers used in this study**

**Supplementary Table 4. I qPCR primers for BR marker genes CPD and DWF4**

**Extended Data Table l. I immobilized receptor density and analyte concentrations used for the GCi assays described in Fig. 2a,c,d**.

**Extended Data Table 2. I BR-mediated root growth inhibition raw data corresponding to Fig. 3c.**

**Extended Data Table 3. I RNA-seq raw counts used to calculate the heat map shown in Fig. 3e.**

**Extended Data Table 4. I immobilized receptor density and analyte concentrations used for the GCi assays described in Fig. 4b,c**.

**Extended Data Table 5. I immobilized receptor density and analyte concentrations used for the GCi assays described in Fig. 5.b,c.**

**Extended Data Table 6. I Wild-type and BRil^mut^ seedling root length raw data corresponding to Fig. 5f.**

**Extended Data Table 7. I immobilized receptor density and analyte concentrations used for the GCi assays described in Fig. 6b,c,d and g.**

**Extended Data Table 8. I BR-mediated root growth inhibition antagonist assay raw data corresponding to Fig. 6g**.

## Supplementary Note 1: Synthesis

### General methods

The NMR spectra were taken on a JEOL JNM-ECA 500 (JEOL, Tokyo, Japan; ^1^H, 500 MHz; ^13^C, 125 MHz) spectrometer equipped with a 5mm JEOL Royal probe. ^1^H NMR and ^13^C NMR chemical shifts (’6) were calibrated using tetramethylsilane (TMS, ^1^H ‘6=0 ppm) or solvents: CDCl_3_ (^1^H ‘6=7.27 ppm, ^13^C ‘6=77.00 ppm) or DMSO-d_6_ (^1^H ‘6=2.46 ppm, ^13^C ‘6=40.00 ppm).

Chemical shifts are given in ppm (’6-scale), coupling constants (J) in Hz. All values were obtained by first order analysis. The NMR data were processed using the ACD/NMR Processor Academic Edition (ver. 12.01). For ESI or APCI HRMS analysis, the samples were dissolved in chloroform or methanol. Samples were analyzed with 1290 Infinity II Liquid Chromatographer (Agilent) with HPLC column infinityLab Poroshell 120 EC-C18 (4.6×50 mm; 2.7 µm). For API HRMS analysis, the samples were dissolved in chloroform (or chloroform:methanol; 1:1; v/v, in the case of hydroxylated compounds) to a concentration 10 µg·mL^-1^. The ASAP (Atmospheric Solids Analysis Probe) was dipped into the sample solution, placed into the ion source and analyzed in full scan mode. The source of the Synapt G2-Si Mass Spectrometer (Waters, Manchester, UK) was operated in positive ionization mode (ASAP+) and, if not stated otherwise, at source temperature of 120 °C. The Corona needle current was kept at 5 µA and the collision energy at value 4. The probe temperature was ramped up from 50 °C to 600 °C in 3 min. Data were acquired from 50 to 1000 Da with 1.0 s scan time in High Resolution Mode and processed using the Masslynx 4.1 software (Waters). Mass accuracy of 1 ppm or less was achieved with the described instrumentation for all compounds.

Merck silica gel Kieselgel 60 (230-400 mesh) was used for column chromatography. The HPLC system consisted of a Waters semi-preparative HPLC system including quaternary pump, liquid handler, UV-VIS and ELSD detectors. The HPLC semi preparative column was filled with silica gel. Reagents and solvents were purchased from Sigma-Aldrich and were not purified.

**Scheme 1:**
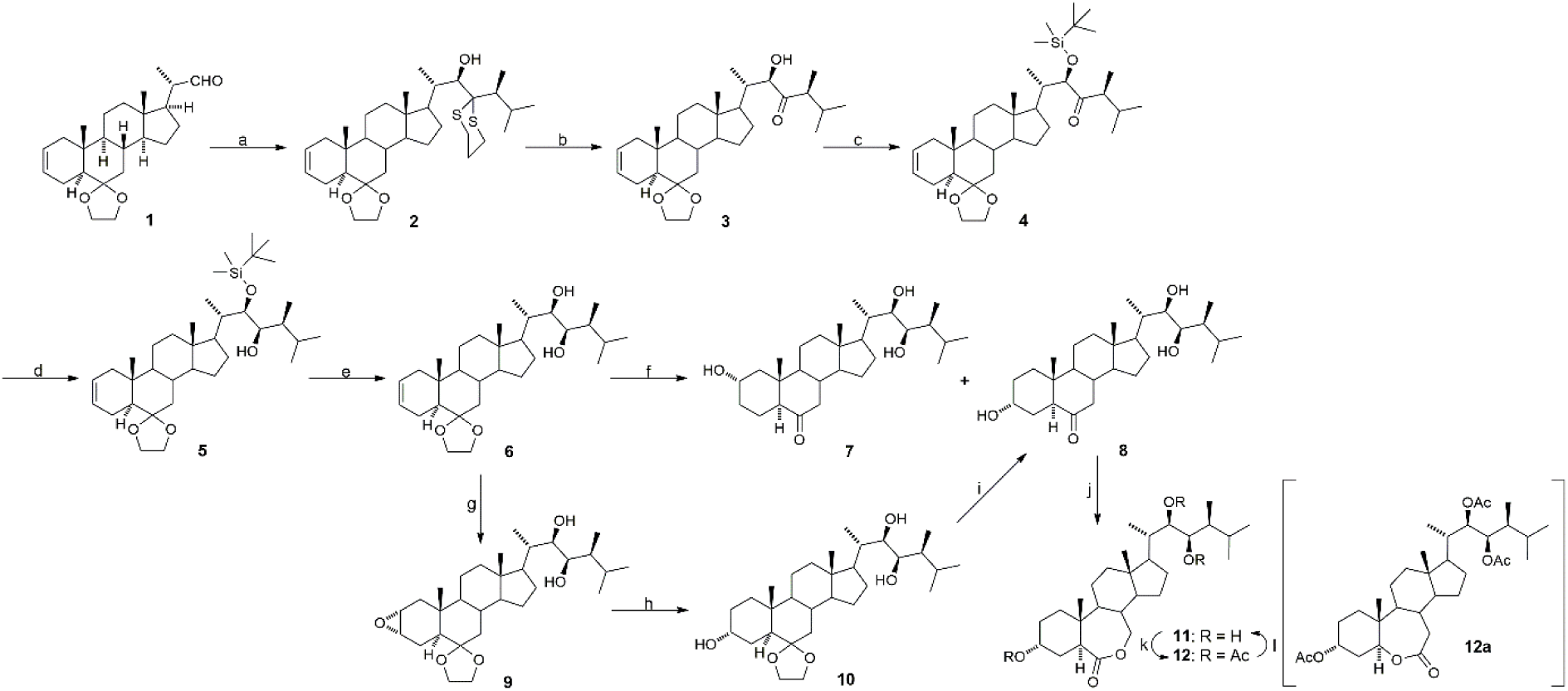
Synthesis of 3-deoxycastasterone (**7**), typhasterol (**8**), and 2-deoxybrassinolide (**11**). **a**: (*S*)-2-(3-methylbutan-2-yl)-1,3-dithiane/n-BuLi/THF, r.t. then -78 °C, 1h; **b**: NaH_2_PO_4_/2- methyl-2-butene/ NaClO_2_/H_2_O/ethanol, 0 °C, 10 min; **c**: TBSOTf/2,6-lutidine/CH_2_Cl_2_, 0 °C, 1 h; **d**: DIBAL-H/toluen, -78 °C, 1h; **e**: TBAF/THF, r.t., 1h; **f**: 1, BH_3_/THF, r.t., 6h; 2, NaOH/H_2_O_2_/H_2_O/THF, r.t., 12h; **g**: MCPBA/EtOAc, r.t., 4h; **h**: LiAlH_4_/THF/Et_2_O, 40 °C, 3h; **i**: HCl/H_2_O/THF, 40 °C, 4h; **j**: TFPAA/CH_2_Cl_2_, r.t., 4h; **k**: pyridine/Ac_2_O, 50 °C, 12h; **l**: MeONa/MeOH/THF, r.t., 1h. Note: absolute configuration for carbons 8, 9, 13, and 17 is shown only for compound **1** but is identical for all compounds prepared.

### Synthesis of (22*R*)-6,6-ethylenedioxy-23,23-(trimethylenedithio)-5*a*-campest-2-en-22-ol (2)

1.6 M solution of *n*-BuLi in hexane (3.0 mL, 4.8 mmol) was added to a solution of (*S*)-2-(3-methylbutan-2-yl)-1,3-dithiane^1^ (790 mg, 4.0 mmol) in THF (15 mL) at room temperature. The mixture was stirred for 30 minutes and then cooled to -78 °C. A solution of aldehyde **1**^2^ (1.0 g, 2.69 mmol) in THF (3 mL) was added dropwise to the reaction mixture. The solution was stirred at -78 °C for 30 min, quenched with saturated aqueous NH_4_Cl (20 mL), and warmed to room temperature. The product was extracted with ethyl acetate. The organic layer was washed with water followed by drying over MgSO_4_, filtrated and evaporated under reduced pressure. The residue was purified by column chromatography on silica gel (3-5% ethyl acetate in cyclohexane) to give alcohol **2** (1.22 g, 81 %) as a viscous oil. ^1^H NMR (500 MHz, DMSO-d_6_): ‘6 5.63 (m, 1H, H-3), 5.51 (m, 1H, H-2), 4.50 (d, 1H, *J* = 6.4 Hz, H-22), 3.92-3.79 (m, 3H, H_ketal_), 3.69 (q, 1H, *J* = 6.6 Hz, H_ketal_), 2.83-2.60 (m, 5H), 2.12-1.69 (m, 11H), 1.62-1.23 (m, 8H), 1.15-0.75 (m, 4H), 1.11 (d, 3H, *J* = 6.4 Hz, CH_3_), 0.99 (d, 3H, *J* = 7.0 Hz, CH_3_), 0.89 (d, 3H, *J* = 6.7 Hz, CH_3_), 0.86 (d, 3H, *J* = 6.7 Hz, CH_3_), 0.80 (s, 3H, CH_3_), 0.66 (s, 3H, CH_3_). ^13^C NMR (125 MHz, DMSO-d_6_): 125.45, 124.86, 108.99, 73.86, 65.27, 63.78, 63.56, 55.49, 53.79, 52.49, 47.57, 43.26, 42.13, 40.80, 40.62, 39.46, 38.13, 35.37, 32.99, 27.73, 27.25, 26.33, 25.51, 24.75, 24.39, 23.73, 21.07, 20.49, 18.44, 14.58, 13.53, 11.59, 9.87. HRMS (ESI+) calculated for C_33_H_55_O_3_S_2_ ([M + H]^+^) 563.3593, found 563.3597.

### Synthesis of (22*R*)-6,6-ethylenedioxy-22-hydroxy-5*a*-campest-2-en-23-one (3)

A solution of NaH_2_PO_4_ (511 mg, 4.26 mmol) in water (11 mL), 2-methyl-2-butene (2.25 mL, 21.3 mmol) and NaClO_2_ (80%, 1.44 g, 16.0 mmol) were sequentially added at 0 °C to a stirred solution of compound **2** (1.2 g, 2.13 mmol) in ethanol (100 mL). The reaction mixture was stirred for 10 min and quenched with aqueous saturated Na_2_S_2_O_3_ (mL). The mixture was diluted with ethyl acetate and extracted with water. The organic layer was dried over MgSO_4_, filtrated and evaporated under reduced pressure. The residue was purified by column chromatography on silica gel (10% ethyl acetate in cyclohexane) to give hydroxyketone 3 (736 mg, 73%) as a viscous oil. ^1^H NMR (500 MHz, CDCl_3_): ‘6 5.66 (m, 1H, H-3), 5.54 (m, 1H, H-2), 4.33 (dd, 1H, *J* = 5.5, 1.5 Hz, H-22), 4.01-3.90 (m, 3H, H_ketal_), 3.80 (q, 1H, *J* = 6.6 Hz, H_ketal_), 3.31 (d, 1H, *J* = 5.5 Hz, OH), 2.45 (p, 1H, *J* = 7.3 Hz, H-24), 2.13-1.60 (m, 12H), 1.57-0.77 (m, 9H), 1.08 (d, 3H, *J* = 7.3 Hz, CH_3_), 0.94 (d, 3H, *J* = 6.7 Hz, CH_3_), 0.892 (s, 3H, CH_3_), 0.887 (d, 3H, *J* = 6.7 Hz, CH_3_), 0.75 (s, 3H, CH_3_), 0.74 (d, 3H, *J* = 6.7 Hz, CH_3_). ^13^C NMR (125 MHz, CDCl_3_): 216.23, 125.64, 124.78, 110.00, 77.31, 65.61, 64.09, 55.86, 53.27, 52.52, 48.11, 48.02, 42.35, 41.15, 41.17, 39.47, 38.18, 35.85, 33.43, 28.70, 28.25, 24.00, 21.50, 21.41, 20.80, 19.03, 15.18, 13.59, 12.52, 11.91. HRMS (ESI+) calculated for C_30_H_49_O_4_ ([M + H]^+^) 473.3631, found 473.3631.

### Synthesis of (22*R*)-6,6-ethylenedioxy-22-((*tert*-butyldimethylsilyl)oxy)-5*a*-campest-2-en- 23-one (4)

TBSOTf (670 µl, 2.92 mmol) was added at 0 °C to a stirred solution of hydroxyketone **3** (700 mg, 1.48 mmol) and 2,6-lutidine (850 µL, 7.3 mmol) in CH_2_Cl_2_ (10 mL). The reaction mixture was stirred for 1 hour at 0 °C and then quenched with saturated aqueous NaHCO_3_ (10 mL). The mixture was diluted with ethyl acetate and extracted with water. The organic layer was dried over MgSO_4_, filtrated and evaporated under reduced pressure. The residue was purified by column chromatography on silica gel (5% ethyl acetate in cyclohexane) to give silyl ether **4** (811 mg, 94%) as a viscous oil. ^1^H NMR (500 MHz, CDCl_3_): ‘6 5.67 (m, 1H, H-3), 5.55 (m, 1H, H-2), 4.31 (d, 1H, *J* = 0.9 Hz, H-22), 4.02-3.90 (m, 3H, H_ketal_), 3.80 (q, 1H, *J* = 6.6 Hz, H_ketal_), 2.38 (dq, 1H, *J* = 8.6, 7.0 Hz, H-24), 2.14-1.60 (m, 11H), 1.57-0.75 (m, 10H), 1.02 (d, 3H, *J* = 7.0 Hz, CH_3_), 0.93 (s, 9H, 3×CH_3_), 0.92 (d, 3H, *J* = 6.7 Hz, CH_3_), 0.89 (s, 3H, CH_3_), 0.83 (d, 3H, *J* = 6.7 Hz, CH_3_), 0.81 (d, 3H, *J* = 6.7 Hz, CH_3_), 0.73 (s, 3H, CH_3_), 0.06 (s, 3H, CH_3_), -0.01 (s, 3H, CH_3_). ^13^C NMR (125 MHz, CDCl_3_): 214.65, 125.65, 124.82, 110.03, 79.17, 65.62, 64.10, 56.17, 53.26, 52.37, 48.49, 48.08, 42.37, 41.20, 41.15, 39.62, 38.53, 35.88, 33.41, 29.11, 28.56, 26.89, 25.88 (3×C), 24.13, 21.56, 21.45, 20.79, 19.39, 15.94, 13.59, 12.87, 11.74, -4.22, -5.00. HRMS (ESI+) calculated for C_36_H_63_O_4_Si ([M + H]^+^) 587.4496, found 587.4491.

### Synthesis of (22*R,*23*R*)-6,6-ethylenedioxy-22-((*tert*-butyldimethylsilyl)oxy)-5*a*-campest-2- en-23-ol (5)

1M Solution of DIBAL-H in toluene (6.4 mL, 6.4 mmol) was added to a stirred solution of compound **4** (750 mg, 1.28 mmol) in dry toluene (30 mL) at -78 °C. The reaction mixture was stirred at the same temperature for 1 hour and then quenched with MeOH (6 mL). Saturated aqueous solution of sodium potassium tartrate (30 mL) was added and the mixture was diluted with ethyl acetate. The aqueous layer was separated and extracted twice with ethyl acetate. The combined organic layers were washed with water, dried over MgSO_4_, filtrated and evaporated under reduced pressure. The residue was purified by column chromatography on silica gel (2- 5% ethyl acetate in cyclohexane) to give alcohol **5** (715 mg, 95%) as a viscous oil (containing 12% of inseparable 23*S*-isomer). ^1^H NMR (500 MHz, CDCl_3_): ‘6 5.56 (m, 1H, H-3), 5.53 (m, 1H, H-2), 3.99-3.88 (m, 3H, H_ketal_), 3.78 (q, 1H, *J* = 6.6 Hz, H_ketal_), 3.58 (d, 1H, *J* = 7.5 Hz, H-23), 3.49 (dd, 1H, *J* = 7.5, 2.9 Hz, H-22), 2.11-1.85 (m, 6H), 1.78-1.37 (m, 9H), 1.24-0.75 (m, 8H), 0.94 (d, 3H, *J* = 6.7 Hz, CH_3_), 0.92 (s, 9H, 3×CH_3_), 0.90 (d, 3H, *J* = 6.7 Hz, CH_3_), 0.89 (d, 3H, *J* = 6.7 Hz, CH_3_), 0.87 (s, 3H, CH_3_), 0.83 (d, 3H, *J* = 7.0 Hz, CH_3_), 0.68 (s, 3H, CH_3_), 0.12 (s, 3H, CH_3_), 0.10 (s, 3H, CH_3_). ^13^C NMR (125 MHz, CDCl_3_): 125.67, 124.80, 110.02, 76.09, 73.02, 65.59, 64.09, 56.18, 53.33, 52.21, 48.07, 42.54, 41.21, 41.15, 40.91, 39.75 (2×C), 35.88, 33.34, 30.79, 28.55, 26.14 (3×C), 24.21, 21.44, 21.04, 20.81, 19.88, 18.60, 13.60, 12.91, 11.61, 10.00, -3.57, -4.09. HRMS (APCI+) calculated for C_36_H_65_O_4_Si ([M + H]^+^) 589.4652, found 589.4651.

### Synthesis of (22*R,*23*R*)-6,6-ethylenedioxy-5*a*-campest-2-ene-22,23-diol (6)

1M Solution of TBAF (6.5 mL, 6.5 mmol) was added to a solution of compound **5** (700 mg, 1.19 mmol) in THF (30 mL). The reaction mixture was stirred at room temperature for 1 hour. Then, it was diluted with ethyl acetate and washed with saturated solution of NH_4_Cl (20 mL). The combined organic layers were washed with water, dried over MgSO_4_, filtrated and evaporated under reduced pressure. The residue was purified by column chromatography on silica gel (25% ethyl acetate in cyclohexane) to give diol **6** (524 mg, 93%) as white solid. ^1^H NMR (500 MHz, CDCl_3_): ‘6 5.67 (m, 1H, H-3), 5.55 (m, 1H, H-2), 4.01-3.90 (m, 3H, H_ketal_), 3.79 (q, 1H, *J* = 6.6 Hz, H_ketal_), 3.73 (dd, 1H, *J* = 8.4, 2.0 Hz, H-23), 3.57 (dd, 1H, *J* = 8.4, 1.1 Hz, H-22), 2.12-1.91 (m, 5H), 1.81-1.37 (m, 10H), 1.31-0.76 (m, 7H), 0.98 (d, 3H, *J* = 6.7 Hz, CH_3_), 0.96 (d, 3H, *J* = 6.7 Hz, CH_3_), 0.91 (d, 3H, *J* = 6.7 Hz, CH_3_), 0.86 (s, 3H, CH_3_), 0.86 (d, 3H, *J* = 7.0 Hz, CH_3_), 0.71 (s, 3H, CH_3_). ^13^C NMR (125 MHz, CDCl_3_): 125.65, 124.78, 110.03, 74.87, 73.42, 65.59, 64.08, 55.87, 53.36, 52.40, 48.03, 42.36, 41.18 (2×C), 39.95, 39.79, 36.79, 35.86, 33.40, 30.75, 27.71, 23.99, 21.40, 20.86 (2×C), 20.77, 13.59, 11.88 (2×C), 10.12. HRMS (ESI+) calculated for C_30_H_51_O_4_ ([M + H]^+^) 475.3787, found 475.3787.

### Synthesis of (22*R,*23*R*)-2**α**,22,23-trihydroxy-5*a*-campestan-6-one (3-deoxycastasterone, 7) and (22*R,*23*R*)-3**α**,22,23-trihydroxy-5*a*-campestan-6-one (typhasterol, 8)

1M Solution of borane in THF (1 mL, 1.0 mmol) was added dropwise to the solution of olefin **6** (50 mg, 0.1 mmol) in dry THF (2 mL). The reaction mixture was stirred at room temperature for 6 h. Then, the reaction mixture was quenched by water (2 mL) and cooled to 10 °C. Aqueous solution of NaOH (10%, 1 mL) was added followed by dropwise addition of 30% hydrogen peroxide (2 mL). This reaction mixture was stirred at room temperature overnight. It was then poured into a mixture of sodium sulfite (500 mg) in water (5 mL) and acetic acid (0.5 mL) and stirred for additional 30 min. After then, the reaction mixture was diluted with ethyl acetate and extracted twice with water. The organic layer was dried over MgSO_4_ and evaporated under reduced pressure. A crude mixture of products was dissolved in a mixture of THF and acetone (5:1, 3 mL) and 5% aqueous HCl (1 mL) was added. The reaction mixture was heated under reflux for 3 h. Then it was diluted with ethyl acetate and extracted twice with water. The organic layer was again dried over MgSO_4_ and evaporated under reduced pressure. The residue was purified by HPLC (75% ethyl acetate in cyclohexane) to give two alcohols, **7** (19 mg, 40%) and **8** (15 mg, 32%), as white solids.

Compound **7** (3-deoxycastasterone): ^1^H NMR (500 MHz, CDCl_3_): ‘6 3.78 (m, 1H, H-2β), 3.73 (dd, 1H, *J* = 8.4, 1.8 Hz, H-23), 3.56 (dd, 1H, *J* = 8.4, 1.2 Hz, H-22), 2.30 (dd, *J* = 13.0, 4.4 Hz, H-7a), 2.17 (dd, *J* = 12.1, 2.9 Hz, H-5α), 2.09-1.94 (m, 3H), 1.85-1.46 (m, 11H), 1.38-1.09 (m, 11H), 0.97 (d, 3H, *J* = 6.7 Hz, CH_3_), 0.95 (d, 3H, *J* = 6.7 Hz, CH_3_), 0.91 (d, 3H, *J* = 6.7 Hz, CH_3_), 0.85 (d, 3H, *J* = 7.0 Hz, CH_3_), 0.76 (s, 3H, CH_3_), 0.69 (s, 3H, CH_3_). ^13^C NMR (125 MHz, CDCl_3_): 211.91, 74.68, 73.50, 67.06, 57.91, 56.58, 54.09, 52.25, 47.07, 46.63, 43.06, 42.75, 40.00, 39.43, 37.67, 36.76, 34.60, 30.73, 27.61, 23.79, 21.28, 20.86, 20.73, 19.48, 14.17, 11.91, 11.88, 10.09. HRMS (ESI+) calculated for C_28_H_49_O_4_ ([M + H]^+^) 449.3631, found 449.3627.

Compound **8** (typhasterol): ^1^H NMR (500 MHz, CDCl_3_): ‘6 4.18 (m, 1H, H-3β), 3.73 (bd, 1H, *J* = 8.3 Hz, H-23), 3.57 (bd, 1H, *J* = 8.3, 1.2 Hz, H-22), 2.74 (t, 1H, *J* = 7.9 Hz, H-5α), 2.31 (dd, *J* = 13.1, 4.6 Hz, H-7a), 2.04-1.94 (m, 3H), 1.84-1.47 (m, 15H), 1.41-1.19 (m, 6H), 1.10 (m, 1H), 0.97 (d, 3H, *J* = 6.7 Hz, CH_3_), 0.95 (d, 3H, *J* = 6.7 Hz, CH_3_), 0.92 (d, 3H, *J* = 6.7 Hz, CH_3_), 0.85 (d, 3H, *J* = 6.9 Hz, CH_3_), 0.74 (s, 3H, CH_3_), 0.69 (s, 3H, CH_3_). ^13^C NMR (125 MHz, CDCl_3_): 212.74, 74.72, 73.50, 65.42, 56.68, 53.74, 52.28, 51.65, 46.80, 42.77, 41.55, 40.01, 39.51, 38.02, 36.78, 31.64, 30.74, 28.14, 27.66, 27.64, 23.79, 21.08, 20.87, 20.75, 12.32, 11.92, 11.88, 10.01. HRMS (APCI+) calculated for C_28_H_49_O_4_ ([M + H]^+^) 449.3631, found 449.3626.

### Synthesis of (22*R,*23*R*)-2**α**,3**α**-epoxy-6,6-ethylenedioxy-5*a*-campestane-22,23-diol (9)

3-Chloroperoxybenzoic acid (77%, 375 mg, 1.68 mmol) was added to a solution of olefin **6** (200 mg, 0.42 mmol) in ethyl acetate (10 mL) and the reaction mixture was stirred at room temperature for 4 hours. The reaction was quenched with saturated aqueous solution of Na_2_SO_3_ (5 mL) and NaHCO_3_ (5 mL). The organic layer was separated, washed with water, dried over MgSO_4_, filtrated and evaporated under reduced pressure. The residue was purified by column chromatography on silica gel (25% ethyl acetate in cyclohexane) to give epoxide **9** (186 mg, 90%) as an off-white solid. ^1^H NMR (500 MHz, CDCl_3_): ‘6 3.99-3.89 (m, 3H, H_ketal_), 3.79 (m, 1H, H_ketal_), 3.72 (dd, 1H, *J* = 8.6, 1.5 Hz, H-23), 3.56 (d, 1H, *J* = 8.4 Hz, H-22), 3.25 (m, 1H), 3.10 (dd, 1H, *J* = 5.8, 4.0 Hz), 2.14 (ddd, 1H, *J* = 14.8, 4.1, 1.2 Hz), 2.09-1.96 (m, 2H), 1.97 (dt, 1H, *J* = 12.7, 3.3 Hz), 1.92 (dd, 1H, *J* = 14.8, 6.1 Hz), 1.81 (m, 1H), 1.75 (dd, 1H, *J* = 13.1, 3.4 Hz), 1.68-0.67 (m, 17H), 0.97 (d, 3H, *J* = 6.7 Hz, CH_3_), 0.95 (d, 3H, *J* = 6.7 Hz, CH_3_), 0.92 (d, 3H, *J* = 6.7 Hz, CH_3_), 0.97 (d, 3H, *J* = 6.7 Hz, CH_3_), 0.95 (d, 3H, *J* = 6.7 Hz, CH_3_), 0.90 (d, 3H, *J* = 6.7 Hz, CH_3_), 0.89 (s, 3H, CH_3_), 0.85 (d, 3H, *J* = 6.9 Hz, CH_3_), 0.69 (s, 3H, CH_3_). ^13^C NMR (125 MHz, CDCl_3_): 109.78, 74.83, 73.39, 65.47, 64.08, 55.71, 53.06, 52.83, 52.35, 50.46, 43.22, 42.25, 40.91, 39.97, 39.68, 39.59, 36.80, 34.73, 33.10, 30.74, 27.69, 23.99, 20.87, 20.81, 20.77, 20.07, 14.73, 11.89, 11.84, 10.13. HRMS (ESI+) calculated for C_30_H_51_O_5_ ([M + H]^+^) 491.3736, found 491.3739.

### Synthesis of (22*R,*23*R*)-6,6-ethylenedioxy-5*a*-campestane-3**α**,22,23-triol (10)

1M Solution of LiAlH_4_ in diethylether (1 mL, 1.0 mmol) was added dropwise to a solution of compound **9** (130 mg, 0.27 mmol) in dry THF (6 mL). The reaction mixture was stirred at 40 °C for 3 hours and then quenched with MeOH (3 mL). Saturated aqueous solution of sodium potassium tartrate (10 mL) was added and the mixture was diluted with ethyl acetate. The aqueous layer was separated and extracted with ethyl acetate (2×). The combined organic layers were washed with water, dried over MgSO_4_, filtrated and evaporated under reduced pressure. The residue was purified by column chromatography on silica gel (90% ethyl acetate in cyclohexane) to give triol **10** (114 mg, 87%) as a viscous oil. ^1^H NMR (500 MHz, CDCl_3_): ‘6 4.13 (m, 1H, H-3β), 4.02-3.92 (m, 3H, H_ketal_), 3.75-3.71 (m, 2H, H-23, H_ketal_), 3.55 (d, 1H, *J* = 8.5 Hz, H-22), 2.08-1.93 (m, 3H), 1.79 (dd, 1H, *J* = 13.1, 3.7 Hz), 1.70-1.05 (m, 22H), 0.97 (d, 3H, *J* = 6.7 Hz, CH_3_), 0.95 (d, 3H, *J* = 6.7 Hz, CH_3_), 0.93 (s, 3H, CH_3_), 0.90 (d, 3H, *J* = 6.7 Hz, CH_3_), 0.85 (d, 3H, *J* = 6.9 Hz, CH_3_), 0.71 (s, 3H, CH_3_). ^13^C NMR (125 MHz, CDCl_3_): 110.59, 75.44, 72.53, 65.68, 65.44, 64.05, 56.22, 53.85, 52.66, 45.05, 42.56, 40.64, 40.09, 39.60, 37.66, 37.17, 33.82, 33.35, 30.87, 27.86, 27.51, 27.18, 24.11, 20.92, 20.87, 20.66, 13.04, 11.93, 11.87, 10.28. HRMS (ESI+) calculated for C_30_H_53_O_5_ ([M + H]^+^) 493.3893, found 493.3895.

### Synthesis of typhasterol (8) from compound 10

Aqueous solution of HCl (5%, 0.5 mL) was added to a solution of compound **10** (80 mg, 0.16 mmol) in THF (5 mL) and the reaction mixture was stirred at 40 °C for 4 hours and then diluted with ethyl acetate. Organic layer was washed twice with water, dried over MgSO_4_, filtrated and evaporated under reduced pressure. The residue was purified by column chromatography on silica gel (90% ethyl acetate in cyclohexane) to give typhasterol (**8**) (75 mg, 90%) as a white solid. The spectroscopic data were entirely consistent with those reported for typhasterol above.

### Synthesis of (22*R,*23*R*)-3**α**,22,23-trihydroxy-7a-homo-7-oxa-5*a*-campestan-6-one (2-deoxybrassinolide, 11)

Freshly prepared trifluoroperoxyacetic acid (4 mL) was added to a solution of typhasterol (**8**) (60 mg, 0.13 mmol) in dichloromethane (10 mL) and the reaction mixture was stirred at room temperature for 4 hours and quenched with saturated aqueous solutions of Na_2_SO_3_ (5 mL) and NaHCO_3_ (5 mL). The aqueous layer was separated and extracted twice with dichloromethane. The combined organic layers were washed with water, dried over MgSO_4_, filtrated and evaporated under reduced pressure to give inseparable crude mixture of isomeric lactones (65 mg) with compound **11** as the major isomer (ratio approx. 3:1). For separation the mixture had to be acetylated. Crude lactones were dissolved in pyridine (6 mL) and acetic anhydride (1 mL) was added. The reaction mixture was stirred at 50 °C for 12 hours. Then, ethanol (5 mL) and ethyl acetate (10 mL) were added and the mixture was stirred for additional 30 min. The mixture was washed twice with water, dried over MgSO_4_, filtrated and evaporated under reduced pressure. The residue was purified by HPLC (30% ethyl acetate in cyclohexane) to give two triacetates, **12** (50 mg, 63%) and **12a** (15 mg, 19%), as a viscous oil.

Compound **12** (40 mg, mmol) was dissolved in THF (5 mL) and 5M methanolic solution of sodium methoxide (0.1 mL) was added. The reaction mixture was stirred at room temperature for 1 hour and quenched with 5% aqueous solution of HCl (1 mL). The mixture was diluted with ethyl acetate, washed twice with water, dried over MgSO_4_, filtrated and evaporated under reduced pressure. The residue was purified by HPLC (ethyl acetate) to give lactone **11** (28 mg, 89%) as a white solid.

Compound **11** (2-deoxybrassinolide): ^1^H NMR (500 MHz, CDCl_3_): ‘6 4.17 (m, 1H, H-3β), 4.11-4.08 (m, 2H, H-7aα, H-7aβ), 3.73 (dd, 1H, *J* = 8.4, 1.2 Hz, H-23), 3.55 (d, 1H, *J* = 8.4 Hz, H-22), 3.19 (dd, 1H, *J* = 12.2, 4.6 Hz, H-5α), 2.14 (ddd, 1H, *J* = 14.9, 12.5, 2.3 Hz), 2.02-1.96 (m, 3H), 1.81-1.19 (m, 19H), 0.90 (m, 1H), 0.97 (d, 3H, *J* = 6.7 Hz, CH_3_), 0.95 (d, 3H, *J* = 6.7 Hz, CH_3_), 0.91 (d, 3H, *J* = 6.7 Hz, CH_3_), 0.90 (s, 3H, CH_3_), 0.85 (d, 3H, *J* = 6.9 Hz, CH_3_), 0.72 (s, 3H, CH_3_). ^13^C NMR (125 MHz, CDCl_3_): 176.78, 74.59, 73.53, 70.39, 64.88, 58.29, 52.25, 51.37, 42.44, 41.75, 40.02, 39.50, 36.79, 36.28, 32.45, 30.73, 29.69, 28.23, 27.57, 24.71, 22.15, 20.85, 20.74, 14.55, 14.12, 11.82, 11.72, 10.10. HRMS (APCI+) calculated for C_28_H_49_O_5_ ([M + H]^+^) 465.3580, found 465.3574.

Compound **12** ((22*R,*23*R*)-7a-homo-7-oxa-6-oxo-5a-campestan-3α,22,23-triyl triacetate): ^1^H NMR (500 MHz, CDCl_3_): ‘6 5.33 (dd, 1H, *J* = 8.9, 1.2 Hz, H-23), 5.15 (d, 1H, *J* = 8.9 Hz, H-22), 5.09 (m, 1H, H-3β), 4.12-4.03 (m, 2H, H-7aα, H-7aβ), 3.02 (dd, 1H, *J* = 12.2, 4.3 Hz, H-5α), 2.17 (ddd, 1H, *J* = 15.0, 12.5, 2.5 Hz), 2.10-1.96 (m, 3H), 2.07, 2.01, 2.00 (all s, 1H, CH_3_), 1.85-1.13 (m, 17H), 1.01 (d, 3H, *J* = 6.7 Hz, CH_3_), 0.96 (d, 3H, *J* = 6.7 Hz, CH_3_), 0.94 (d, 3H, *J* = 6.7 Hz, CH_3_), 0.91 (d, 3H, *J* = 6.7 Hz, CH_3_), 0.90 (s, 3H, CH_3_), 0.73 (s, 3H, CH_3_). ^13^C NMR (125 MHz, CDCl_3_): 176.04, 170.56, 170.51, 170.30, 75.67, 74.05, 70.29, 68.41, 58.37, 52.29, 51.42, 42.55, 42.46, 39.78, 39.56, 39.43, 36.97, 36.11, 33.67, 30.34, 29.73, 27.98, 25.14, 24.69, 22.12, 21.37, 20.93, 20.86 (2×C), 20.31, 14.61, 12.71, 11.59, 11.05. HRMS (APCI+) calculated for C_34_H_55_O_8_ ([M + H]^+^) 591.3897, found 591.3896.

Compound **12a** ((22*R,*23*R*)-7a-homo-6-oxa-7-oxo-5a-campestan-3α,22,23-triyl triacetate): ^1^H NMR (500 MHz, CDCl_3_): ‘6 5.33 (dd, 1H, *J* = 8.9, 1.5 Hz, H-23), 5.15 (d, 1H, *J* = 8.9 Hz, H-22), 5.14 (m, 1H, H-3β), 4.46 (dd, 1H, *J* = 11.2, 5.5 Hz, H-5α), 2.53 (d, 1H, *J* = 13.3 Hz, H-7aβ), 2.45 (dd, 1H, *J* = 13.3, 11.3 Hz, H-7aα), 2.12-1.95 (m, 3H), 2.07, 2.01, 2.00 (all s, 1H, CH_3_), 1.90-1.09 (m, 17H), 1.01 (d, 3H, *J* = 6.7 Hz, CH_3_), 0.96 (d, 3H, *J* = 6.7 Hz, CH_3_), 0.95 (d, 3H, *J* = 6.7 Hz, CH_3_), 0.914 (s, 3H, CH_3_), 0.908 (d, 3H, *J* = 6.7 Hz, CH_3_), 0.73 (s, 3H, CH_3_). ^13^C NMR (125 MHz, CDCl_3_): 174.74, 170.53 (2×C), 170.16, 79.67, 75.61, 74.03, 69.56, 58.02, 55.45, 52.62, 42.52, 39.79, 39.64, 39.58, 38.03, 36.97, 34.80, 32.93, 31.84, 30.36, 27.65, 25.14, 24.83, 22.18, 21.31, 20.94, 20.86 (2×C), 20.31, 12.70, 11.64, 11.59, 11.06. HRMS (APCI+) calculated for C_34_H_55_O_8_ ([M + H]^+^) 591.3897, found 591.3888.

**Scheme 2:**
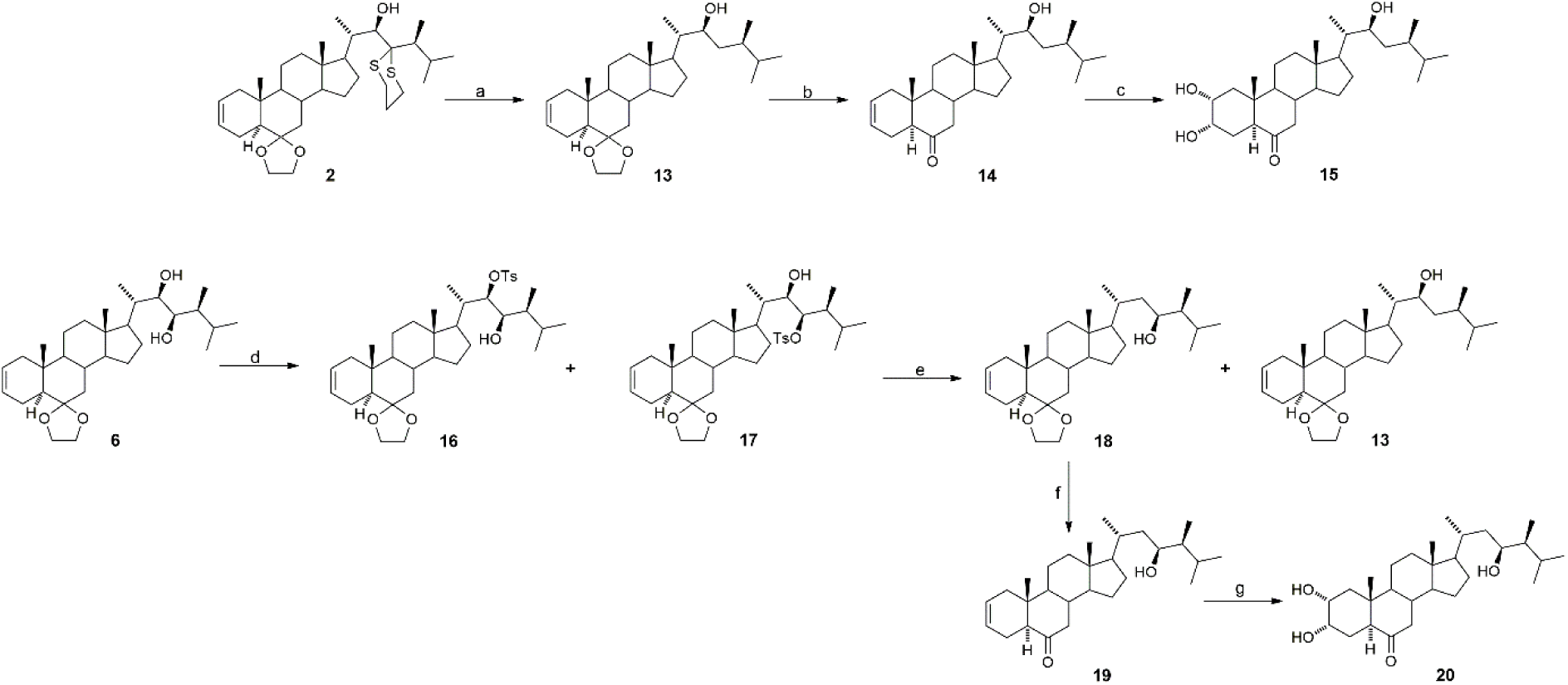
Synthesis of 23-deoxycastasterone (**15**), 22-deoxycastaterone (**20**). **a**: Ra-Ni/EtOH, r.t., 6h; **b**: HCl/H_2_O/THF, 40 °C, 4h; **c**: OsO_4_/K_2_CO_3_/K_2_[Fe(CN)_6_]/CH_3_SO_2_NH_2_/HQ-4-ClBz/t-BuOH/H_2_O, r.t., 10 h; **d**: *p*-TsCl/pyridine, r.t., 3 days; **e**: LiAlH_4_/THF/Et_2_O, r.t., 3h; **f**: HCl/H_2_O/THF, 40 °C, 2h; **g**: OsO_4_/K_2_CO_3_/K_2_[Fe(CN)_6_]/CH_3_SO_2_NH_2_/HQ-4-ClBz/t- BuOH/H_2_O, r.t., 10 h.

### Synthesis of (22*S*)-6,6-ethylenedioxy-5*a*-campest-2-en-22-ol (13)

Raney-Nickel catalyst 2400 (Merck, 1.4g) was added to a solution of compound **2** (200 mg, mmol) in ethanol (20 mL). The resulting suspension was shaken vigorously for 6 hours and then the liquid was decanted. Catalyst was washed with ethanol (2× 10 mL) and the combined ethanolic fractions were evaporated under reduced pressure. The residue was purified by column chromatography on silica gel (5-10% ethyl acetate in cyclohexane) to give alcohol **13** (85 mg, 52%) as a viscous oil. ^1^H NMR (500 MHz, CDCl_3_): ‘6 5.67 (m, 1H, H-3), 5.55 (m, 1H, H-2), 4.01-3.89 (m, 3H, H_ketal_), 3.81-3.75 (m, 2H, H-22, H_ketal_), 2.11-1.90 (m, 4H), 1.81-1.70 (m, 3H), 1.63-0.76 (m, 18H), 0.895 (d, 3H, *J* = 6.7 Hz, CH_3_), 0.889 (s, 3H, CH_3_), 0.880 (d, 3H, *J* = 6.7 Hz, CH_3_), 0.84 (d, 3H, *J* = 6.7 Hz, CH_3_), 0.82 (d, 3H, *J* = 6.7 Hz, CH_3_) 0.71 (s, 3H, CH_3_). ^13^C NMR (125 MHz, CDCl_3_): 125.67, 124.80, 110.05, 71.66, 65.59, 64.08, 55.89, 53.41, 52.58, 48.05, 42.46, 41.21 (2×C), 39.79, 39.35, 39.29, 35.87, 35.29, 33.40, 32.00, 27.71, 24.08, 21.42, 20.87, 19.98, 17.80, 15.76, 13.60, 11.92, 11.22. HRMS (ESI+) calculated for C_30_H_51_O_3_ ([M + H]^+^) 459.3838, found 459.3834.

### Synthesis of (22*S*)-22-hydroxy-5*a*-campest-2-en-6-one (14)

Aqueous solution of HCl (5%, 0.5 mL) was added to a solution of compound **13** (70 mg, 0.15 mmol) in THF (5 mL) and the reaction mixture was stirred at 40 °C for 4 hours and then diluted with ethyl acetate. Organic layer was washed twice with water, dried over MgSO_4_, filtrated and evaporated under reduced pressure. The residue was purified by column chromatography on silica gel (10% ethyl acetate in cyclohexane) to give compound **14** (55 mg, 87%) as a white solid. ^1^H NMR (500 MHz, CDCl_3_): ‘6 5.68 (m, 1H, H-3), 5.56 (m, 1H, H-2), 3.67 (td, *J* = 6.8, 1.1 Hz, H-22), 2.37-2.34 (m, 2H), 2.25 (m, 1H), 2.06-1.92 (m, 4H), 1.74 (m, 1H), 1.62-0.96 (m, 17H), 0.89 (d, 3H, *J* = 6.7 Hz, CH_3_), 0.87 (d, 3H, *J* = 6.7 Hz, CH_3_), 0.82 (d, 3H, *J* = 6.7 Hz, CH_3_), 0.80 (d, 3H, *J* = 7.0 Hz, CH_3_), 0.71 (s, 3H, CH_3_), 0.68 (s, 3H, CH_3_). ^13^C NMR (125 MHz, CDCl_3_): 212.06, 124.87, 124.46, 71.48, 56.59, 53.74, 53.27, 52.39, 46.92, 42.66, 39.99, 39.41, 39.31, 39.26, 39.22, 37.70, 35.22, 31.96, 27.57, 23.83, 21.64, 21.07, 19.92, 17.77, 15.71, 13.45, 11.82, 11.16. HRMS (ESI+) calculated for C_28_H_47_O_2_ ([M + H]^+^) 415.3576, found 415.3578.

### Synthesis of (22*S*)-2**α**,3**α**,22-trihydroxy-5*a*-campestan-6-one (23-deoxycastasterone, 15)

Solution of osmium tetroxide in *t*-butanol (0.2M, 0.15 mL, 0.03 mmol) was added to a solution of compound **14** (40 mg, 0.1 mmol), hydroquinidine 4-chlorobenzoate (25 mg; 0.05 mmol), methanesulfonamide (5 mg; 0.05 mmol), potassium carbonate (100 mg; 0.72 mmol), and potassium ferricyanide (240 mg; 0.73 mmol) in the mixture of *t*-butanol and water (5 mL; 1 : 1 v/v). The reaction mixture was stirred at room temperature for 10 h and quenched with saturated solution of Na_2_SO_3_ (3 mL) was then added. After an additional 30 minutes of stirring, the reaction mixture was diluted with ethyl acetate (30 mL) and extracted with water (2 × 20 mL). Organic layer was dried over MgSO_4_, filtrated and evaporated under reduced pressure. The residue was purified by column chromatography on silica gel (0.5% isopropanol in ethyl acetate) to give triol **15** (31 mg, 72%) as a white solid. ^1^H NMR (500 MHz, CDCl_3_ + 5% CD_3_OD): 3.95 (m, 1H, H-3β), 3.73 (m, 1H, H-2β), 3.71 (bt, 1H, *J* = 6.4 Hz, H-22), 2.65 (dd, 1H, *J* = 12.4, 2.3 Hz, H-5α), 2.25 (dd, 1H, *J* = 13.0, 4.6 Hz, H-7β), 2.01-1.21 (m, 21H), 1.06 (m, 1H), 0.85 (d, 3H, *J* = 6.7 Hz, CH_3_), 0.83 (d, 3H, *J* = 6.7 Hz, CH_3_), 0.78 (d, 3H, *J* = 7.0 Hz, CH_3_), 0.77 (d, 3H, *J* = 7.0 Hz, CH_3_), 0.71 (s, 3H, CH_3_), 0.64 (s, 3H, CH_3_). ^13^C NMR (125 MHz, CDCl_3_ + 5% CD_3_OD): 213.04, 71.30, 67.97, 67.88, 56.48, 53.57, 52.27, 50.71, 46.62, 42.75, 42.54, 39.84, 39.31, 39.13, 39.08, 37.70, 35.14, 31.90, 29.57, 27.50, 23.77, 21.09, 19.84, 17.68, 15.59, 13.42, 11.81, 11.08. HRMS (ESI+) calculated for C_28_H_49_O_4_ ([M + H]^+^) 449.3631, found 449.3633.

### Synthesis of (22*R,*23*R*)-6,6-ethylenedioxy-23-hydroxy-5*a*-campest-2-en-22-yl *p*-toluenesulfonate (16) and (22*R,*23*R*)-6,6-ethylenedioxy-22-hydroxy-5*a*-campest-2-en- 23-yl *p*-toluenesulfonate (17)

*p*-Toluenesulfonyl chloride (200 mg, 1.05 mmol) was added to a solution of compound **6** (100 mg; 0.21 mmol) in pyridine (4 mL). The reaction mixture was stirred at room temperature for 3 days. Then, the mixture was diluted with ethyl acetate and aqueous solution of HCl (5%, 5 mL) was added. The aqueous layer was separated and extracted with ethyl acetate (2×). The combined organic layers were washed with water, dried over MgSO_4_, filtrated and evaporated under reduced pressure. The residue was purified by column chromatography on silica gel (10% ethyl acetate in cyclohexane). This chromatography did not lead to separation of both tosylates and they were used in the next reaction step as a mixture of isomers. Yield of both isomers 91 mg (69%). Analytical sample was used for HPLC separation (8% ethyl acetate in cyclohexane) to give two tosylates, **16** and **17,** as a viscous oil.

Compound **16**: ^1^H NMR (500 MHz, CDCl_3_): 7.85 (d, 2H, *J* = 8.4 Hz, H_Ts_), 7.34 (d, 2H, *J* = 8.4 Hz, H_Ts_), 5.67 (m, 1H, H-3), 5.55 (m, 1H, H-2), 4.77 (d, 1H, *J* = 8.6 Hz, H-22), 4.01-3.91 (m, 4H, H-23, H_ketal_),3.80 (m, 1H, H_ketal_), 2.46 (s, 3H, CH_3_), 2.31 (d, 1H, *J* = 4.9 Hz, OH), 2.13-0.68 (m, 22H), 0.95 (d, 3H, *J* = 6.7 Hz, CH_3_), 0.94 (d, 3H, *J* = 6.7 Hz, CH_3_), 0.90 (d, 3H, *J* = 7.0 Hz, CH_3_), 0.88 (d, 3H, *J* = 7.0 Hz, CH_3_), 0.87 (s, 3H, CH_3_), 0.66 (s, 3H, CH_3_). ^13^C NMR (125 MHz, CDCl_3_): 144.50, 134.67, 129.61 (2×C), 127.62 (2×C), 125.72, 124.69, 109.92, 91.04, 71.41, 65.60, 64.12, 55.87, 53.52, 51.87, 48.08, 42.24, 41.26, 41.19, 40.88, 39.66, 37.72, 35.84, 33.27, 30.59, 27.95, 23.95, 21.70, 21.41, 20.84, 20.76, 20.70, 13.61, 12.57, 11.61, 10.17. HRMS (APCI+) calculated for C_37_H_57_O_6_S ([M + H]^+^) 629.3876, found 629.3865.

Compound **17**: ^1^H NMR (500 MHz, CDCl_3_): 7.83 (d, 2H, *J* = 8.3 Hz, H_Ts_), 7.34 (d, 2H, *J* = 8.3 Hz, H_Ts_), 5.66 (m, 1H, H-3), 5.55 (m, 1H, H-2), 5.04 (d, 1H, *J* = 8.6 Hz, H-22), 4.00-3.89 (m, 3H, H_ketal_), 3.81-3.75 (m, 2H, H-22, H_ketal_), 2.45 (s, 4H, OH, CH_3_), 2.13-0.76 (m, 22H), 1.00 (d, 3H, *J* = 6.7 Hz, CH_3_), 0.94 (d, 3H, *J* = 6.7 Hz, CH_3_), 0.888 (s, 3H, CH_3_), 0.8875 (d, 3H, *J* = 7.0 Hz, CH_3_), 0.84 (d, 3H, *J* = 7.0 Hz, CH_3_), 0.70 (s, 3H, CH_3_). ^13^C NMR (125 MHz, CDCl_3_): 144.50, 134.44, 129.65 (2×C), 127.58 (2×C), 125.64, 124.81, 110.03, 89.23, 72.97, 65.59, 64.09, 55.79, 53.32, 52.06, 48.03, 42.29, 41.18, 41.16, 40.57, 39.73, 37.32, 35.86, 33.41, 29.76, 27.65, 23.95, 21.65, 21.42, 21.14, 20.86, 20.66, 13.60, 11.96, 11.77, 11.15. HRMS (APCI+) calculated for C_37_H_57_O_6_S ([M + H]^+^) 629.3876, found 629.3868.

### Synthesis of (23*S*)-6,6-ethylenedioxy-5*a*-campest-2-en-23-ol (18) and (22*S*)-6,6- ethylenedioxy-5*a*-campest-2-en-22-ol (13)

1M Solution of LiAlH_4_ in diethylether (1 mL, 1.0 mmol) was added dropwise to a solution of compounds **16** and **17** (50 mg, 0.08 mmol) in dry THF (5 mL). The reaction mixture was stirred at room temperature for 3 hours and then quenched with MeOH (3 mL). Saturated aqueous solution of sodium potassium tartrate (10 mL) was added and the mixture was diluted with ethyl acetate. The aqueous layer was separated and extracted with ethyl acetate (2×). The combined organic layers were washed with water, dried over MgSO_4_, filtrated and evaporated under reduced pressure. The residue was purified by column chromatography on silica gel (5-10% ethyl acetate in cyclohexane) to give crude mixture of compounds **13** and **18** (32 mg). Both compounds were separated by HPLC (8% ethyl acetate in cyclohexane) to afford two alcohols **13** (11 mg, 31%) and **18** (14 mg, 39%) as a viscous oil. The spectroscopic data for compound **13** were entirely consistent with those reported above.

Compound **18**: ^1^H NMR (500 MHz, CDCl_3_): 5.67 (m, 1H, H-3), 5.55 (m, 1H, H-2), 4.01-3.88 (m, 4H, H-23, H_ketal_), 3.79 (q, 1H, *J* = 6.6 Hz, H_ketal_), 2.12-1.86 (m, 5H), 1.80-0.75 (m, 20H), 0.98 (d, 3H, *J* = 6.7 Hz, CH_3_), 0.96 (d, 3H, *J* = 6.7 Hz, CH_3_), 0.93 (d, 3H, *J* = 6.7 Hz, CH_3_), 0.89 (s, 3H, CH_3_), 0.84 (d, 3H, *J* = 7.0 Hz, CH_3_), 0.70 (s, 3H, CH_3_). ^13^C NMR (125 MHz, CDCl_3_): 125.68, 124.81, 110.05, 70.98, 65.58, 64.09, 57.00, 55.93, 53.42, 48.07, 42.61, 42.45, 42.06, 41.22, 41.20, 39.80, 35.88, 33.97, 33.33, 30.43, 28.40, 24.13, 21.43, 21.11, 20.85, 20.49, 19.21, 13.60, 11.97, 9.27. HRMS (APCI+) calculated for C_30_H_51_O_3_ ([M + H]^+^) 459.3838, found 459.3831.

### Synthesis of (23*S*)-23-hydroxy-5*a*-campest-2-en-6-one (19)

Aqueous solution of HCl (5%, 0.5 mL) was added to a solution of compound **18** (12 mg, 0.03 mmol) in THF (3 mL) and the reaction mixture was stirred at 40 °C for 2 hours and then diluted with ethyl acetate. Organic layer was washed twice with water, dried over MgSO_4_, filtrated and evaporated under reduced pressure. The residue was purified by column chromatography on silica gel (10% ethyl acetate in cyclohexane) to give compound **19** (9 mg, 83%) as a viscous oil. ^1^H NMR (500 MHz, CDCl_3_): 5.69 (m, 1H, H-3), 5.57 (m, 1H, H-2), 3.90 (ddd, 1H, *J* = 7.9, 5.7, 2.6 Hz, H-23), 2.38-2.34 (m, 2H), 2.27 (m, 1H), 2.09-1.89 (m, 5H), 1.75 (m, 1H), 1.66-1.04 (m, 16H), 0.99 (d, 3H, *J* = 6.7 Hz, CH_3_), 0.96 (d, 3H, *J* = 6.7 Hz, CH_3_), 0.93 (d, 3H, *J* = 6.7 Hz, CH_3_), 0.84 (d, 3H, *J* = 7.0 Hz, CH_3_), 0.72 (s, 3H, CH_3_), 0.69 (s, 3H, CH_3_). ^13^C NMR (125 MHz, CDCl_3_): 212.09, 124.94, 124.51, 70.97, 58.87, 58.67, 53.81, 53.34, 46.97, 42.87, 42.49, 41.98, 40.04, 39.50, 39.32, 37.67, 33.93, 30.41, 28.31, 23.93, 21.70, 21.11, 21.09, 20.48, 19.18, 13.50, 11.91, 9.26. HRMS (APCI+) calculated for C_28_H_47_O_2_ ([M + H]^+^) 415.3576, found 415.3569.

### Synthesis of (23*S*)-2**α**,3**α**,23-trihydroxy-5*a*-campestan-6-one (22-deoxycastasterone, 20)

Solution of osmium tetroxide in *t*-butanol (0.2M, 0.05 mL, 0.01 mmol) was added to a solution of compound **19** (8 mg, 0.02 mmol), hydroquinidine 4-chlorobenzoate (10 mg; 0.02 mmol), methanesulfonamide (3 mg; 0.03 mmol), potassium carbonate (20 mg; 0.14 mmol), and potassium ferricyanide (45 mg; 0.14 mmol) in the mixture of *t*-butanol and water (3 mL; 1: 1 v/v). The reaction mixture was stirred at room temperature for 10 h and quenched with saturated solution of Na_2_SO_3_ (2 mL) was then added. After an additional 30 minutes of stirring, the reaction mixture was diluted with ethyl acetate (20 mL) and extracted with water (2 × 20 mL). Organic layer was dried over MgSO_4_, filtrated and evaporated under reduced pressure. The residue was purified by HPLC (70% ethyl acetate in cyclohexane) to give triol **20** (6 mg, 69%) as a white solid (lyophilized from *t*-butanol). ^1^H NMR (500 MHz, CDCl_3_ + 10% CD_3_OD): 4.03 (m, 1H, H-3β), 3.89 (ddd, 1H, *J* = 8.0, 5.7, 2.6 Hz, H-23), 3.75 (m, 1H, H-2β), 2.68 (dd, 1H, *J*= 12.2, 3.4 Hz, H-5α), 2.29 (dd, 1H, *J* = 13.1, 4.6 Hz, H-7β), 2.07-1.05 (m, 22H), 0.97 (d, 3H, *J* = 6.7 Hz, CH_3_), 0.95 (d, 3H, *J* = 6.7 Hz, CH_3_), 0.92 (d, 3H, *J* = 6.7 Hz, CH_3_), 0.83 (d, 3H, *J* = 6.7 Hz, CH_3_), 0.75 (s, 3H, CH_3_), 0.67 (s, 3H, CH_3_). ^13^C NMR (125 MHz, CDCl_3_ + 10% CD_3_OD): 212.38, 70.94, 68.33, 68.20, 56.79, 56.57, 53.62, 50.68, 46.72, 42.98, 42.58, 42.43, 41.89, 40.12, 39.37, 37.65, 33.90, 30.38, 28.29, 26.25, 23.90, 21.14, 21.09, 20.47, 19.11, 13.52, 11.96, 9.24. HRMS (APCI+) calculated for C_28_H_49_O_4_ ([M + H]^+^) 449.3631, found 449.3629.

---

**^1^H and ^13^C NMR spectra of compounds prepared**

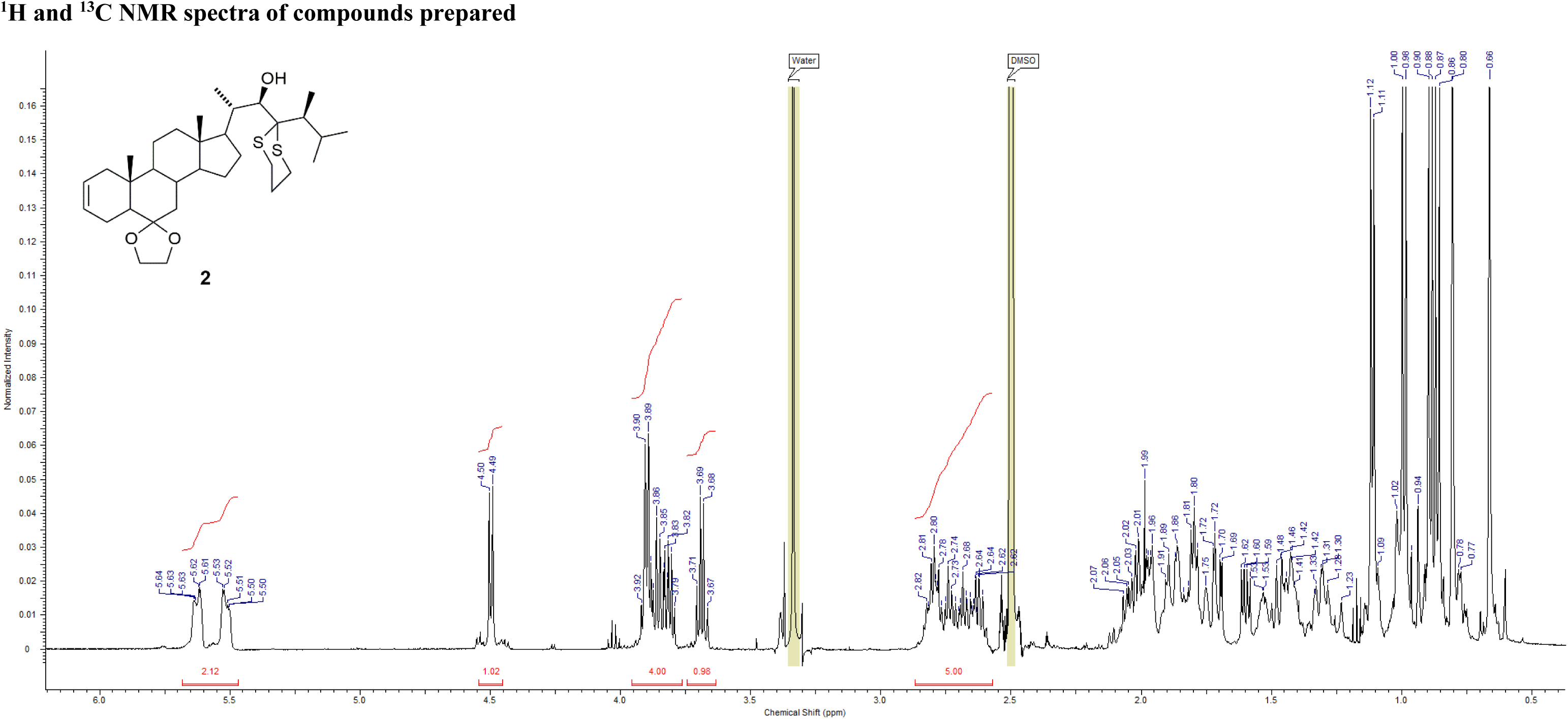
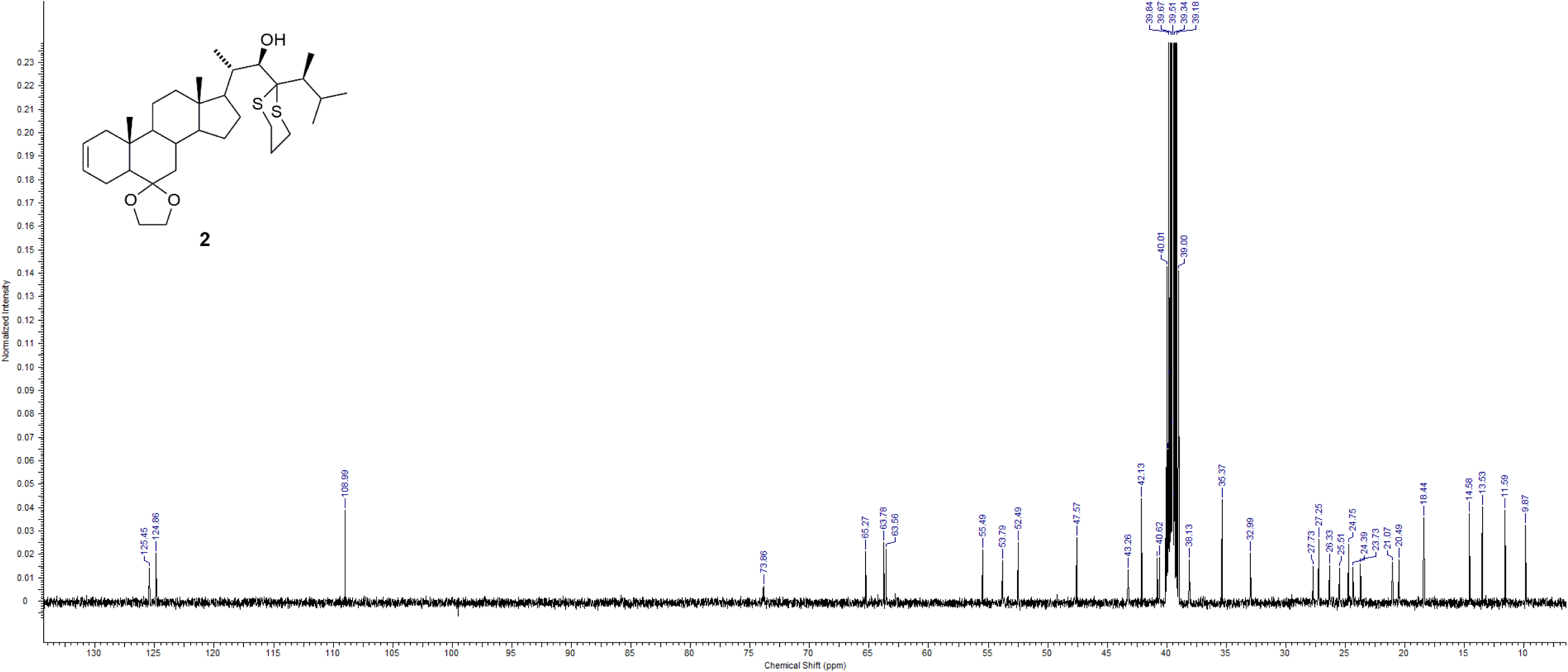
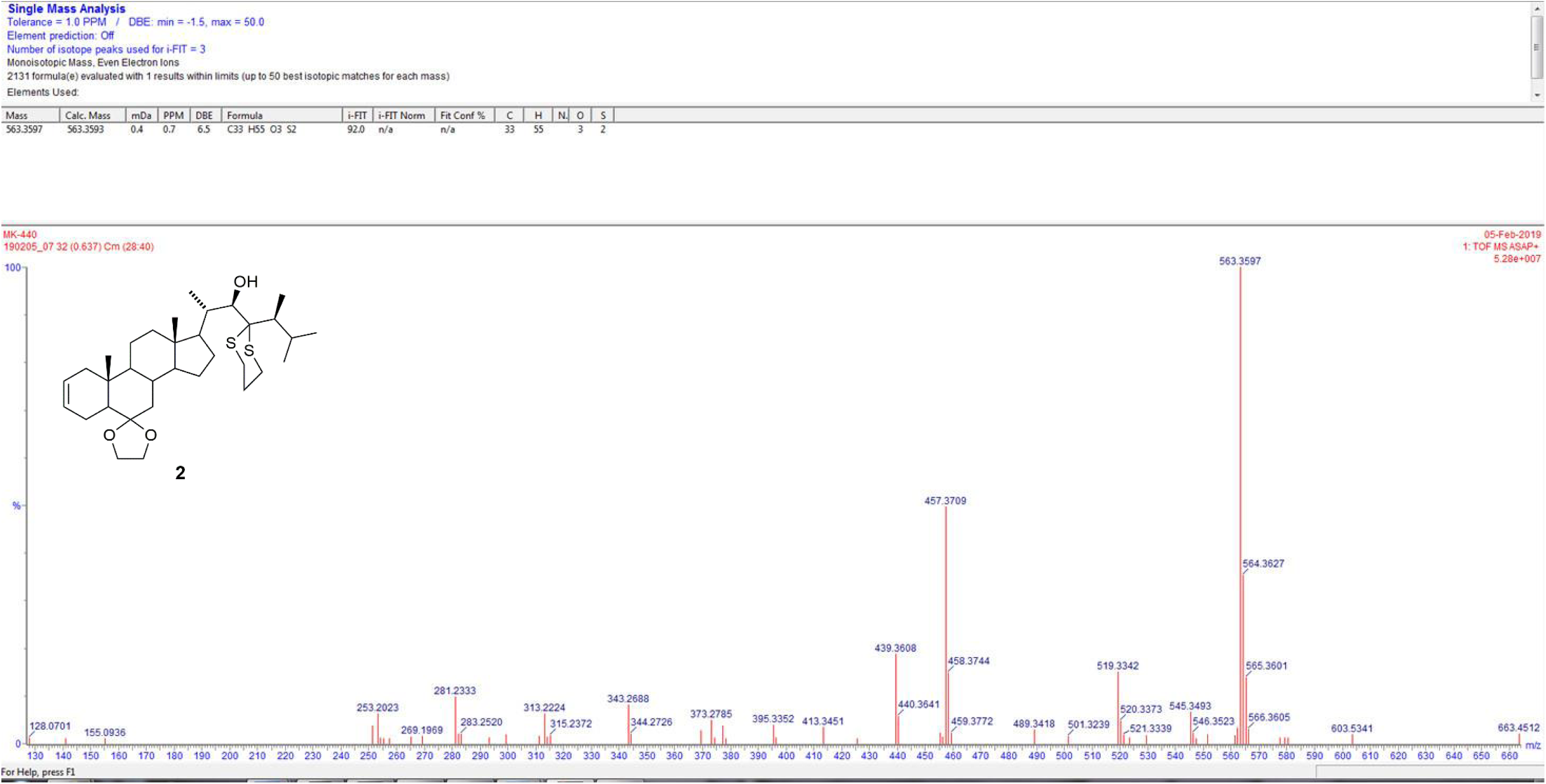
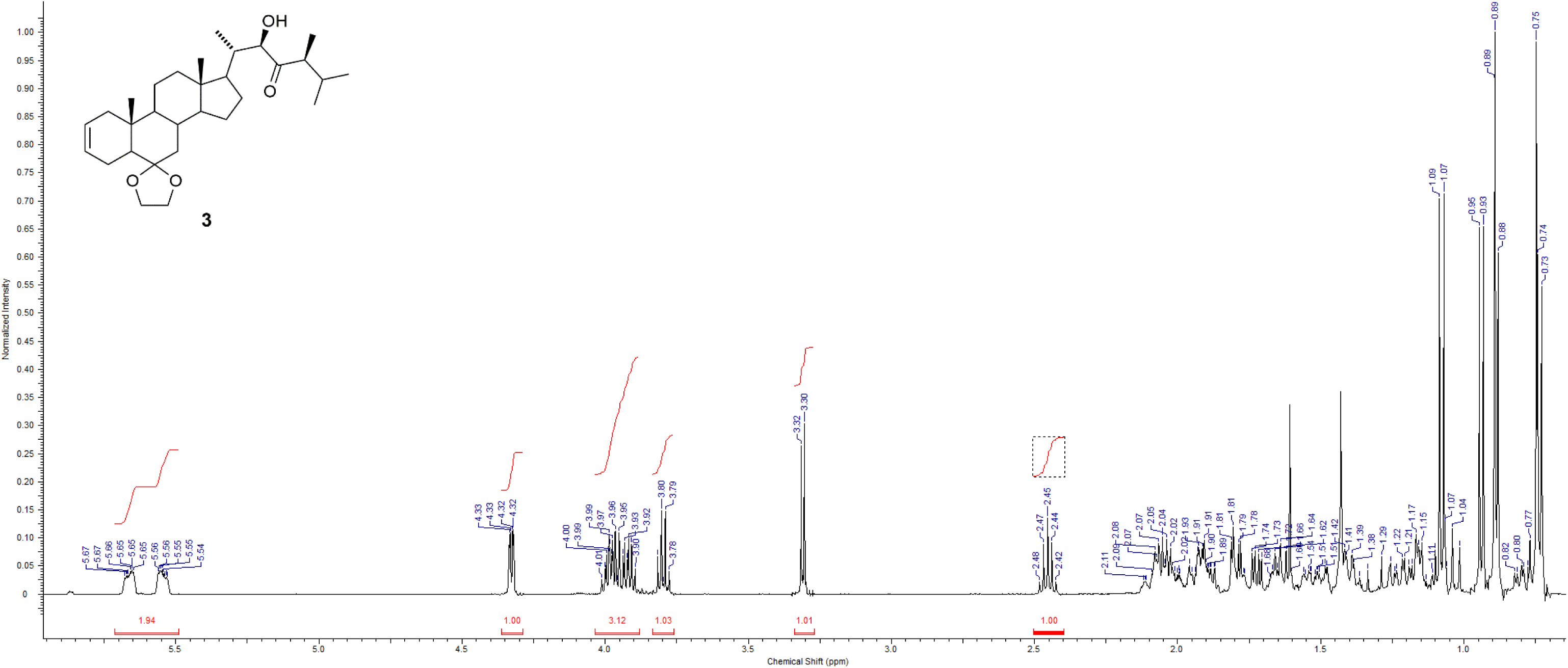
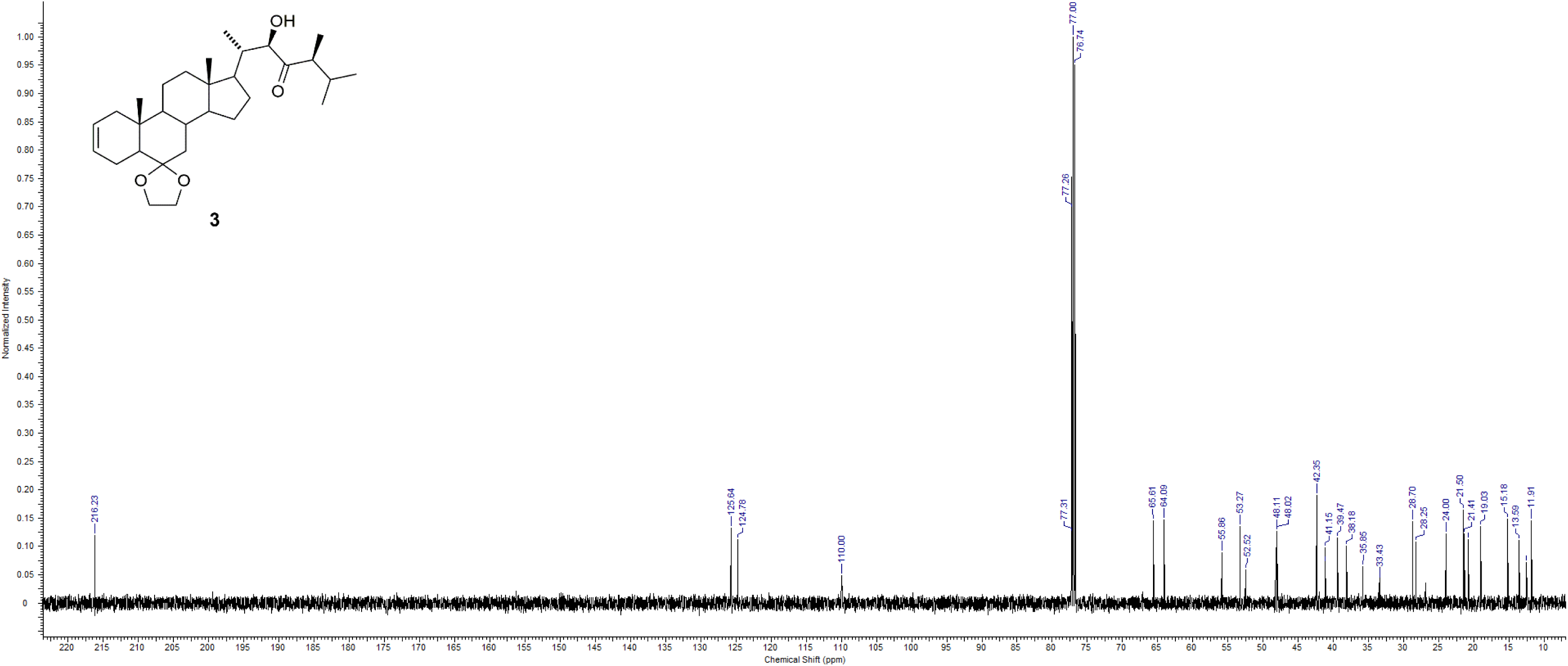
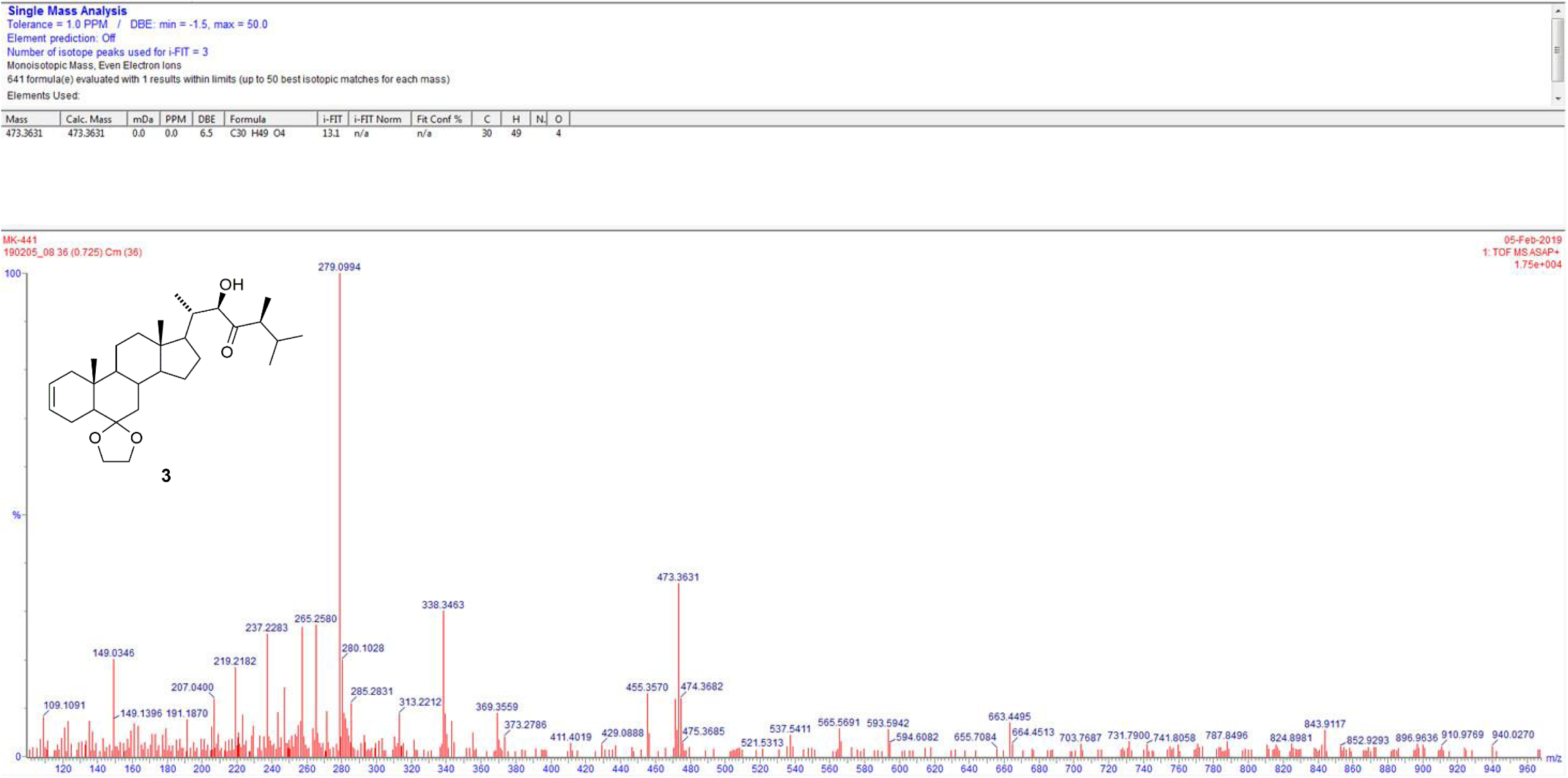
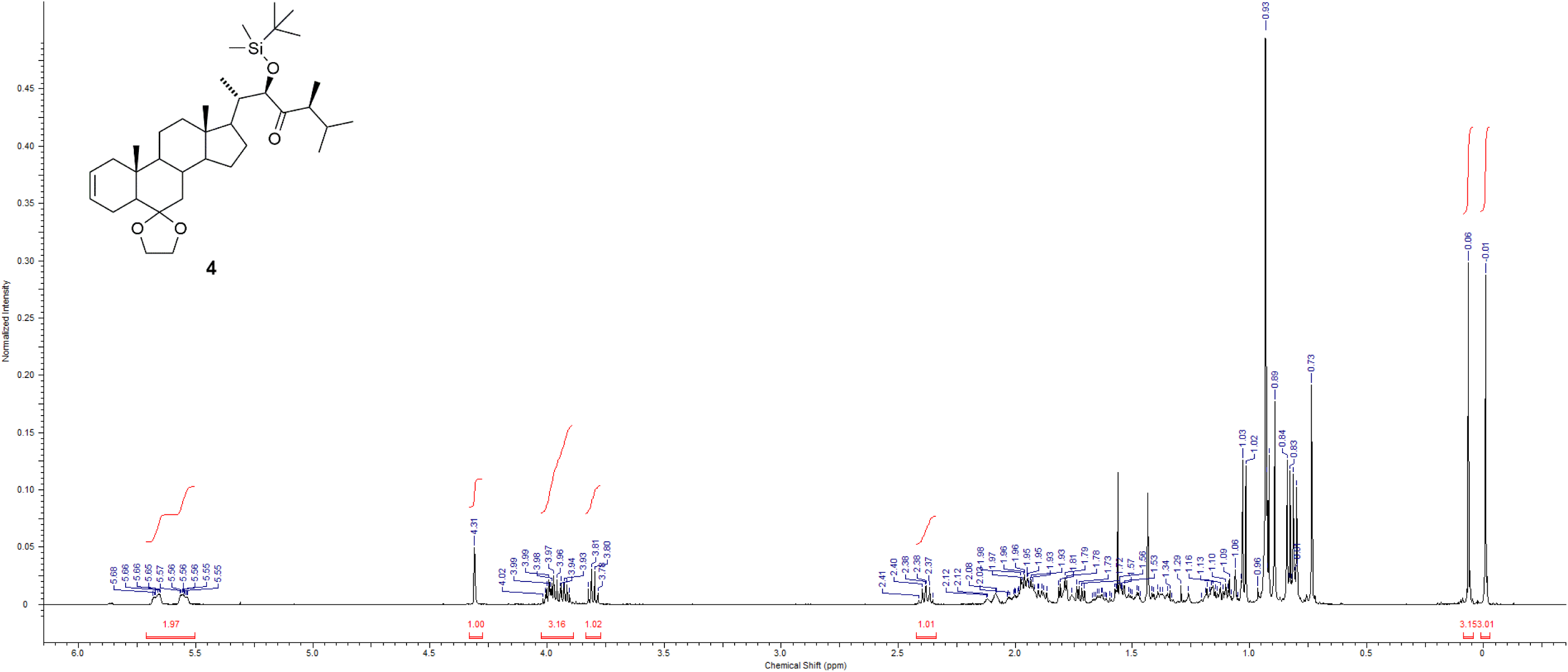
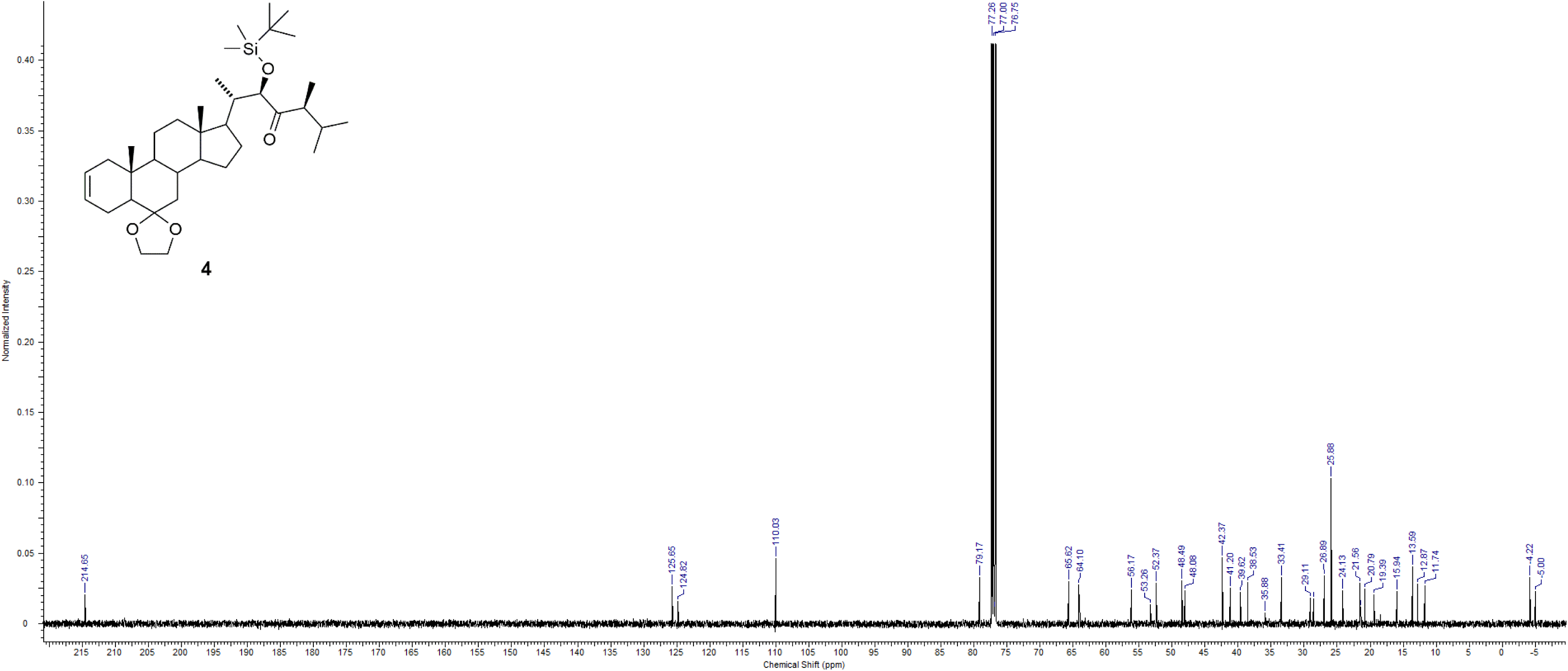
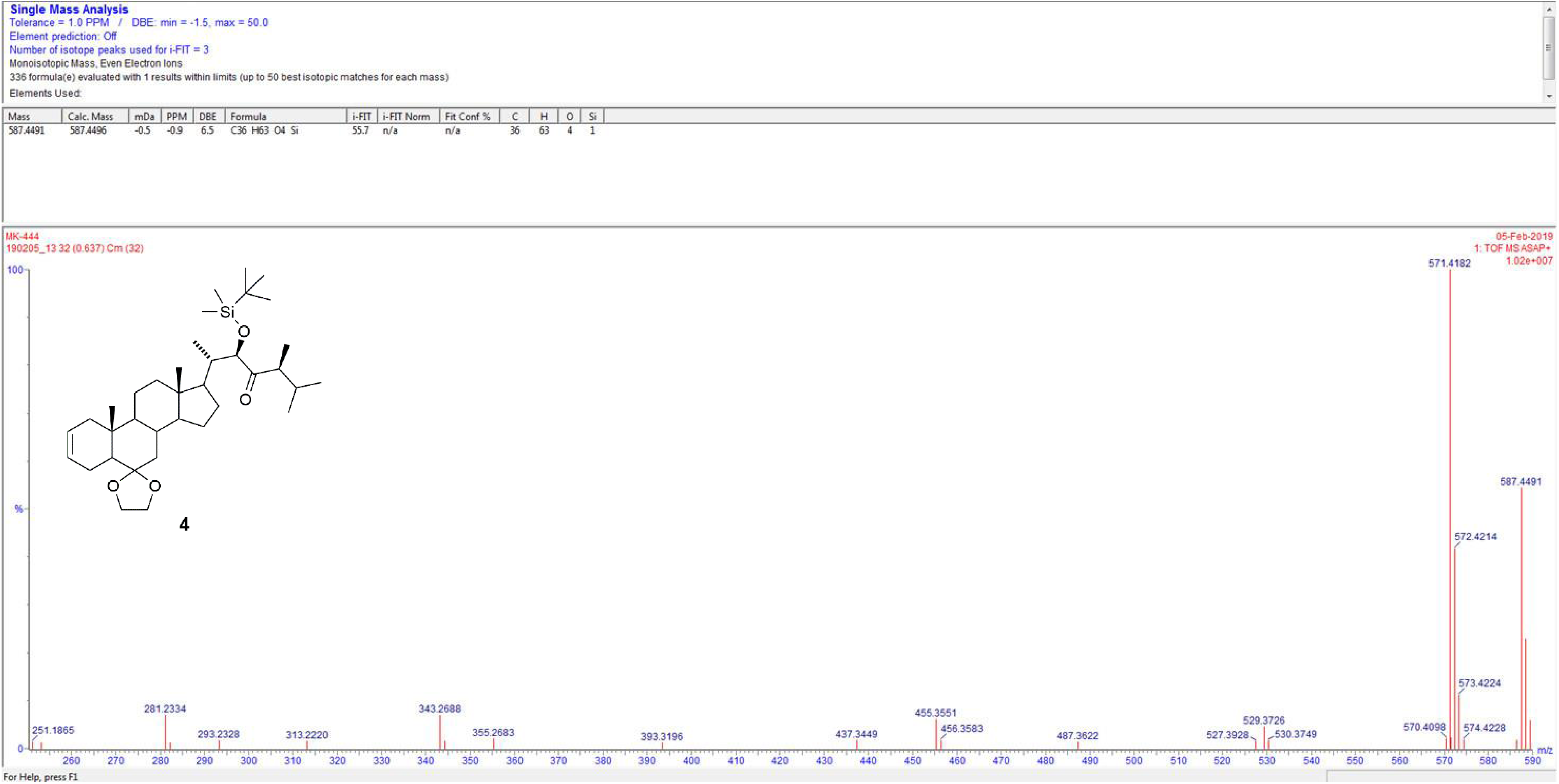
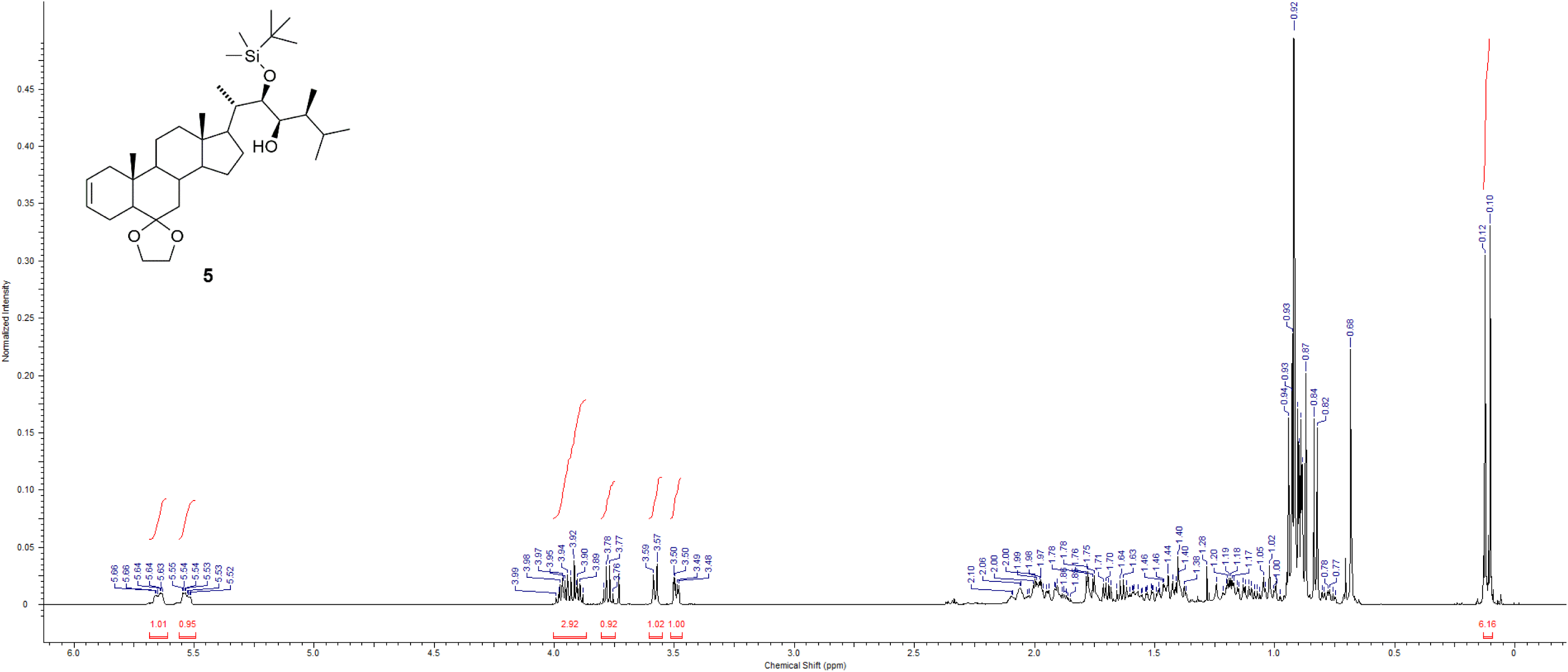
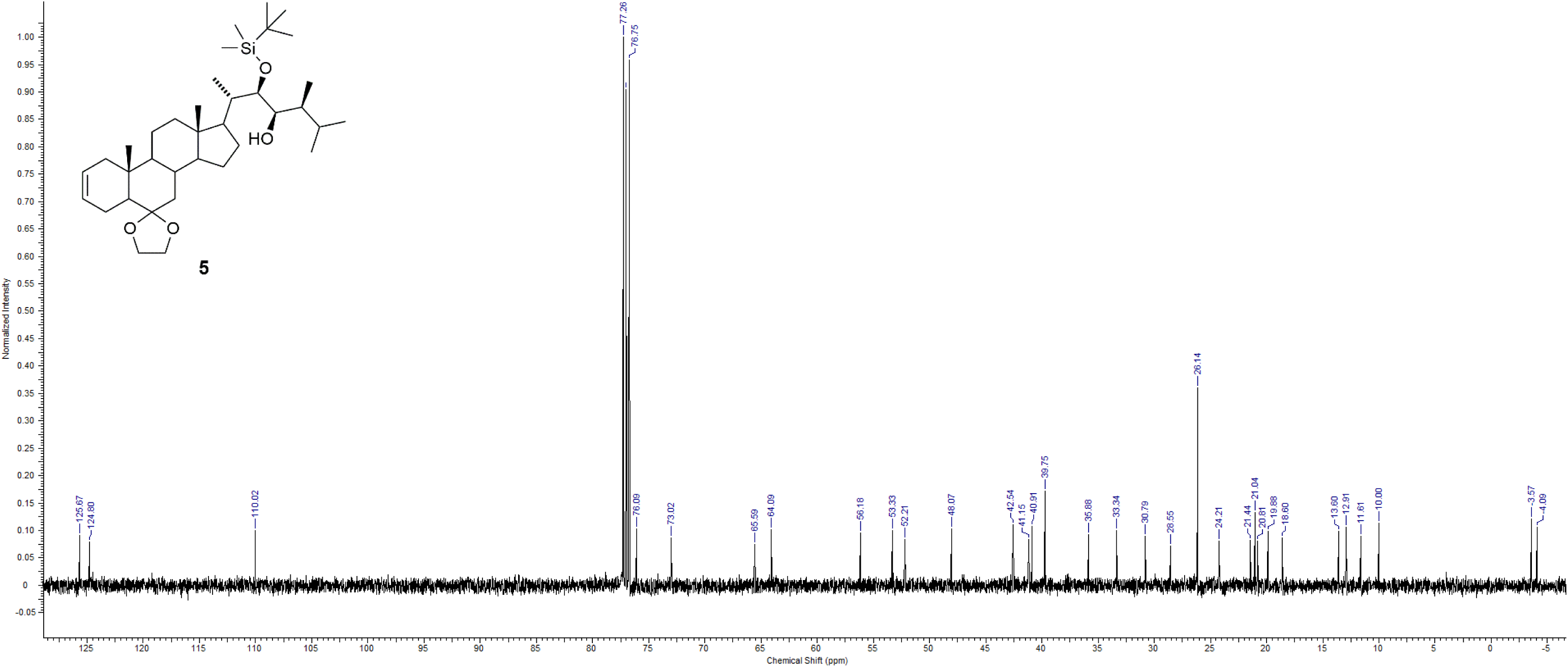
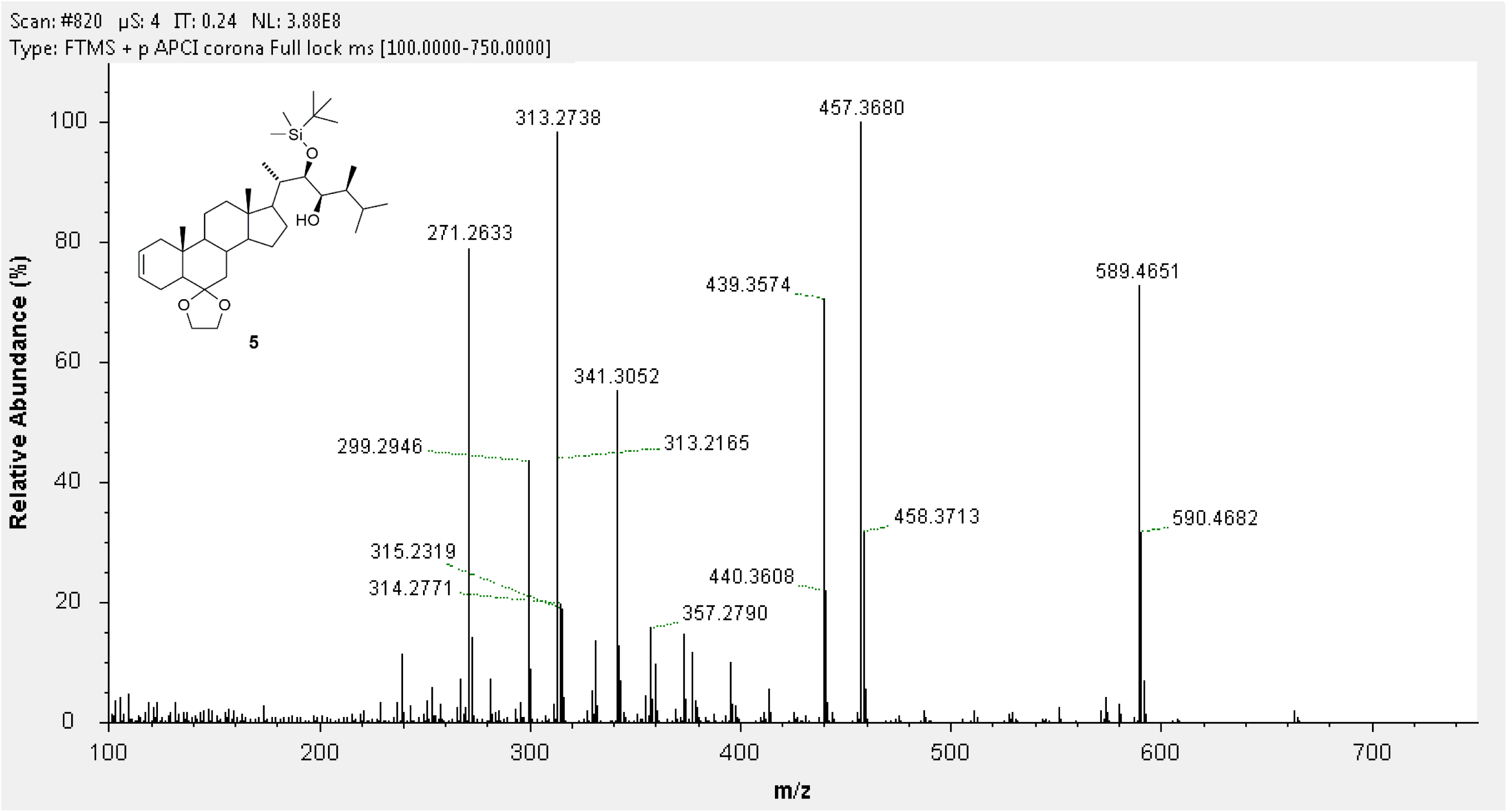
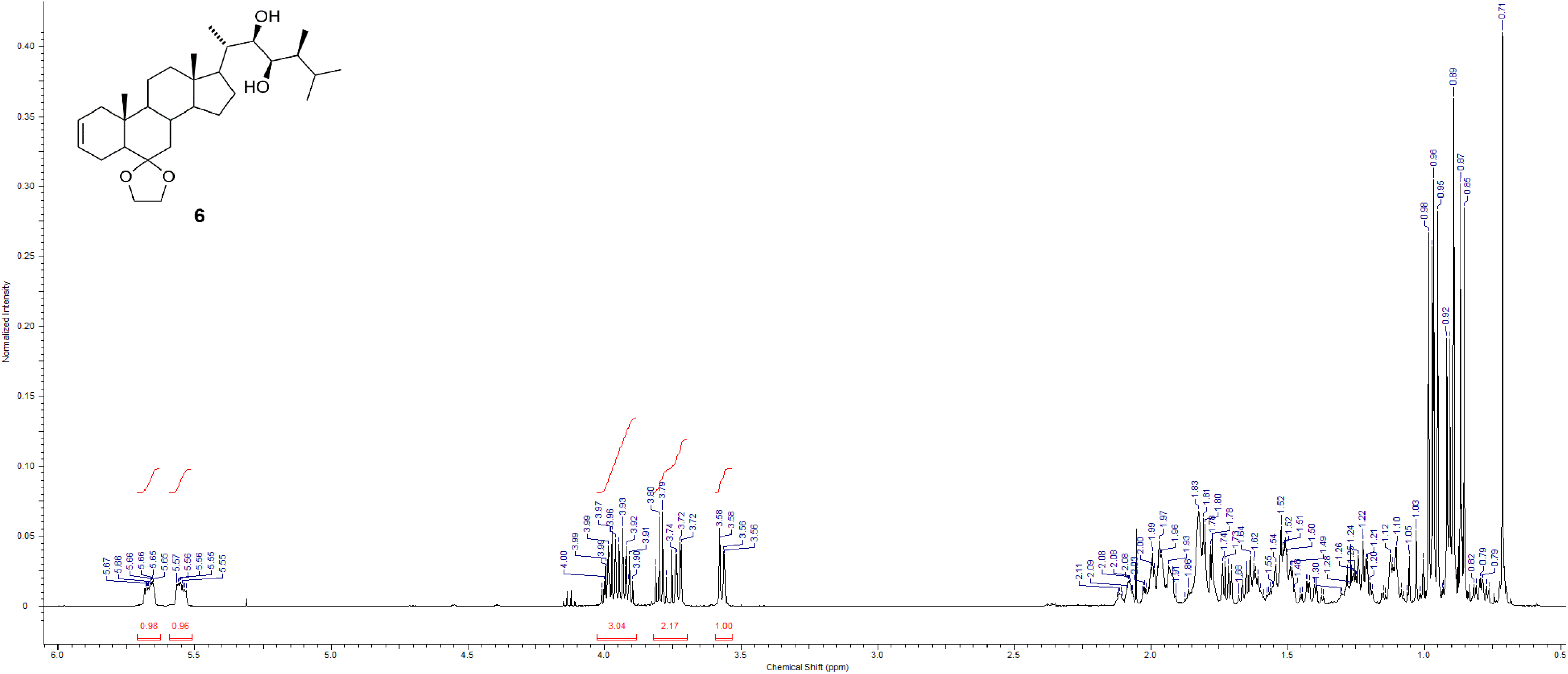
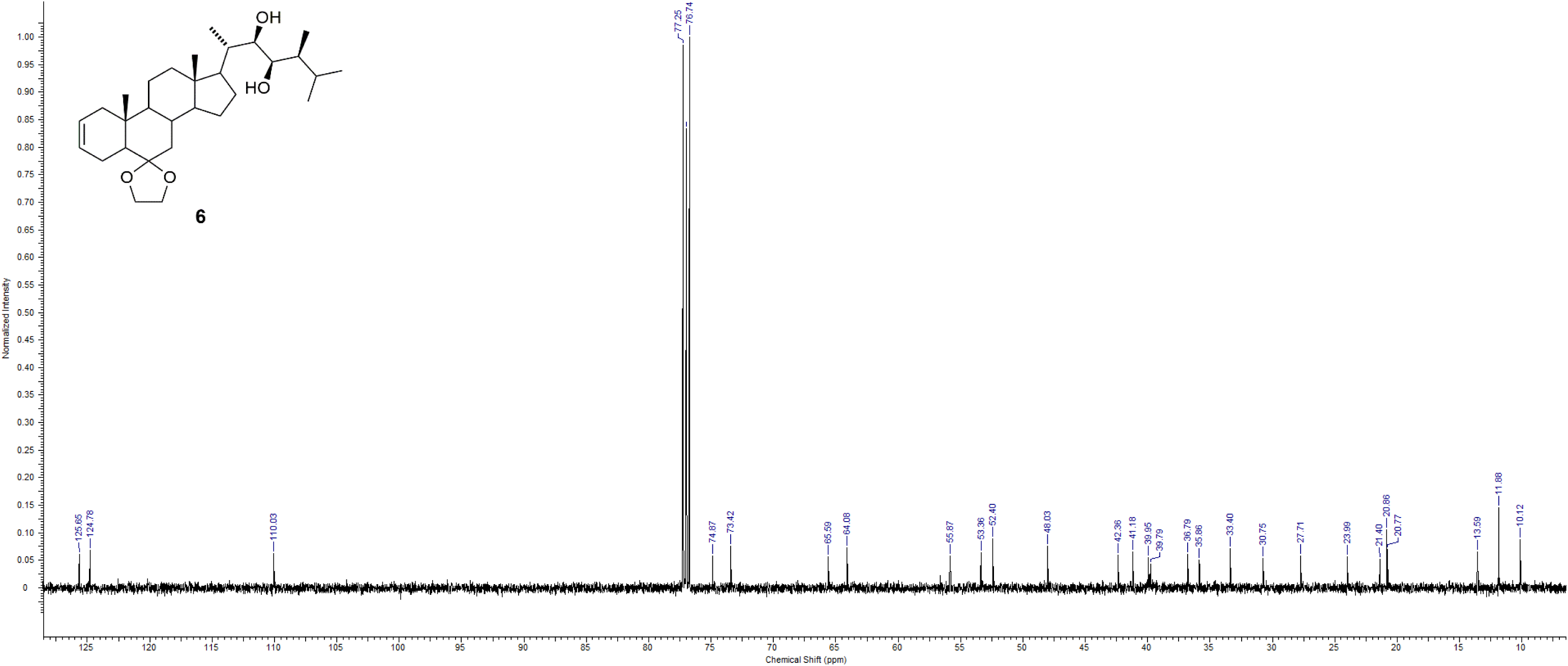
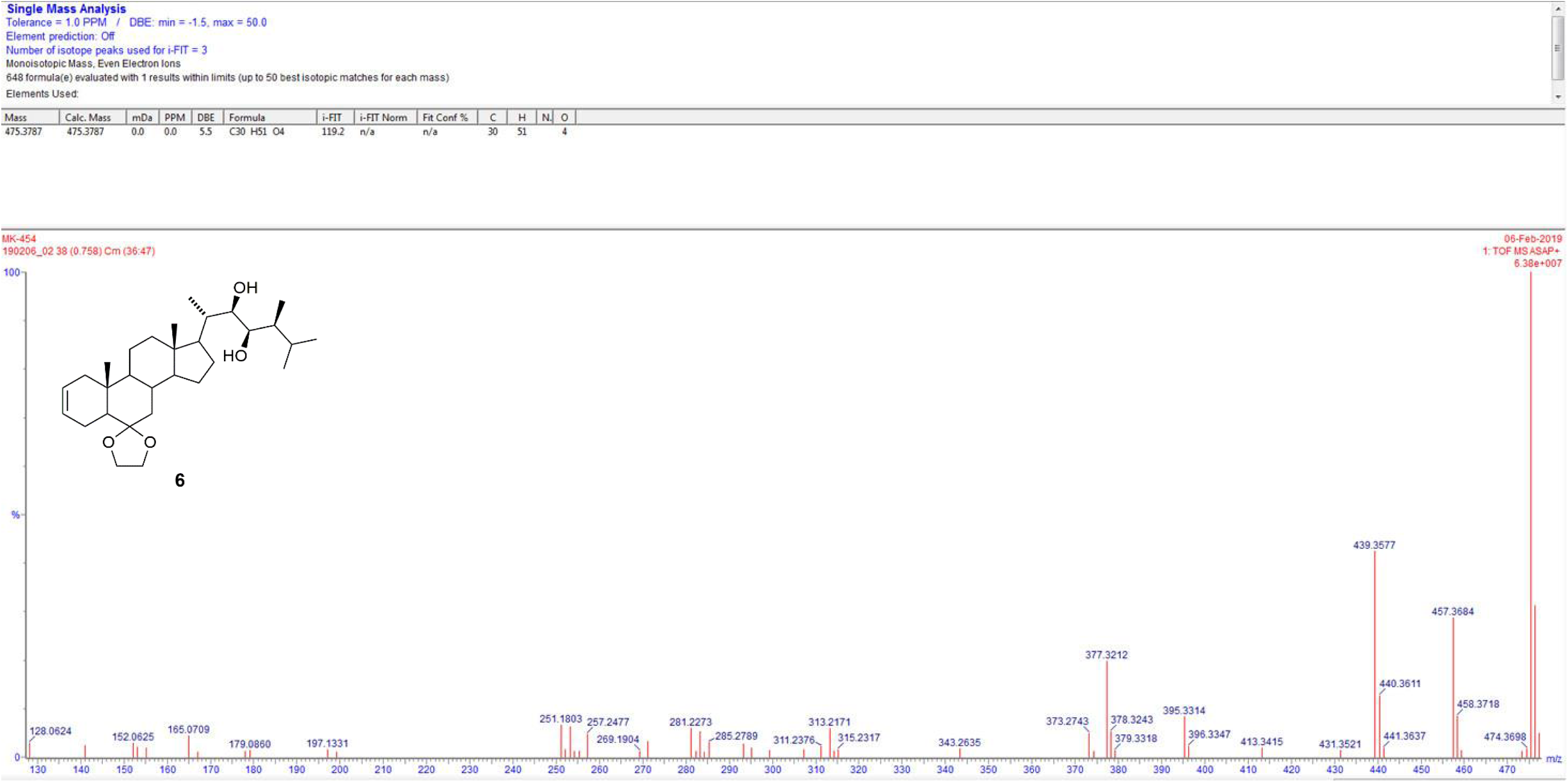
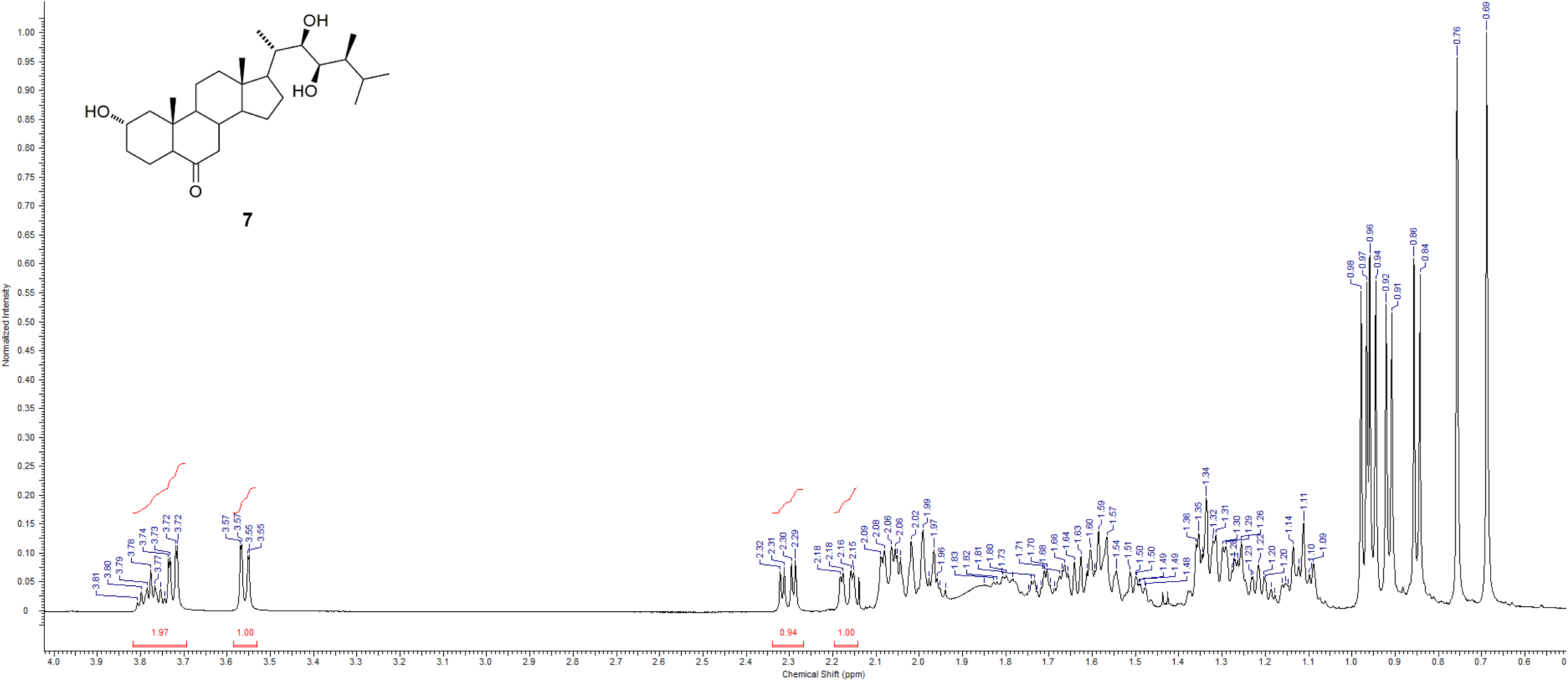
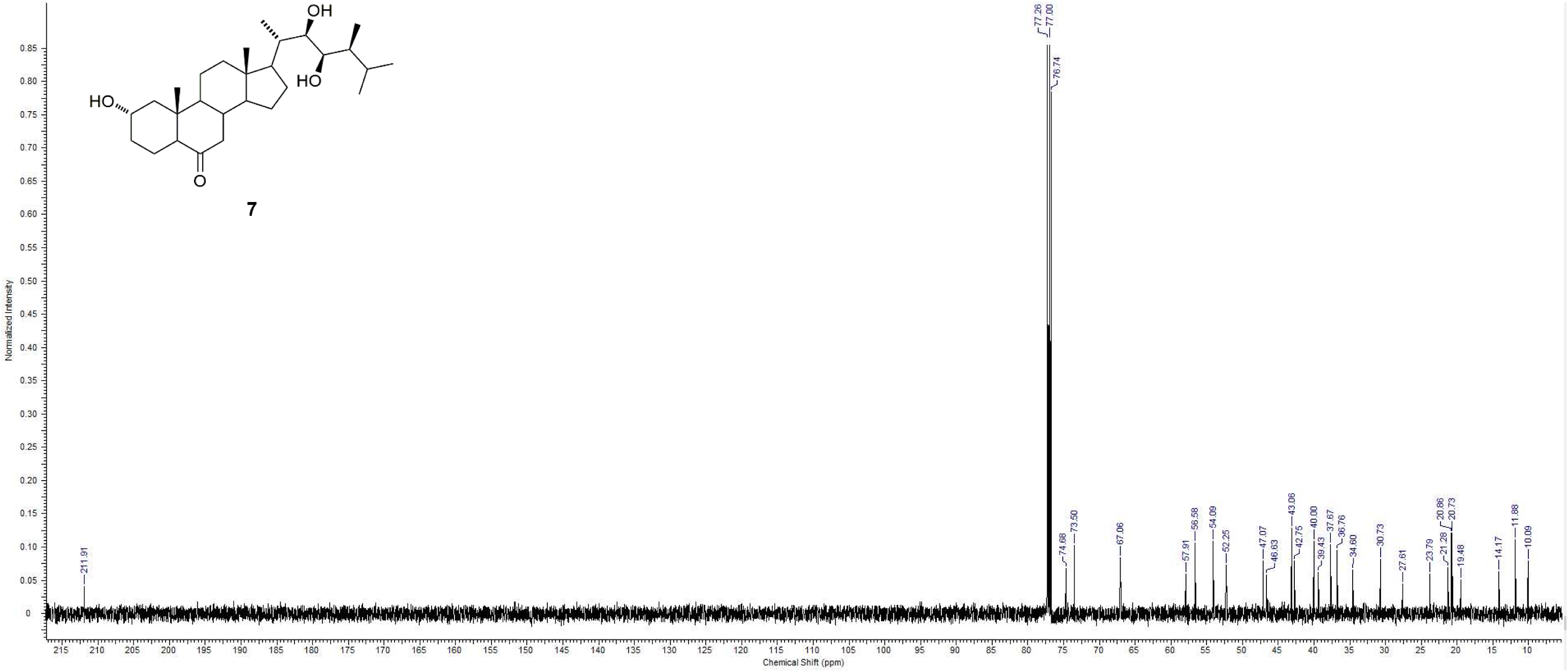
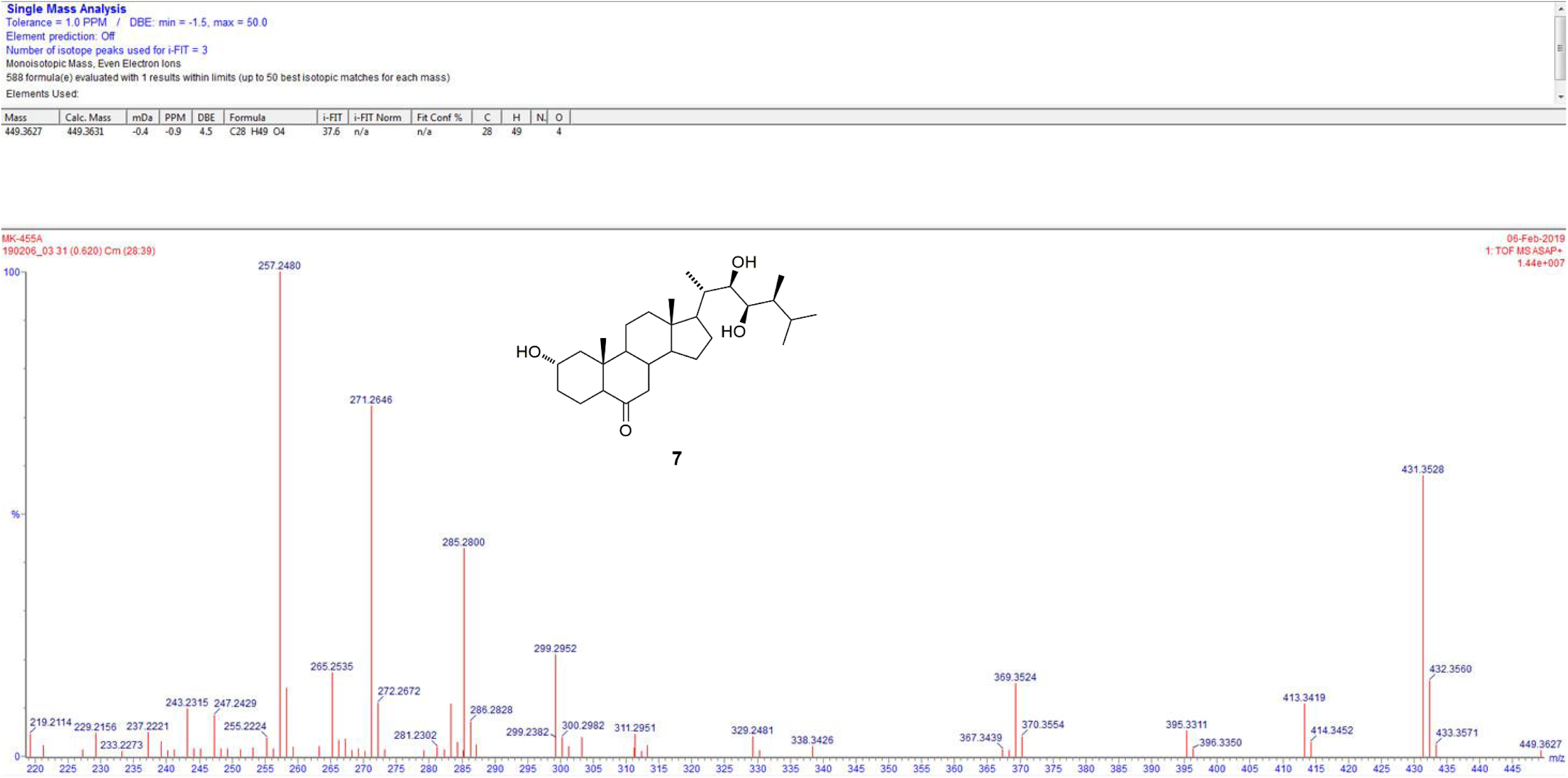
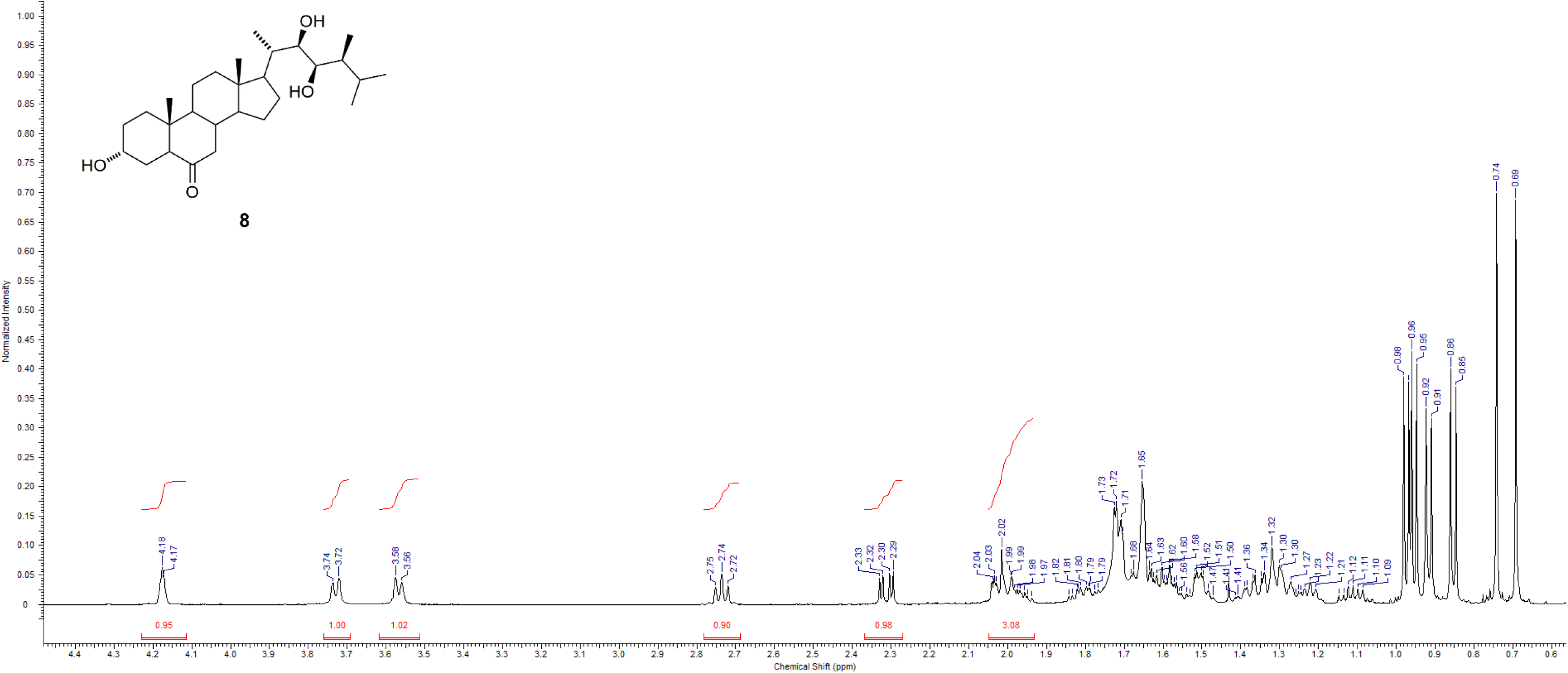
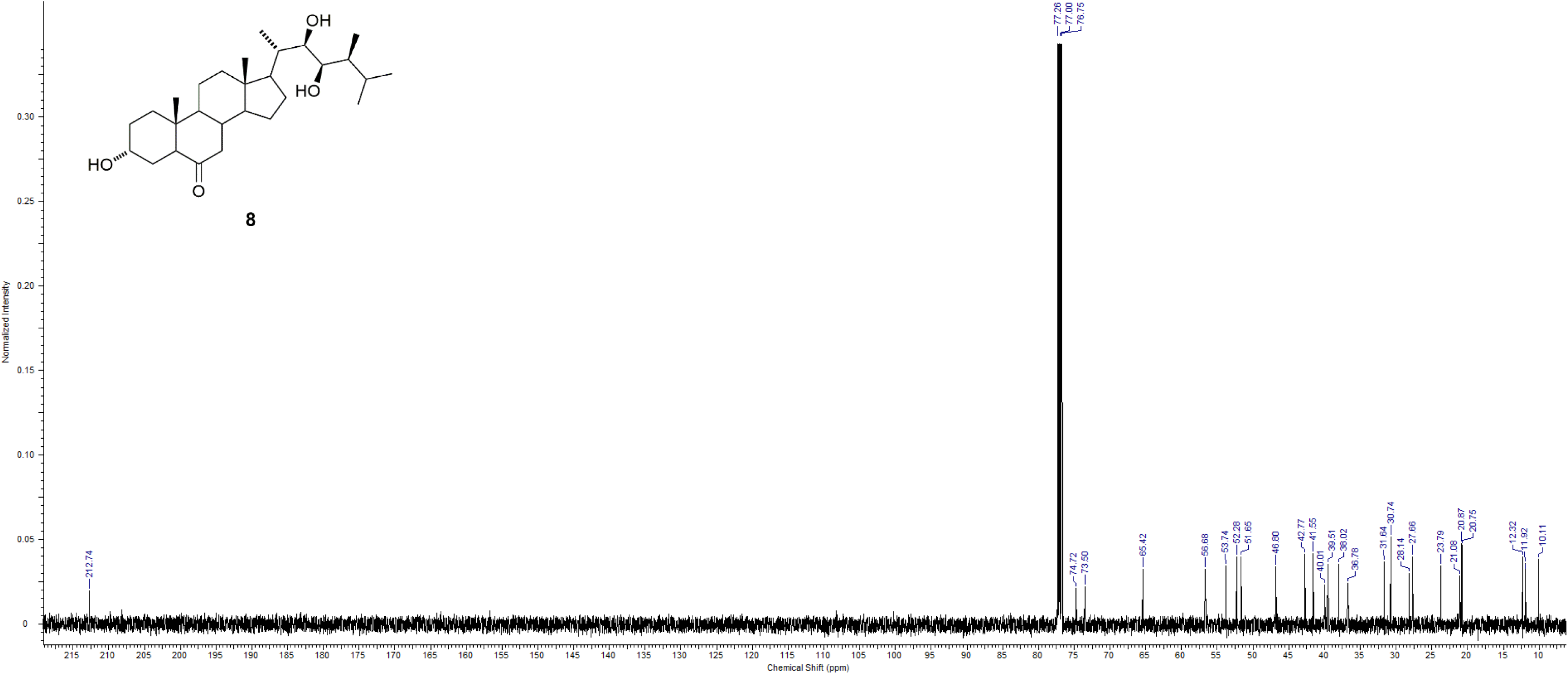
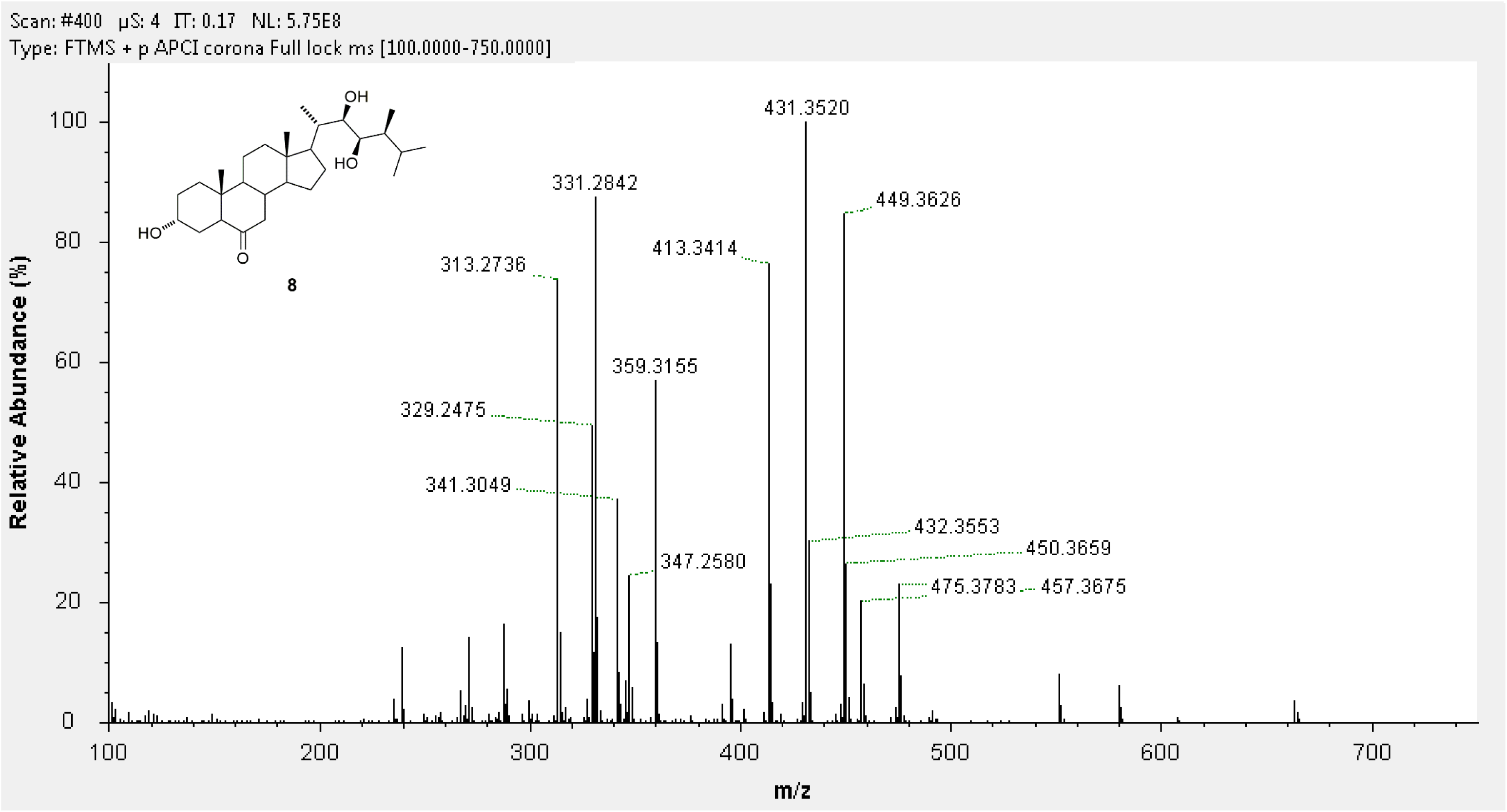
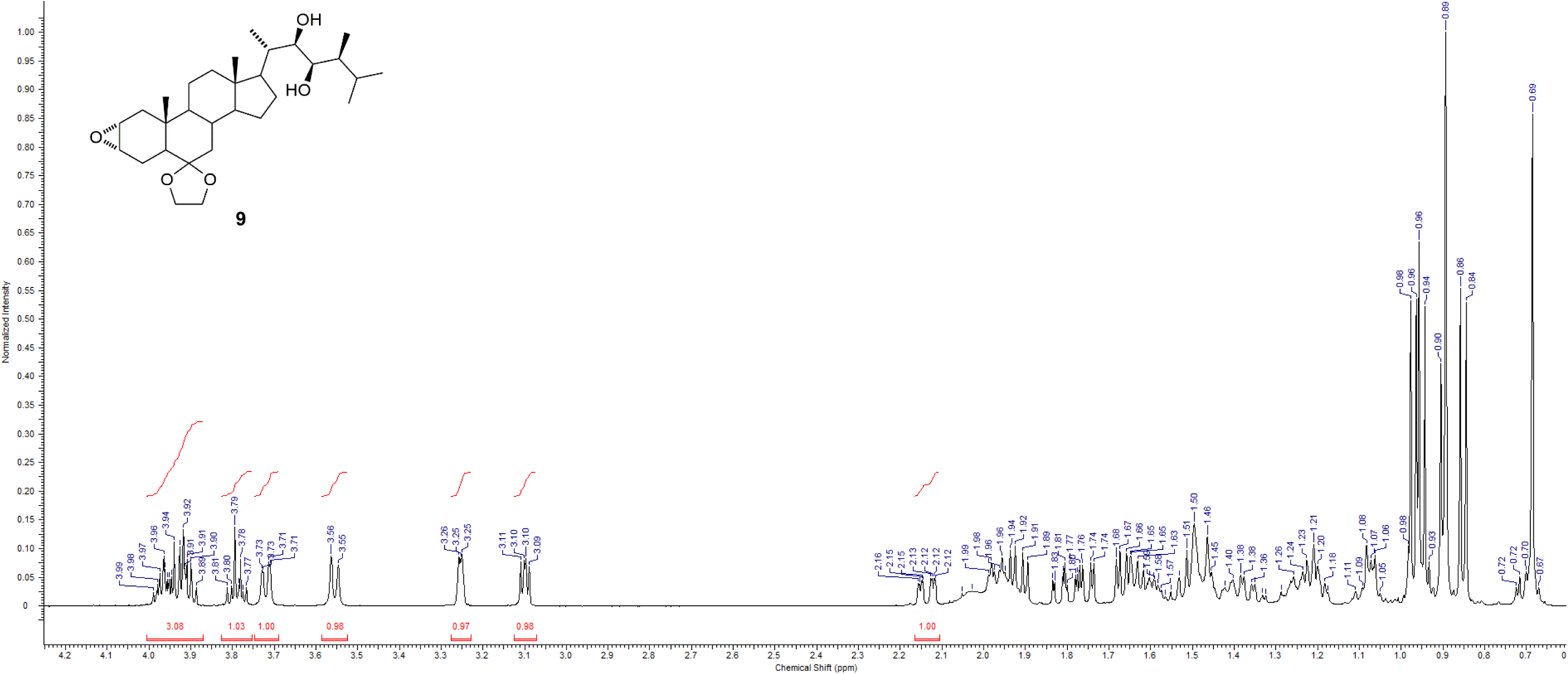
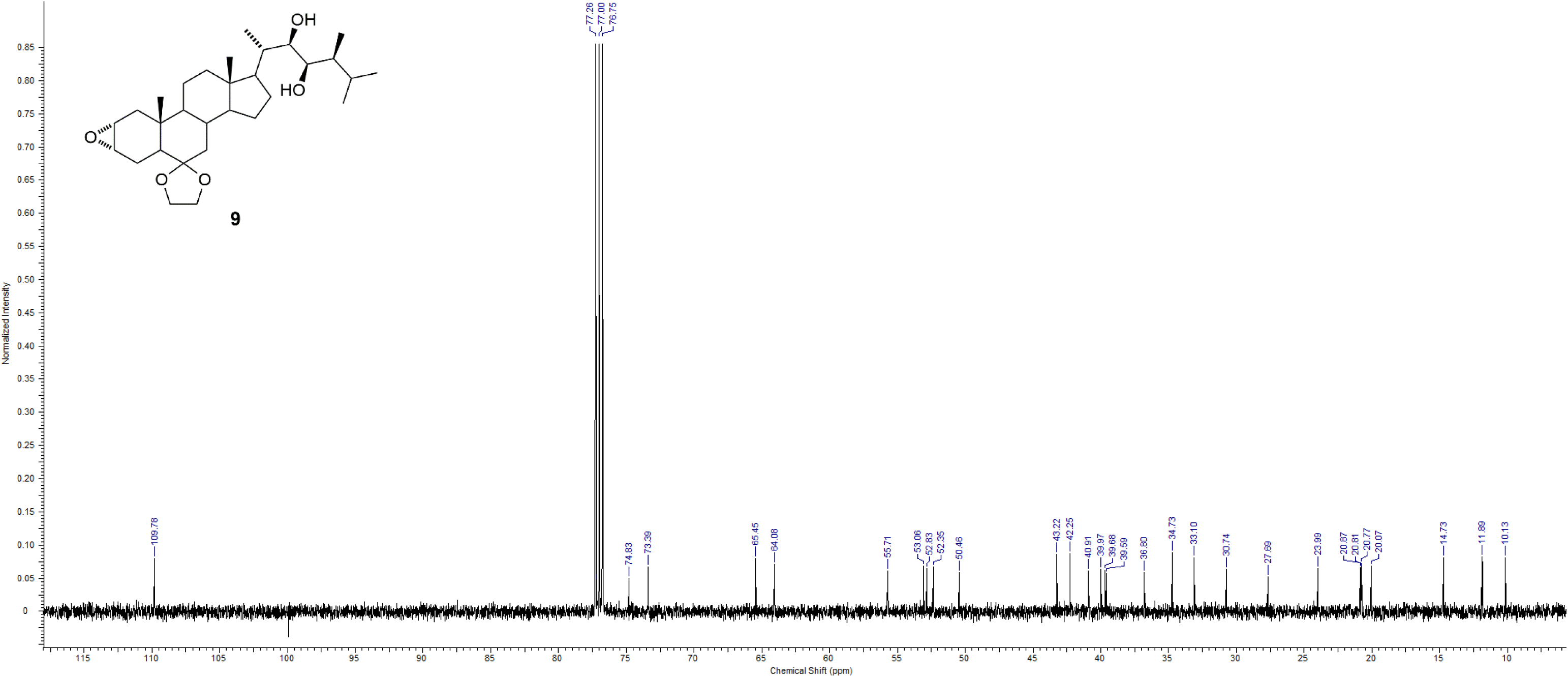
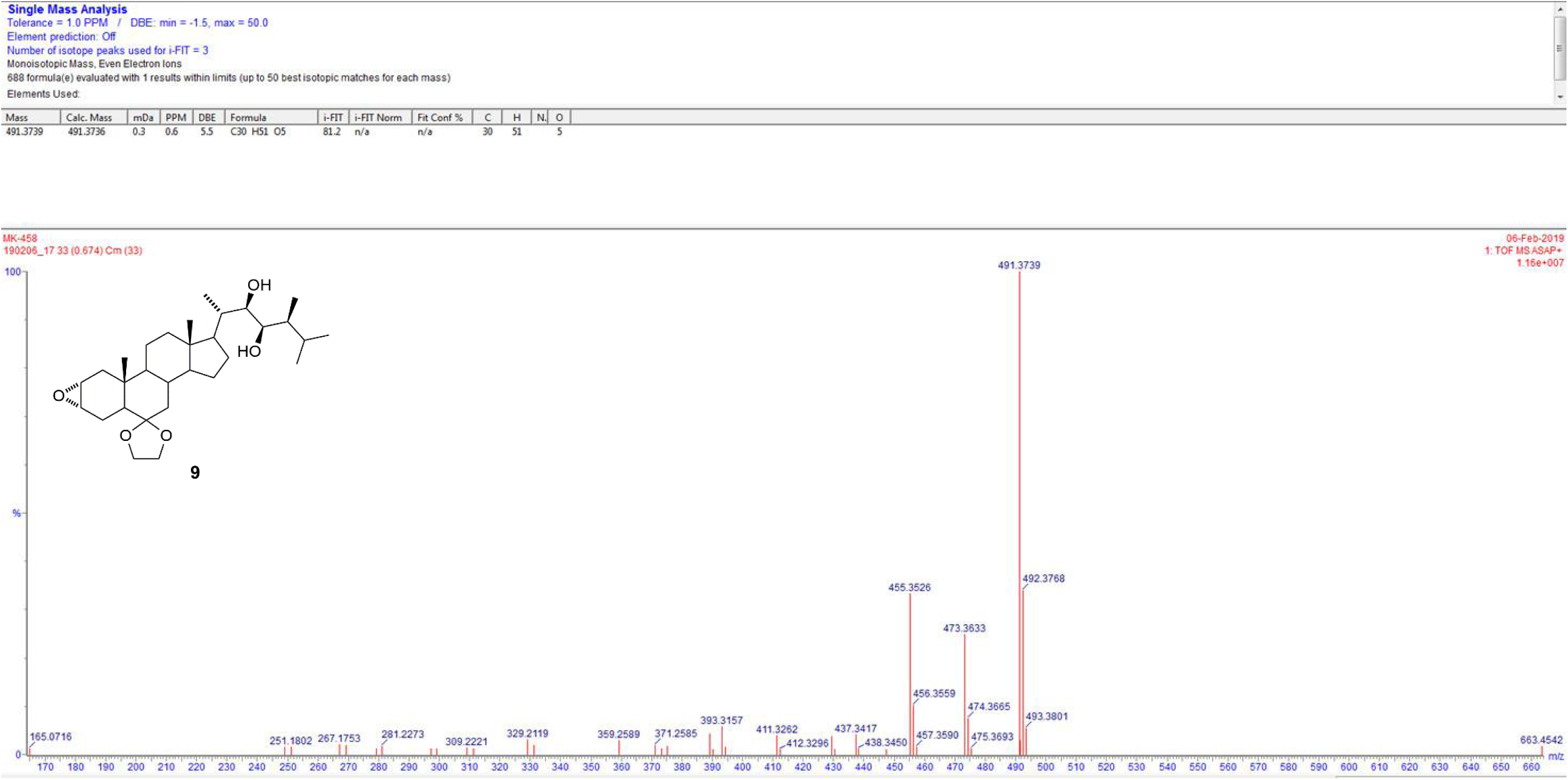
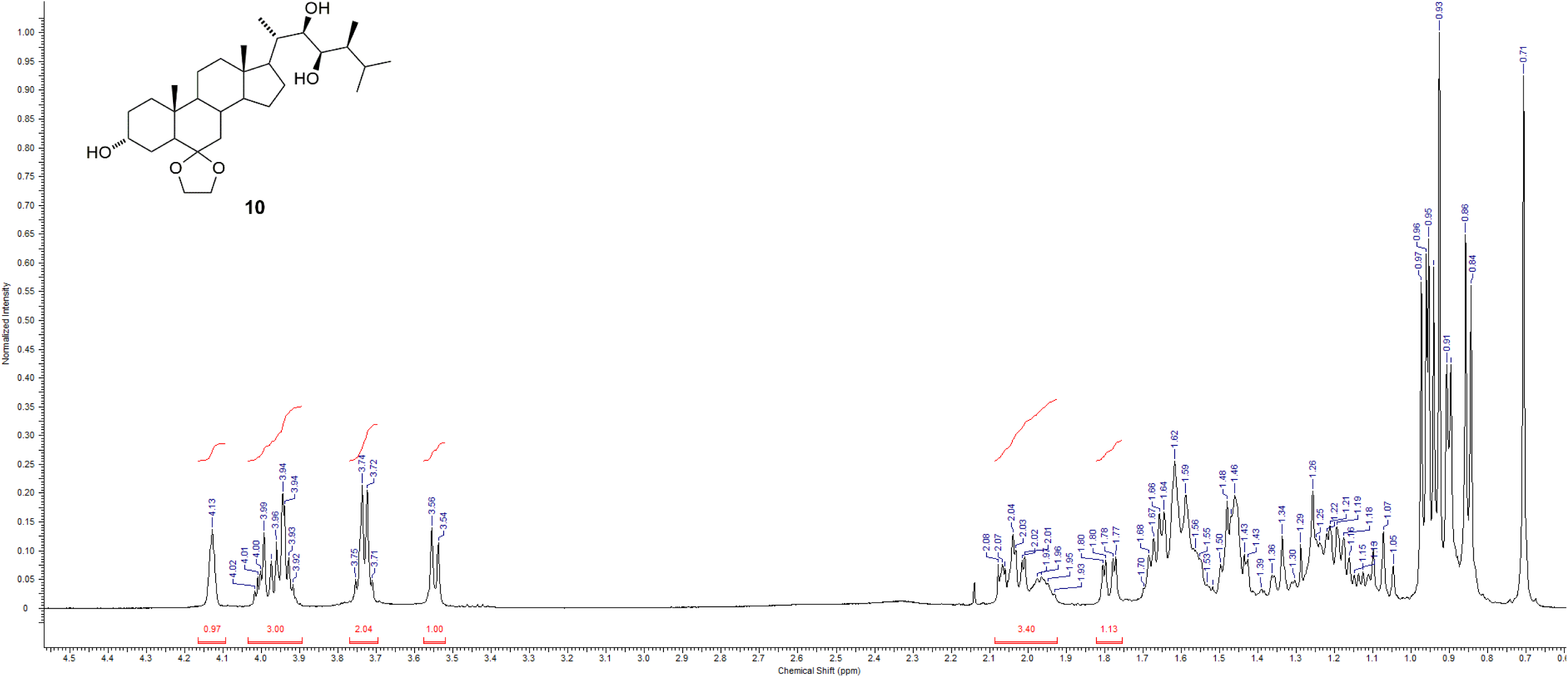
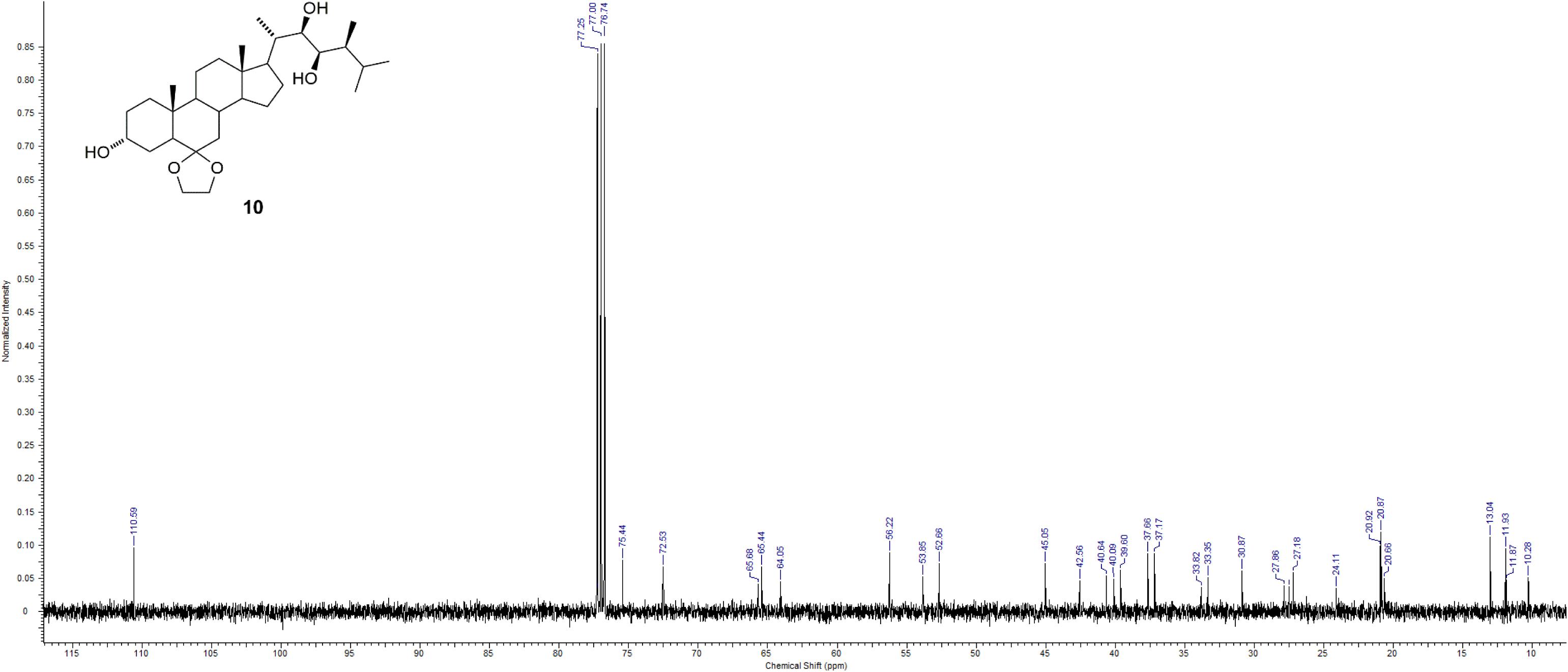
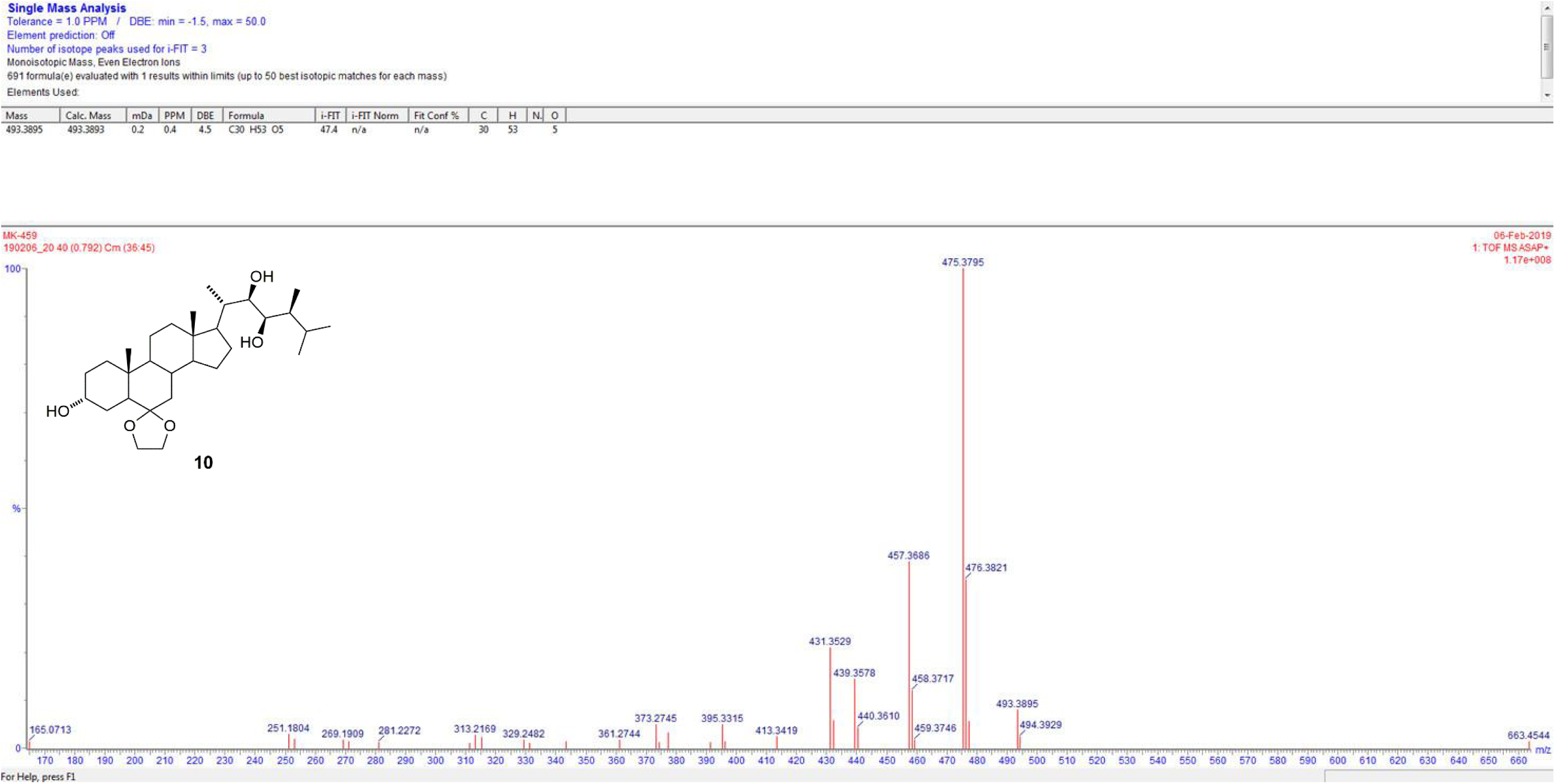
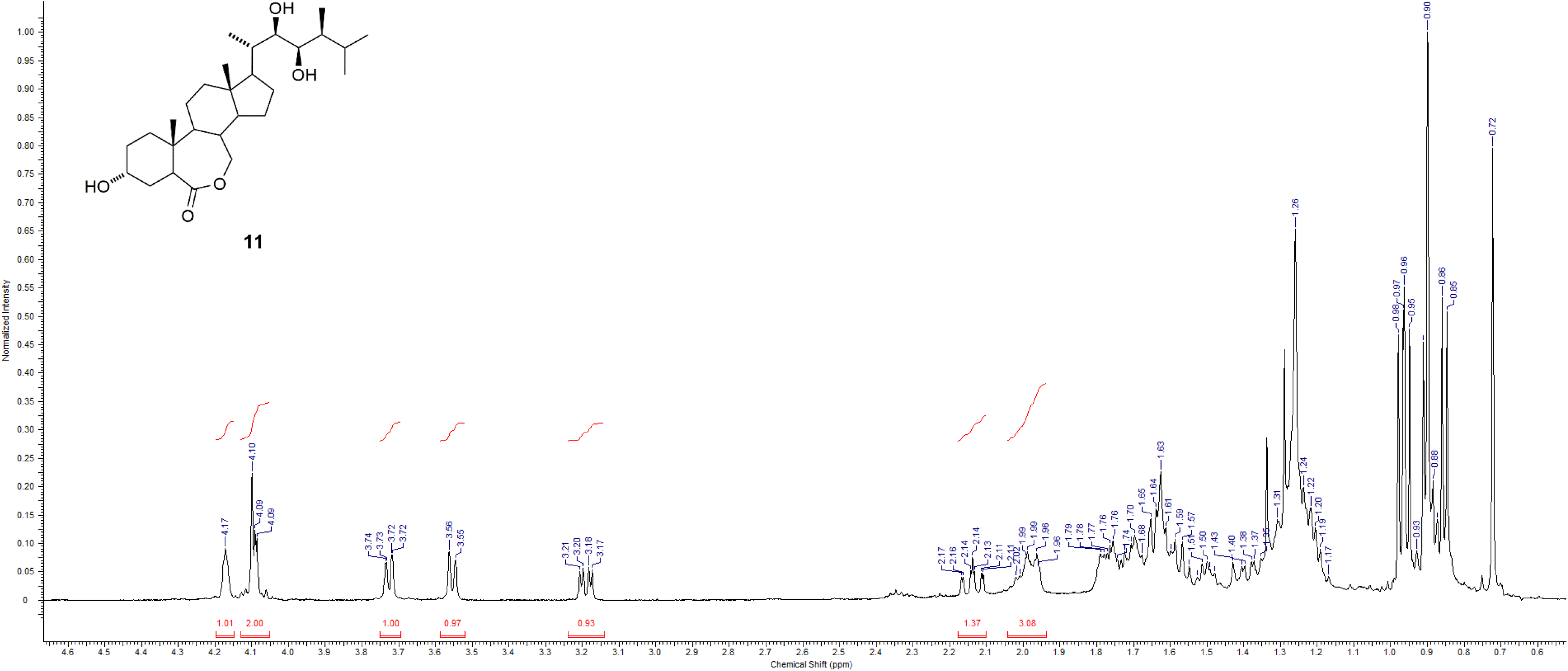
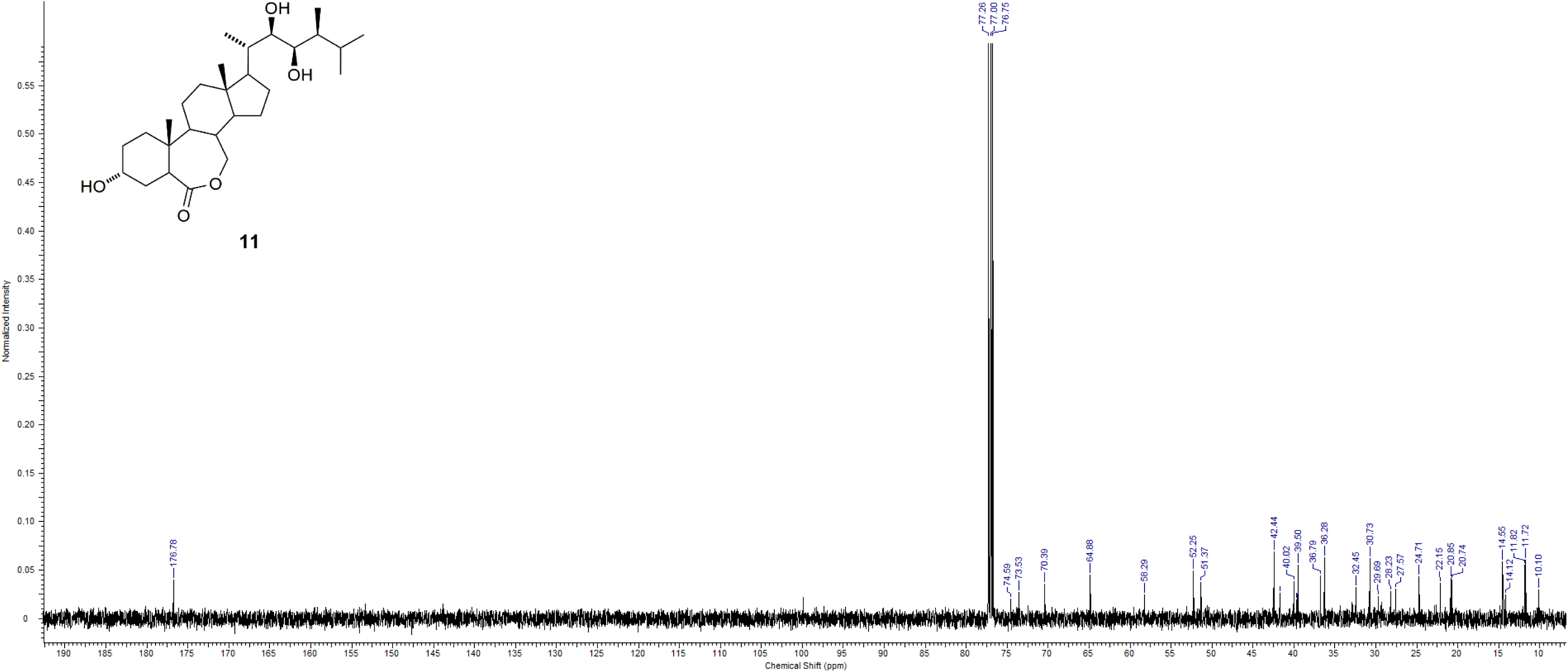
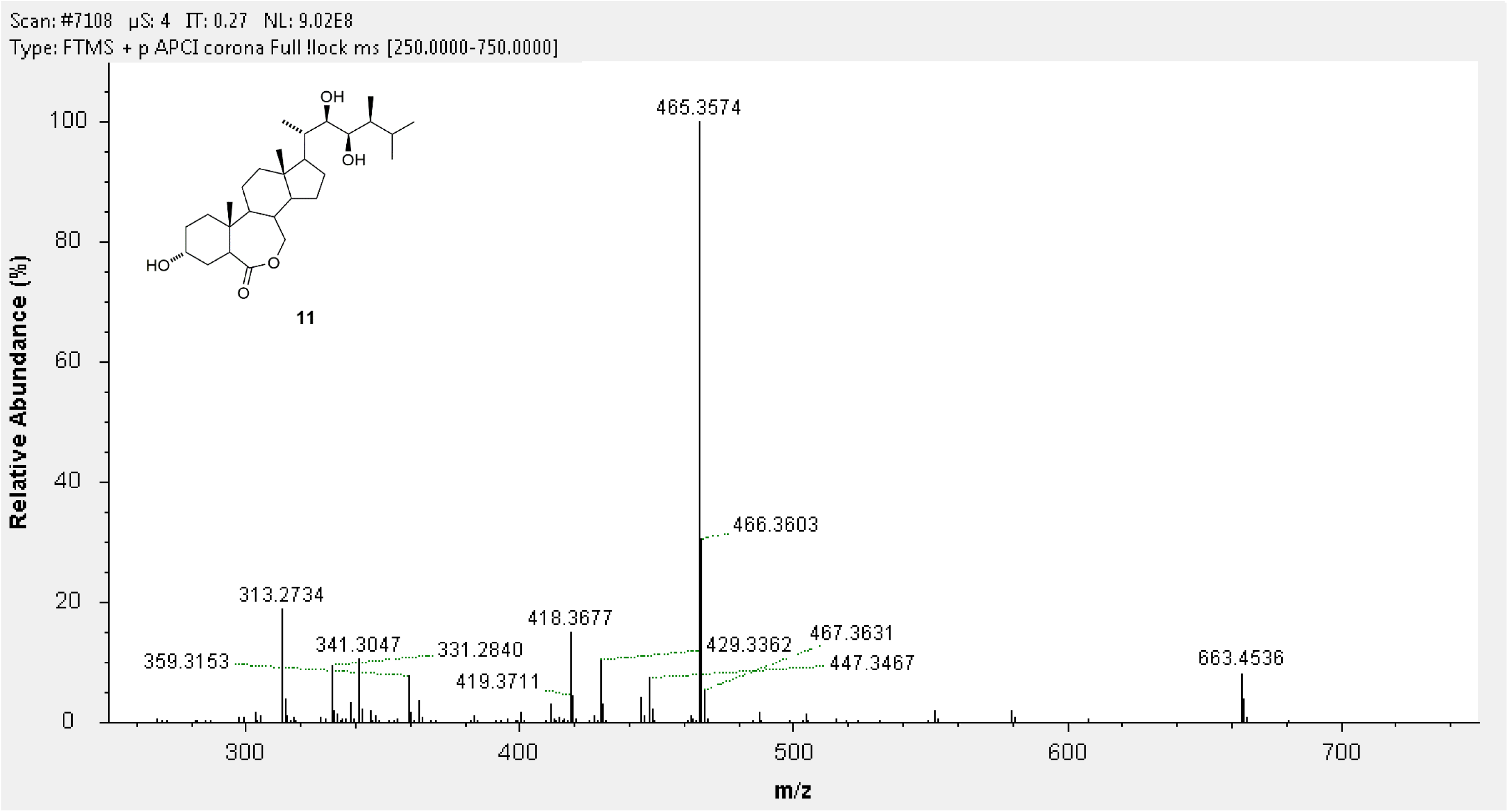
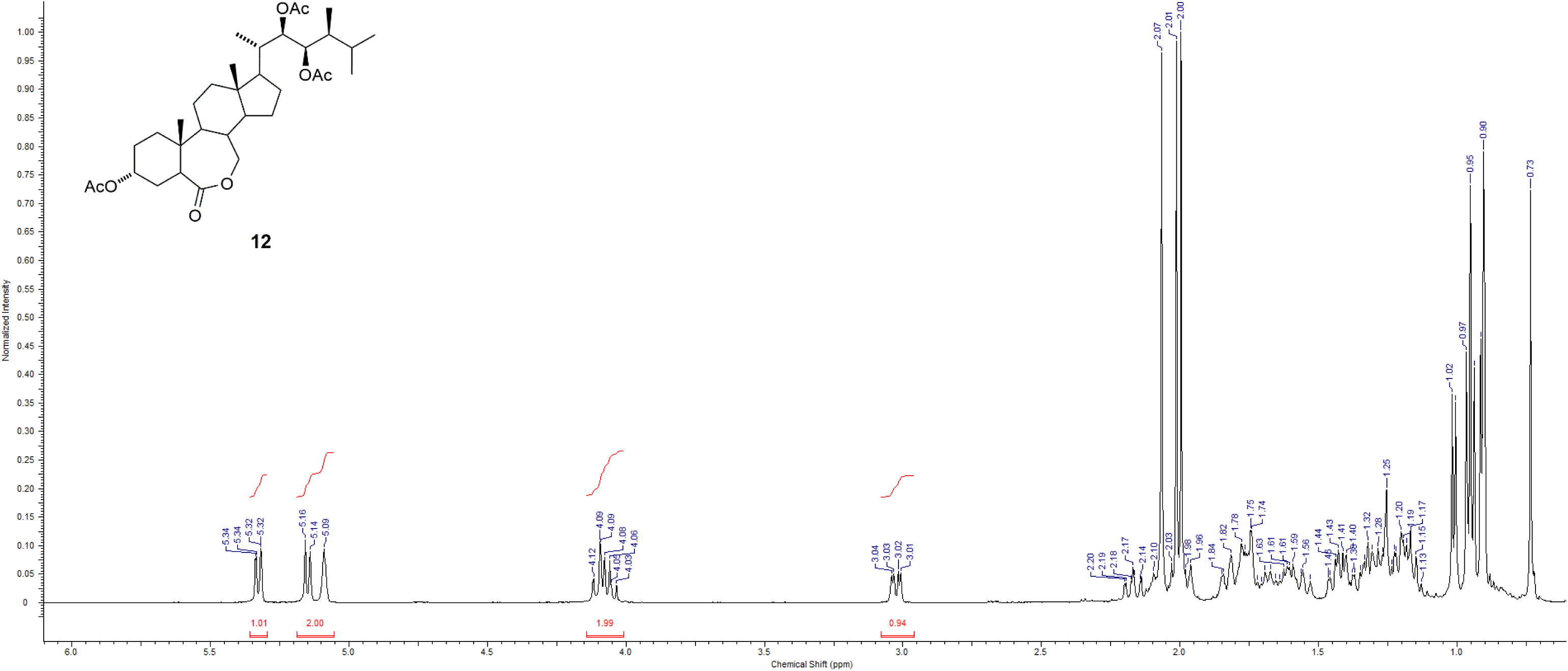
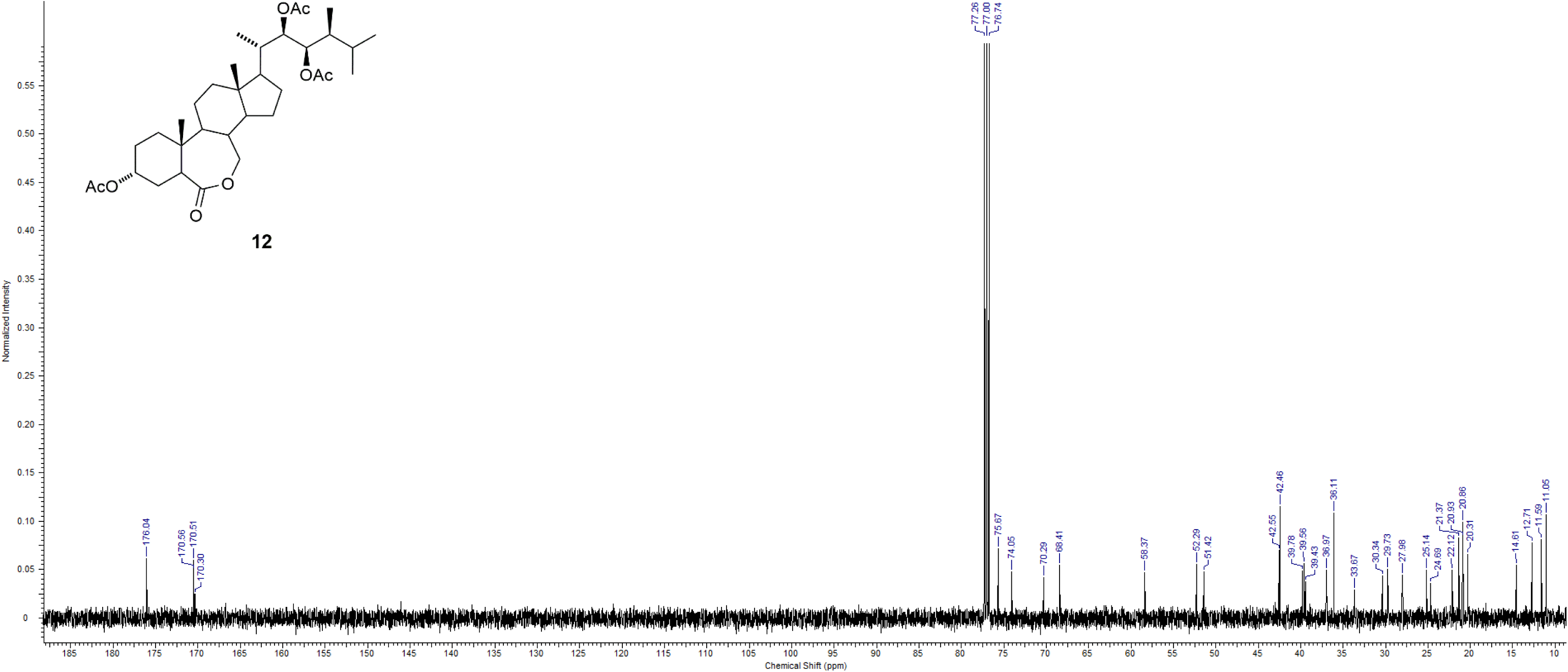
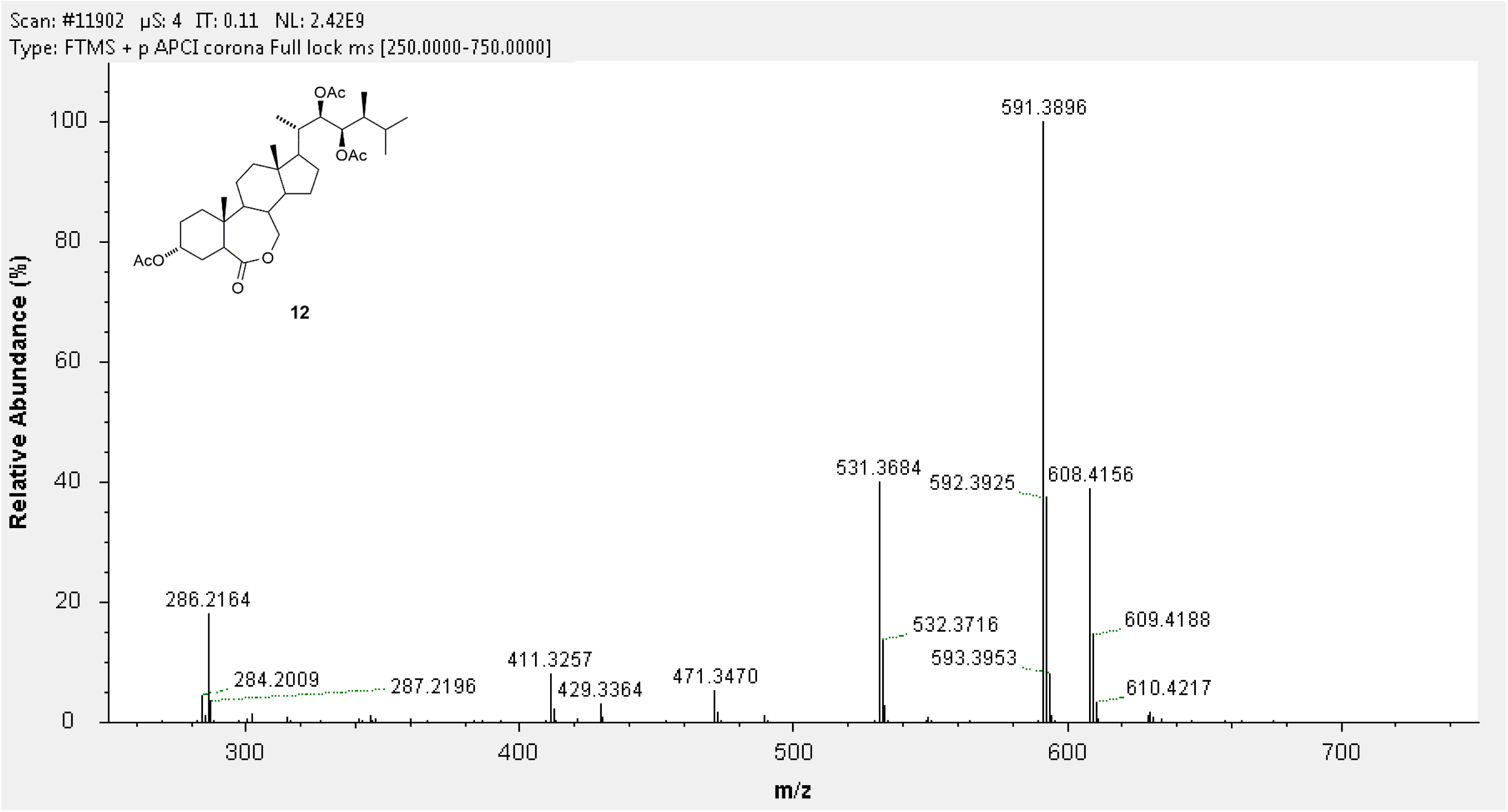
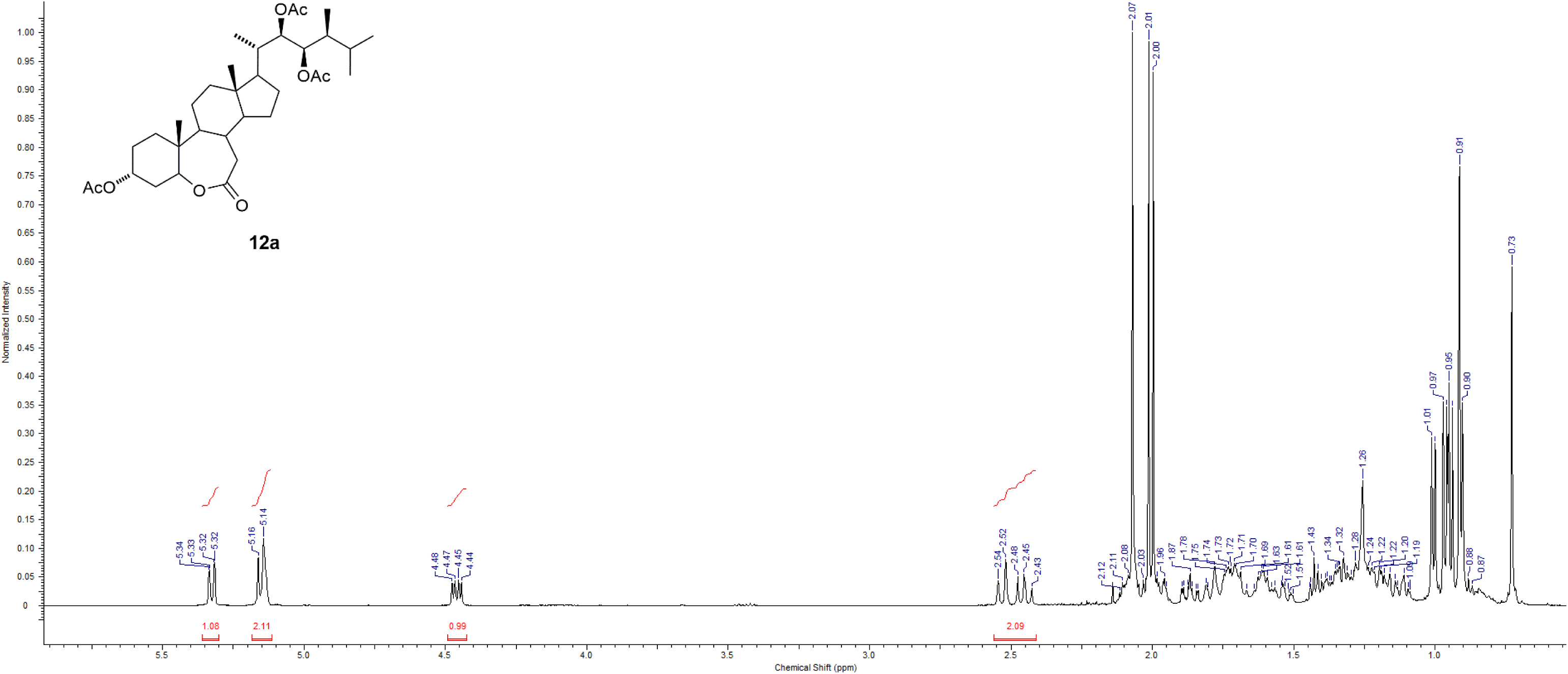
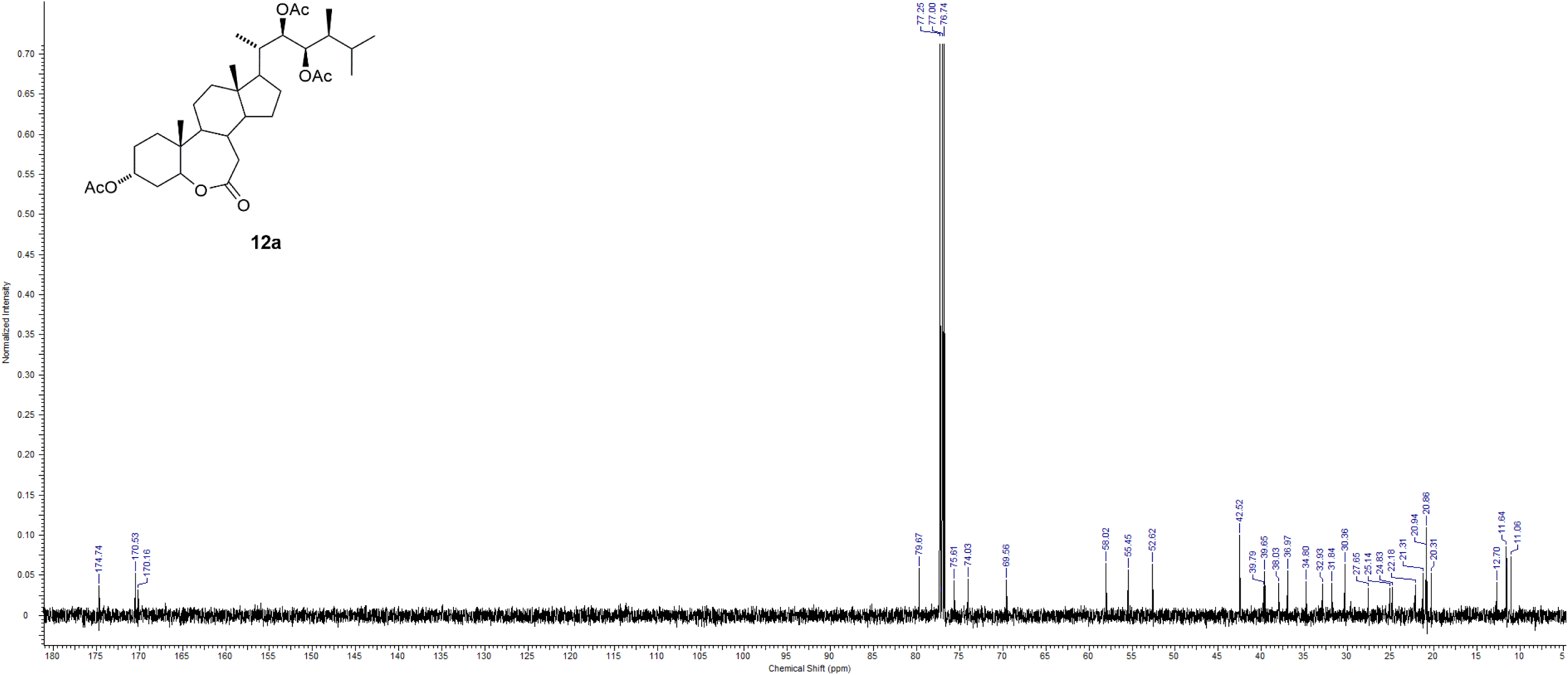
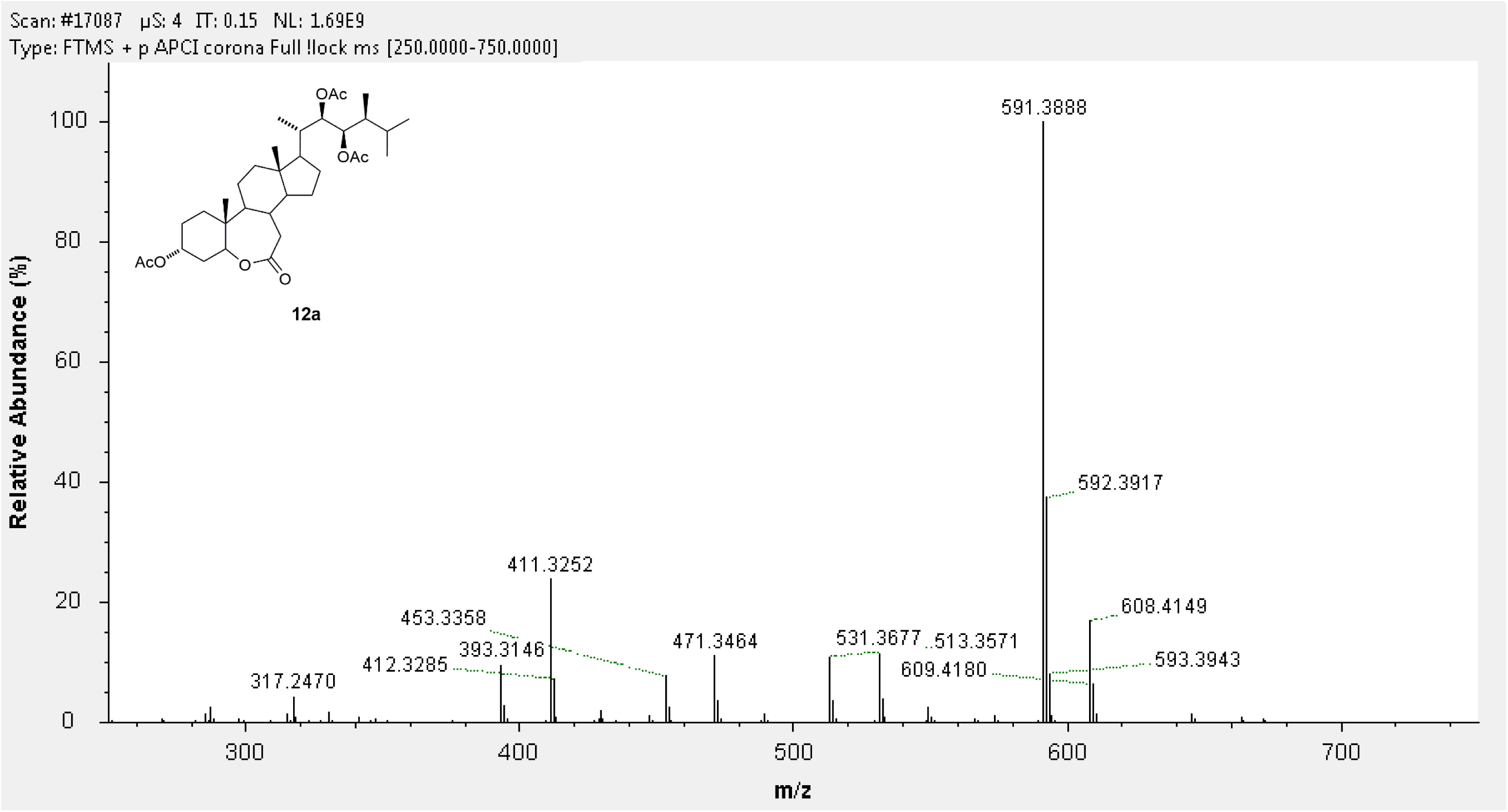
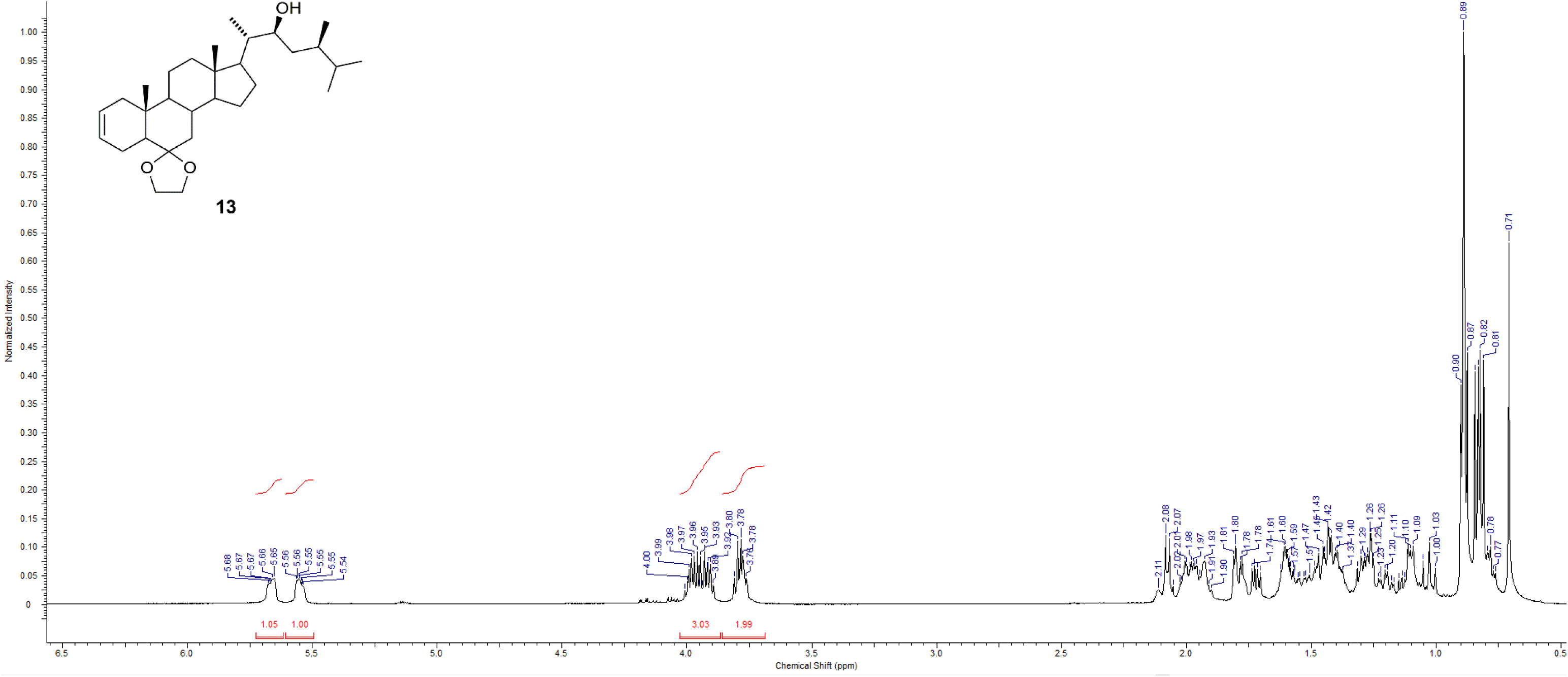
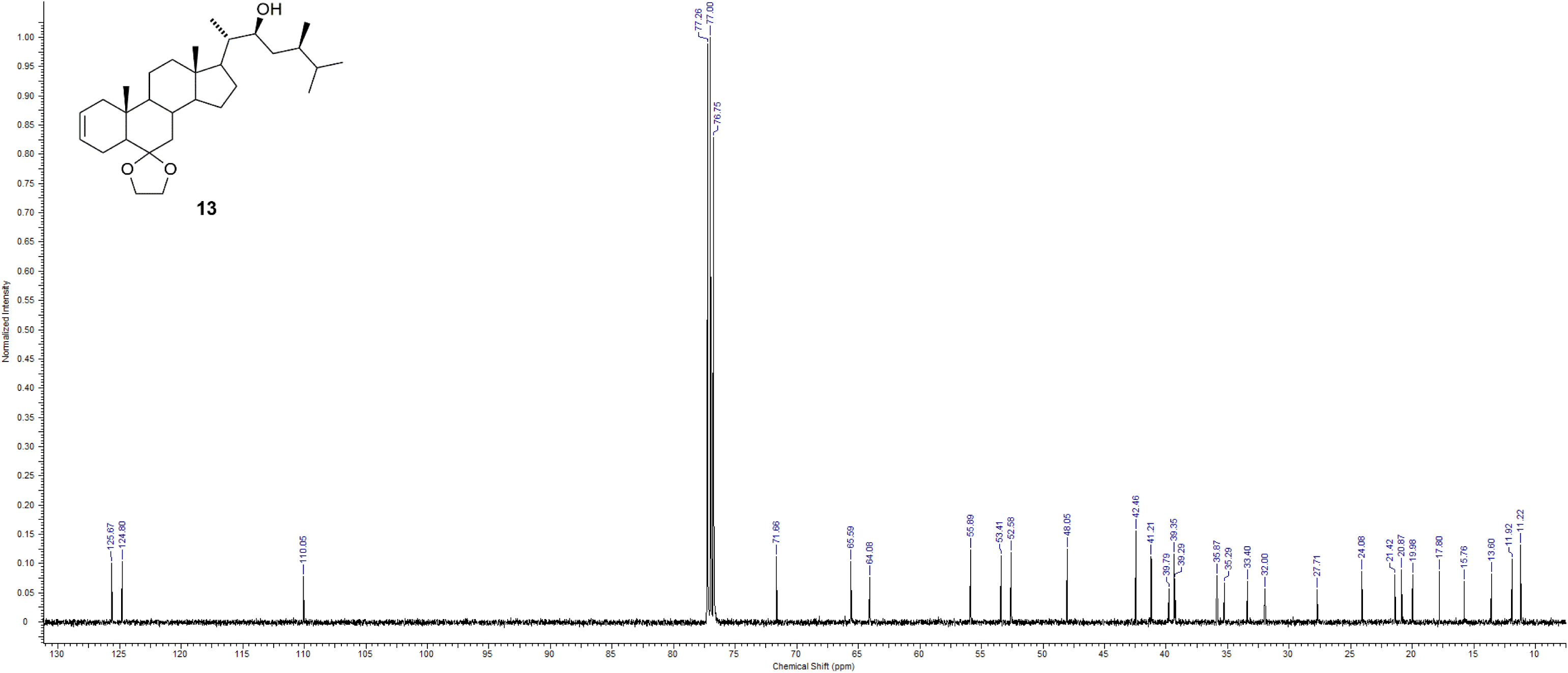
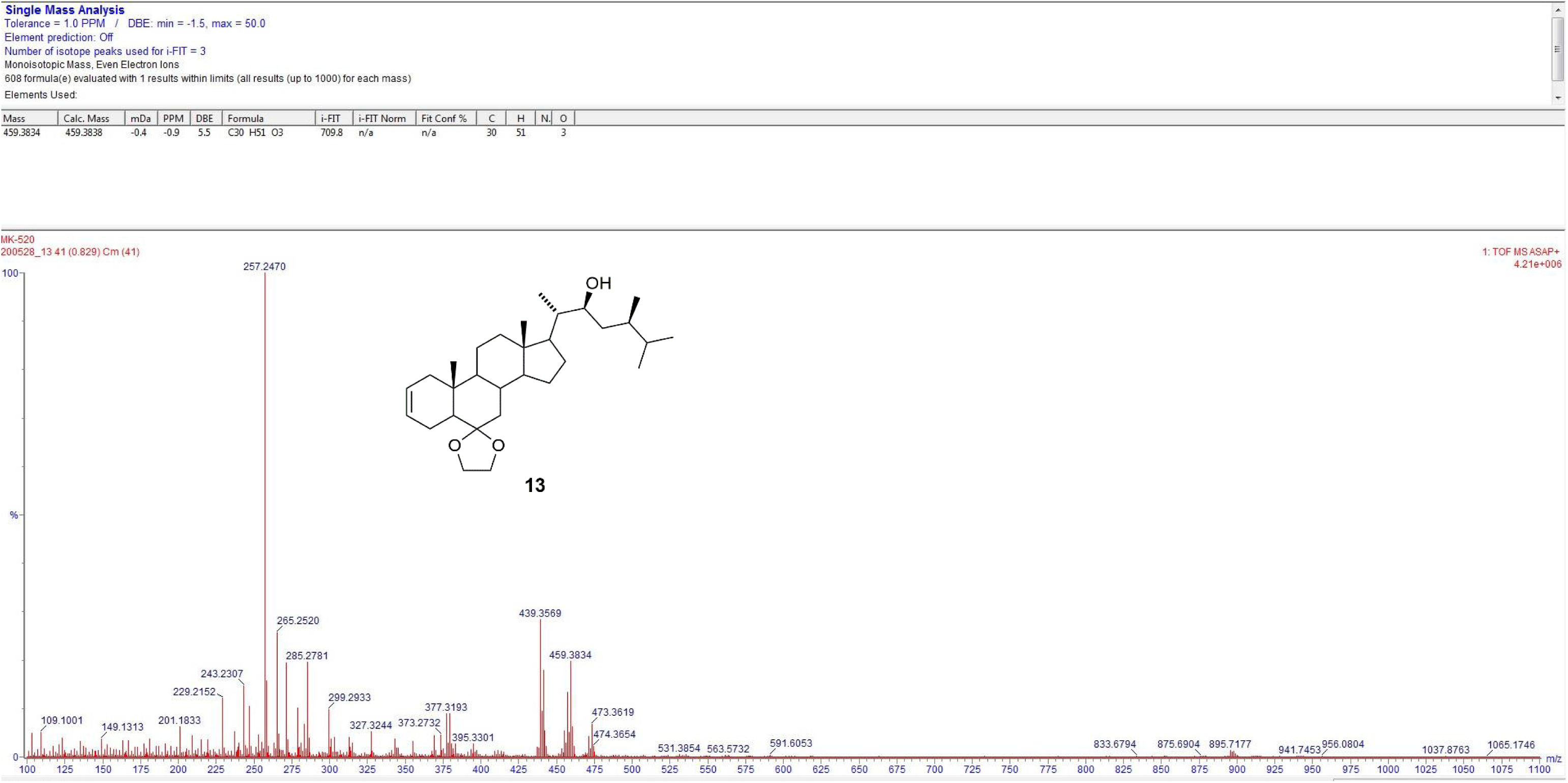
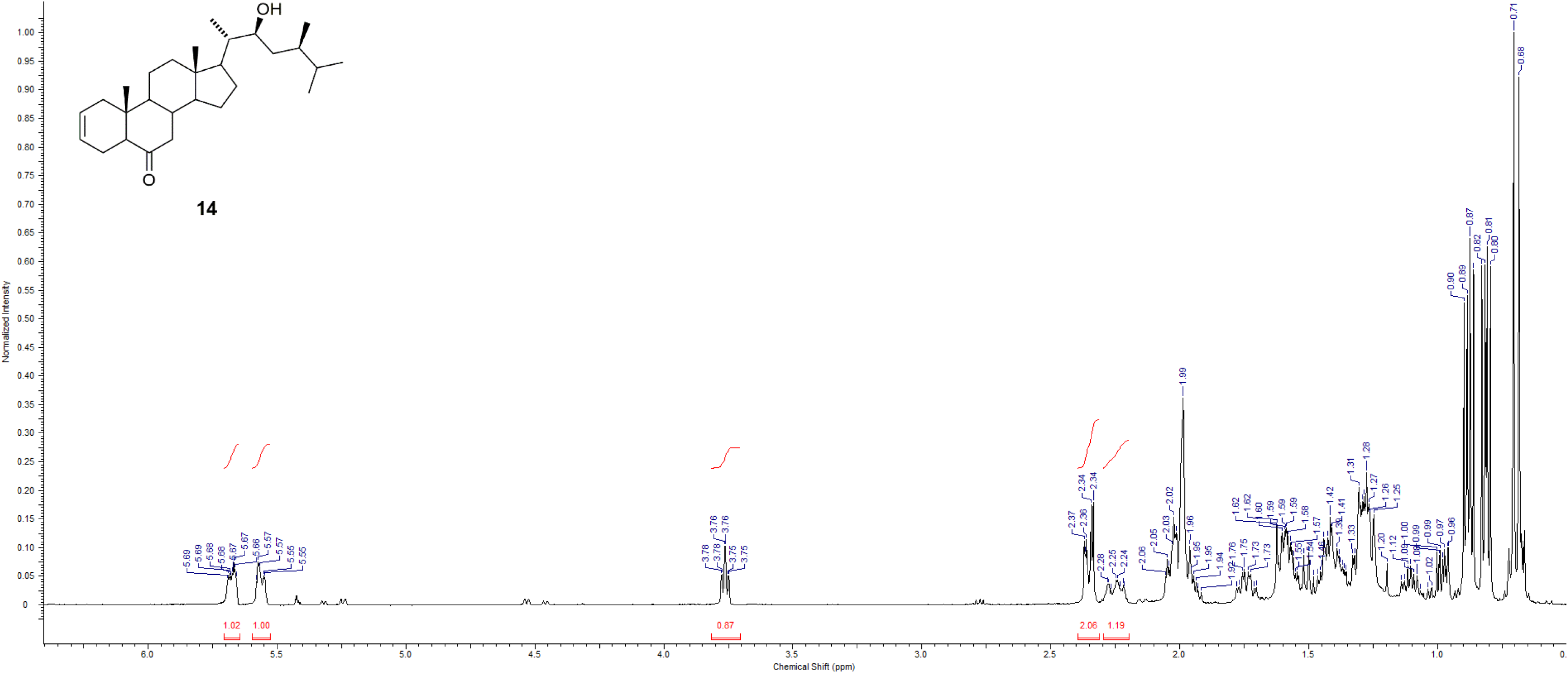
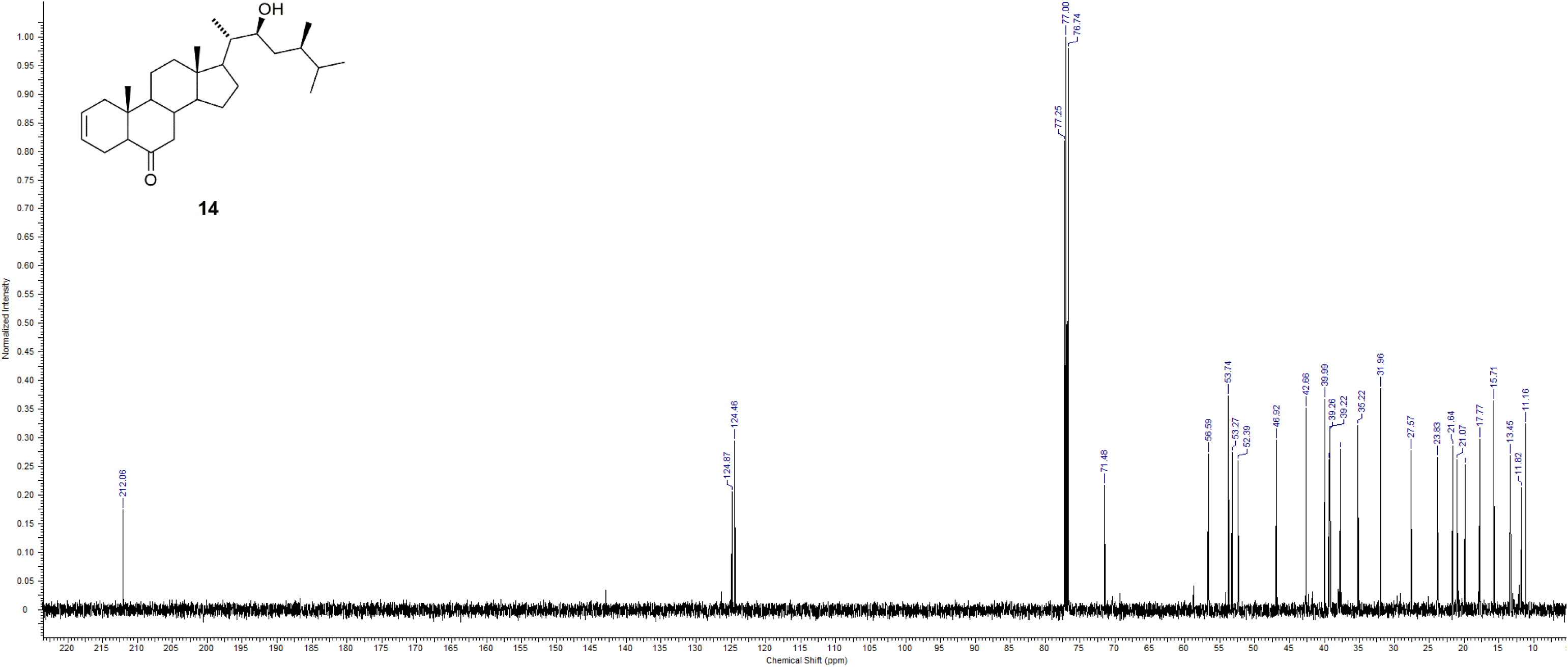
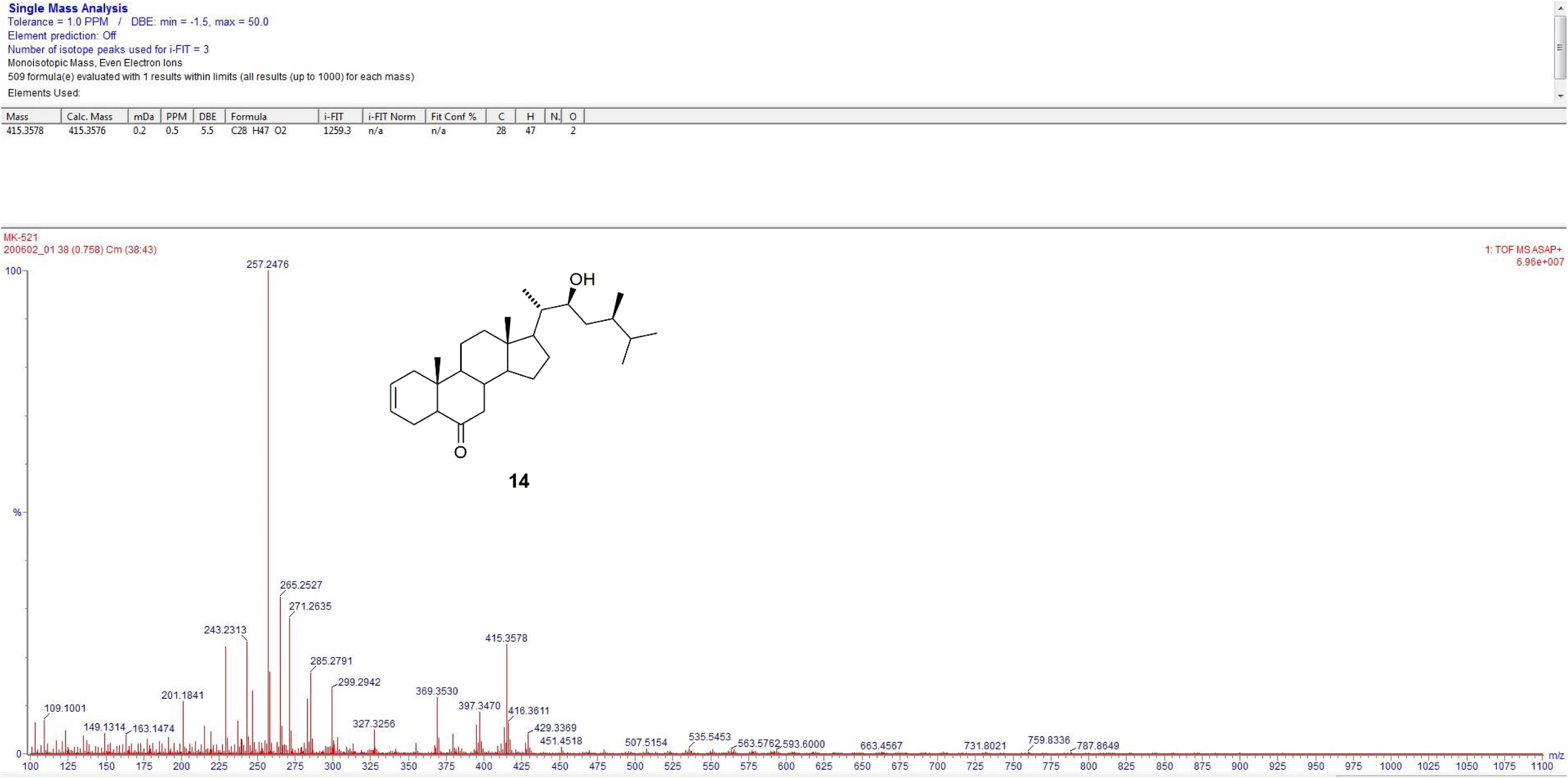
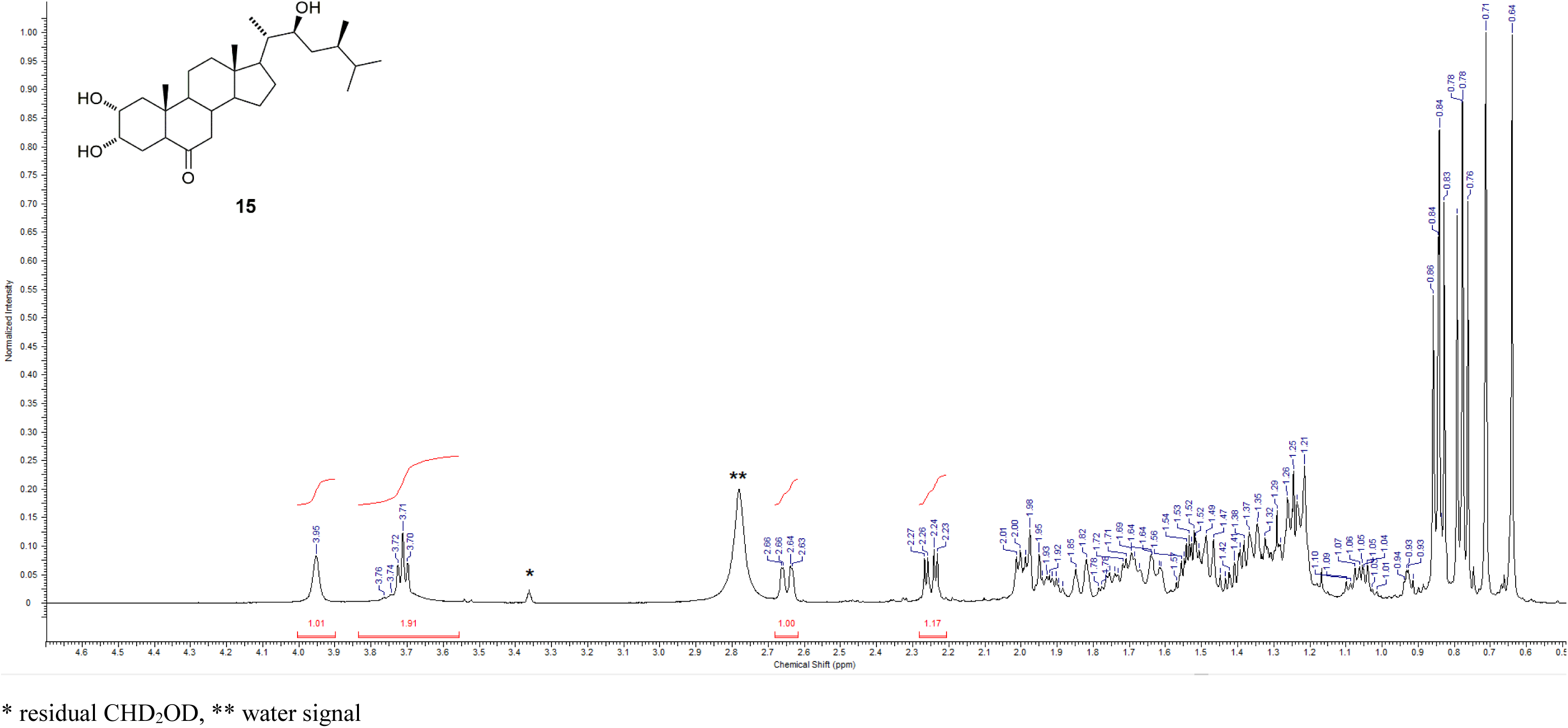
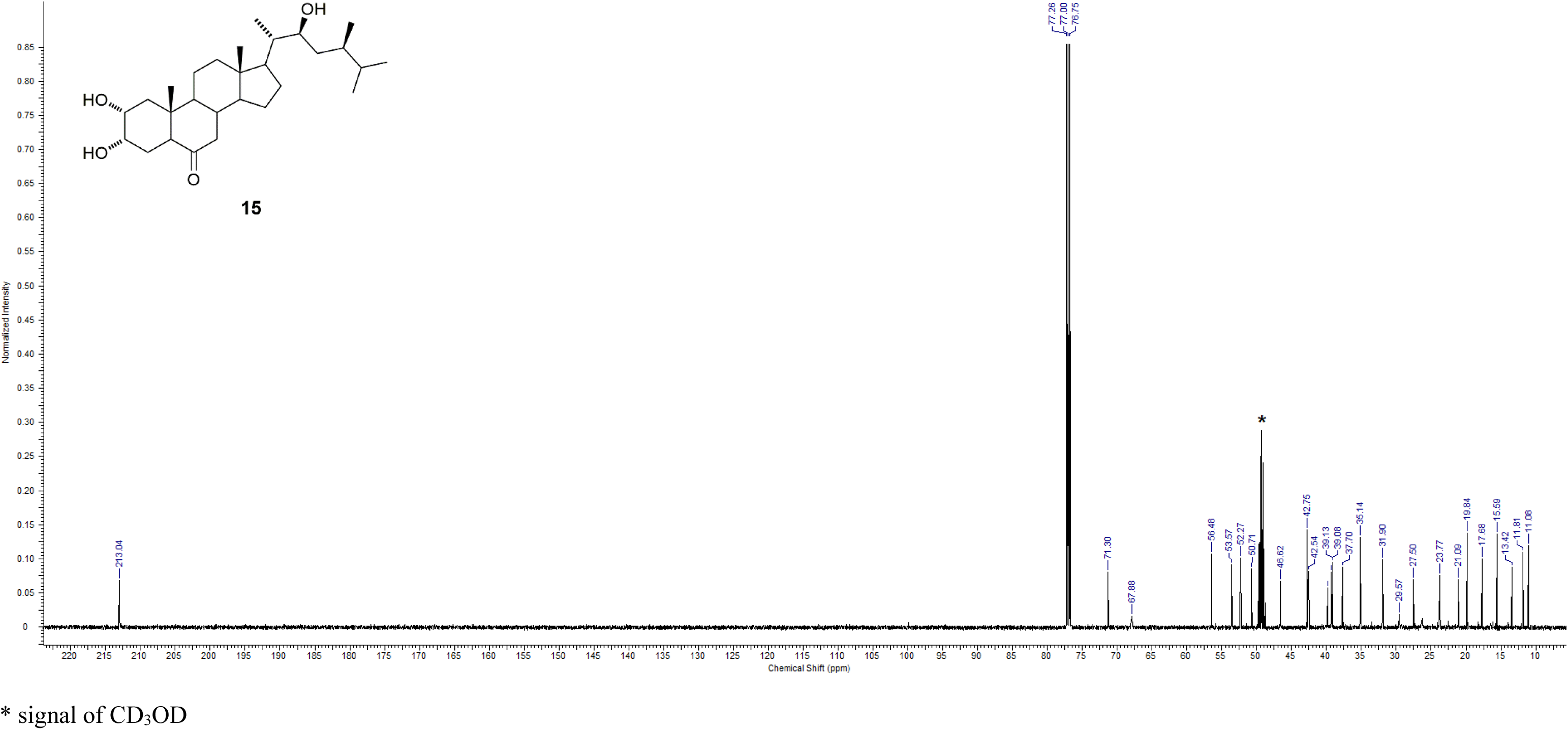
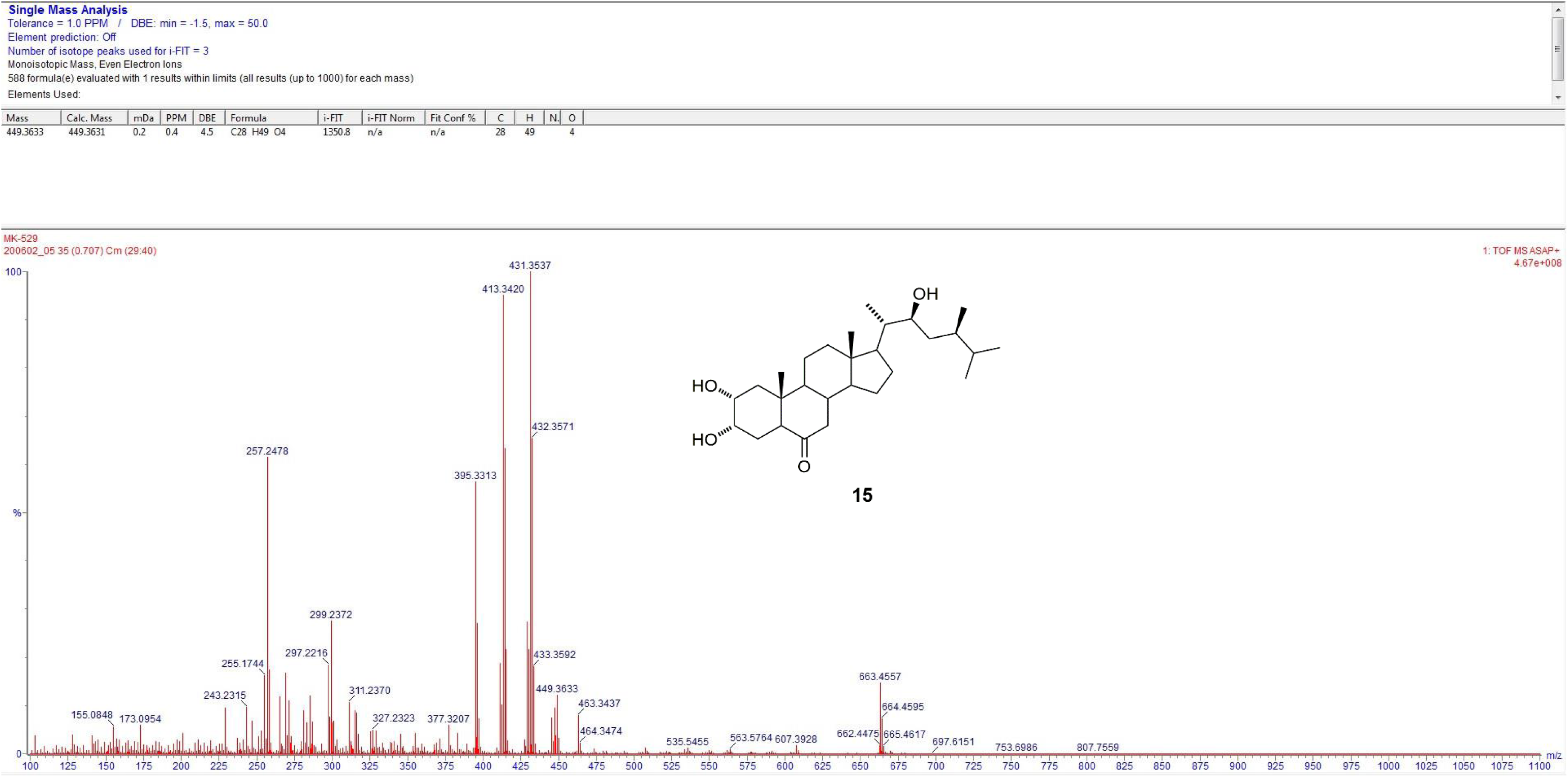
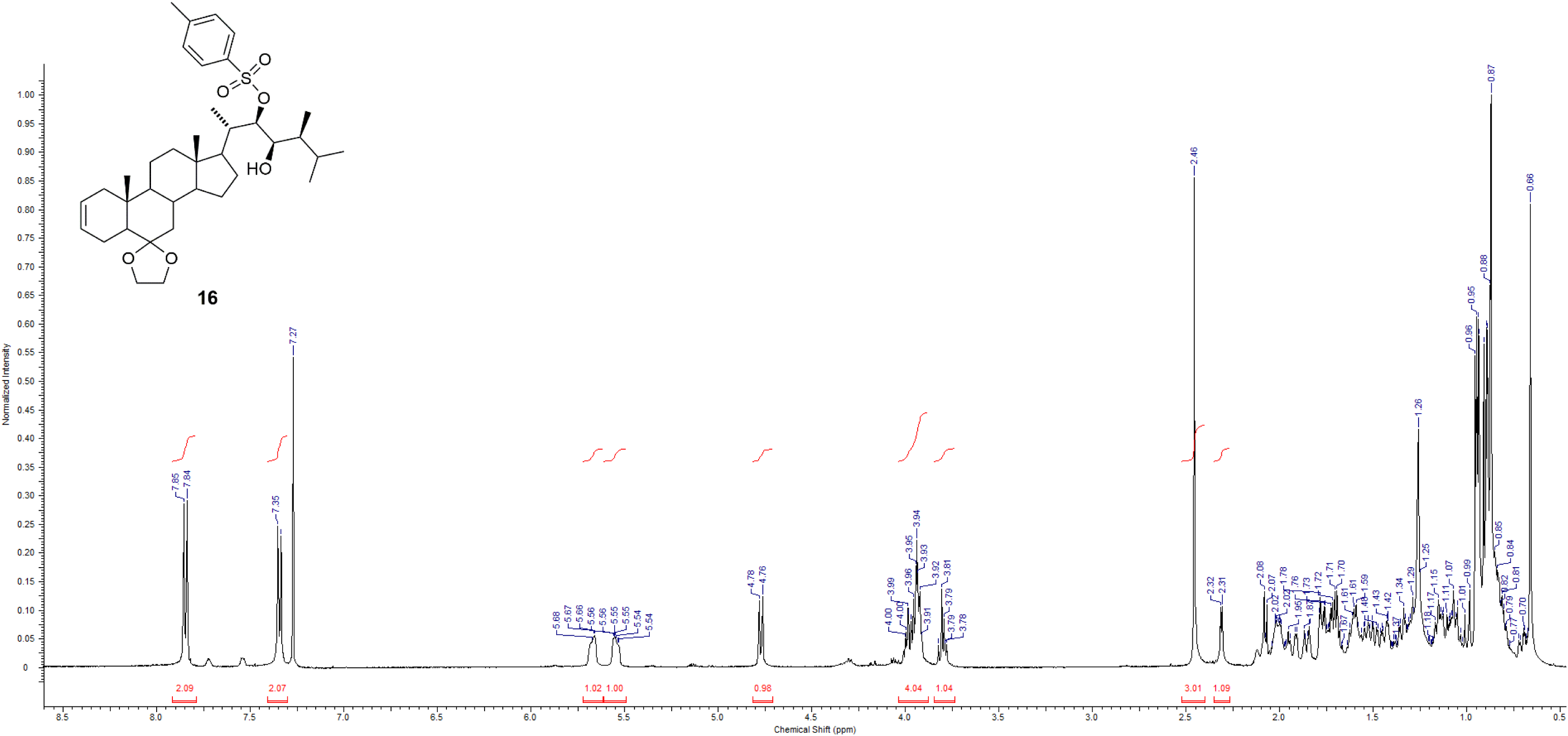
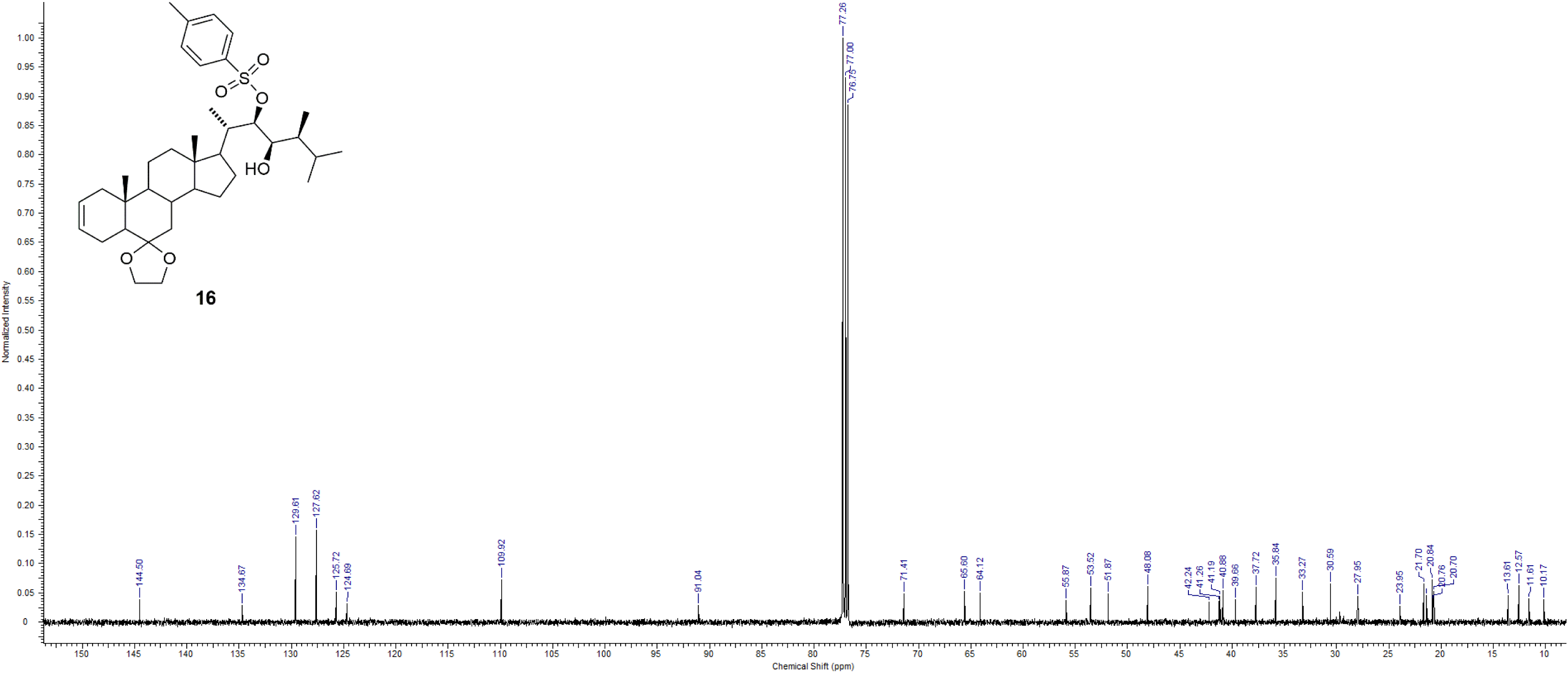
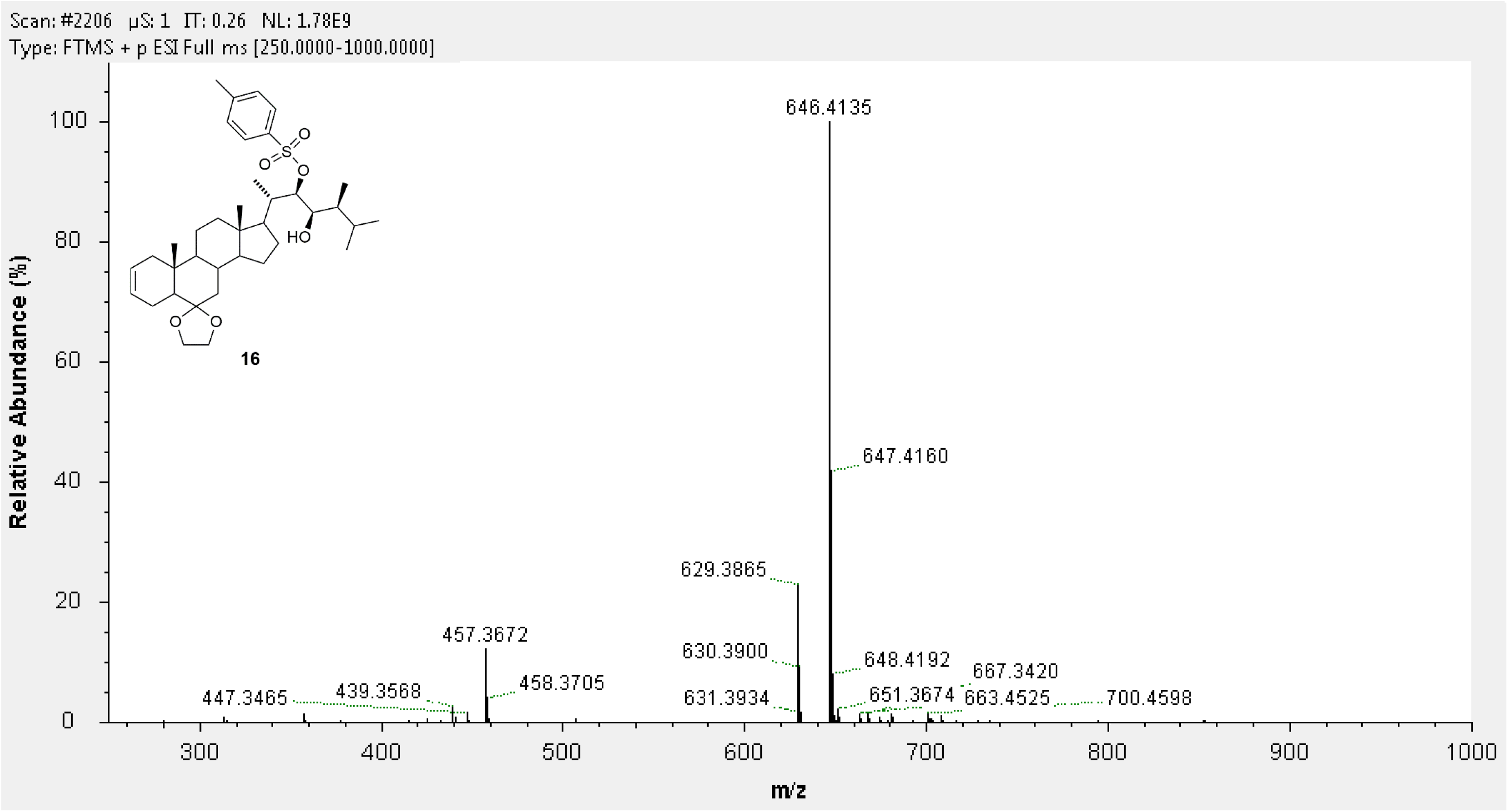
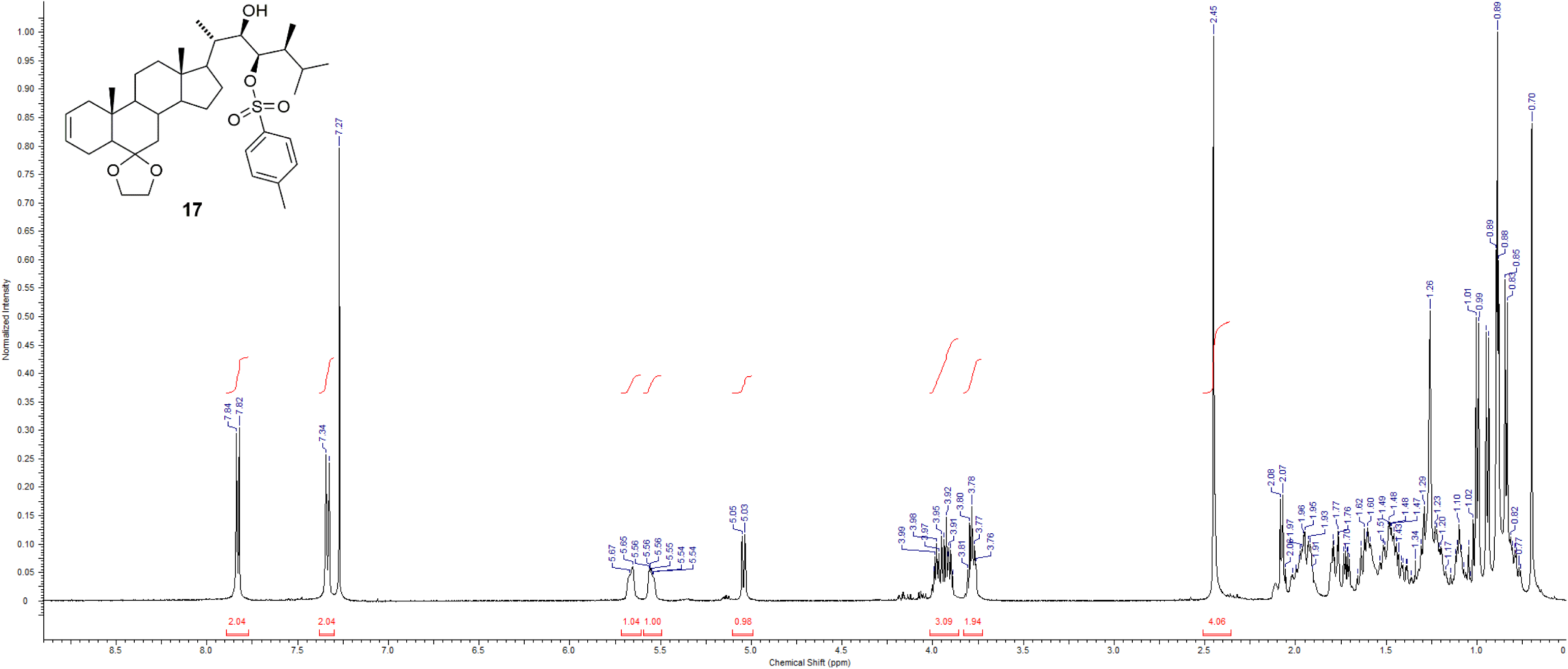
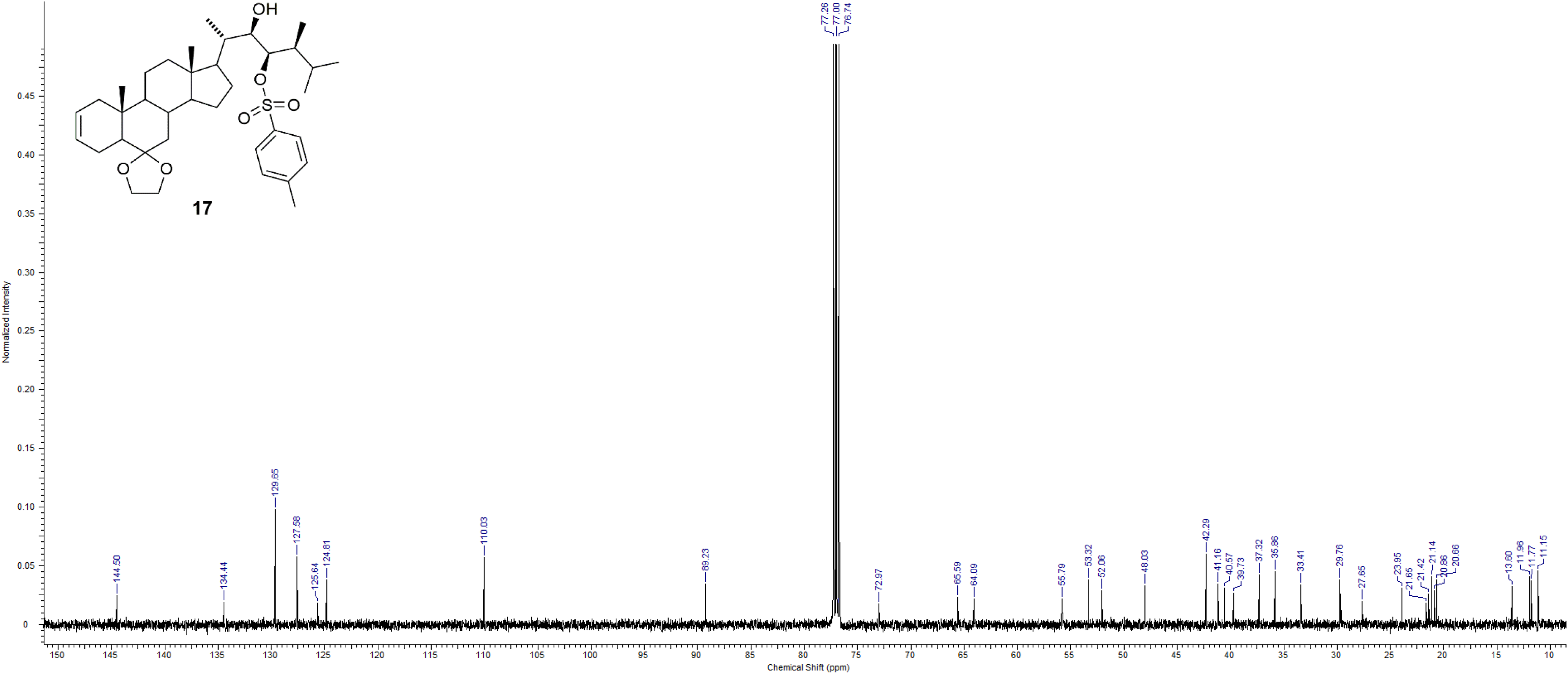
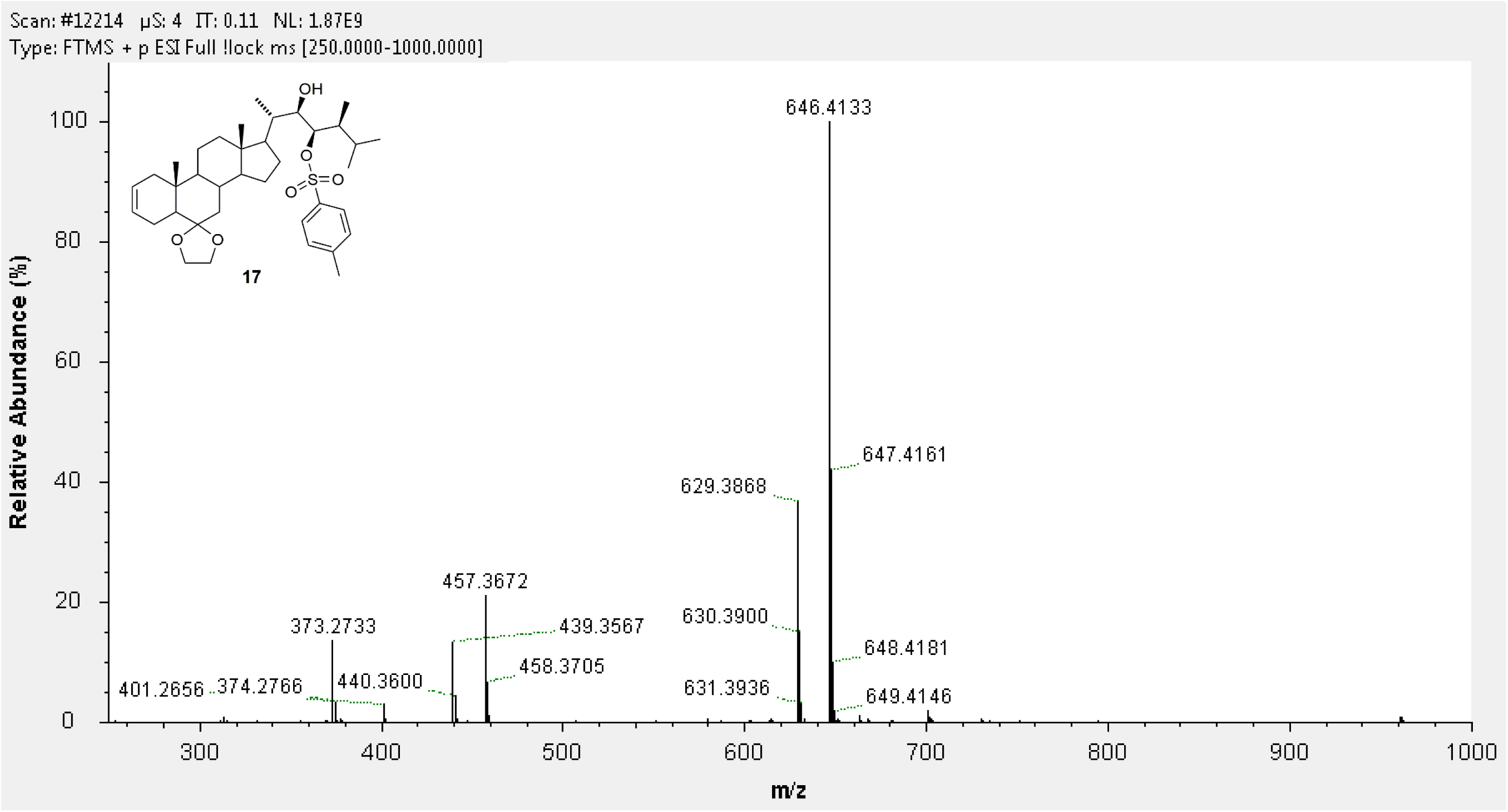
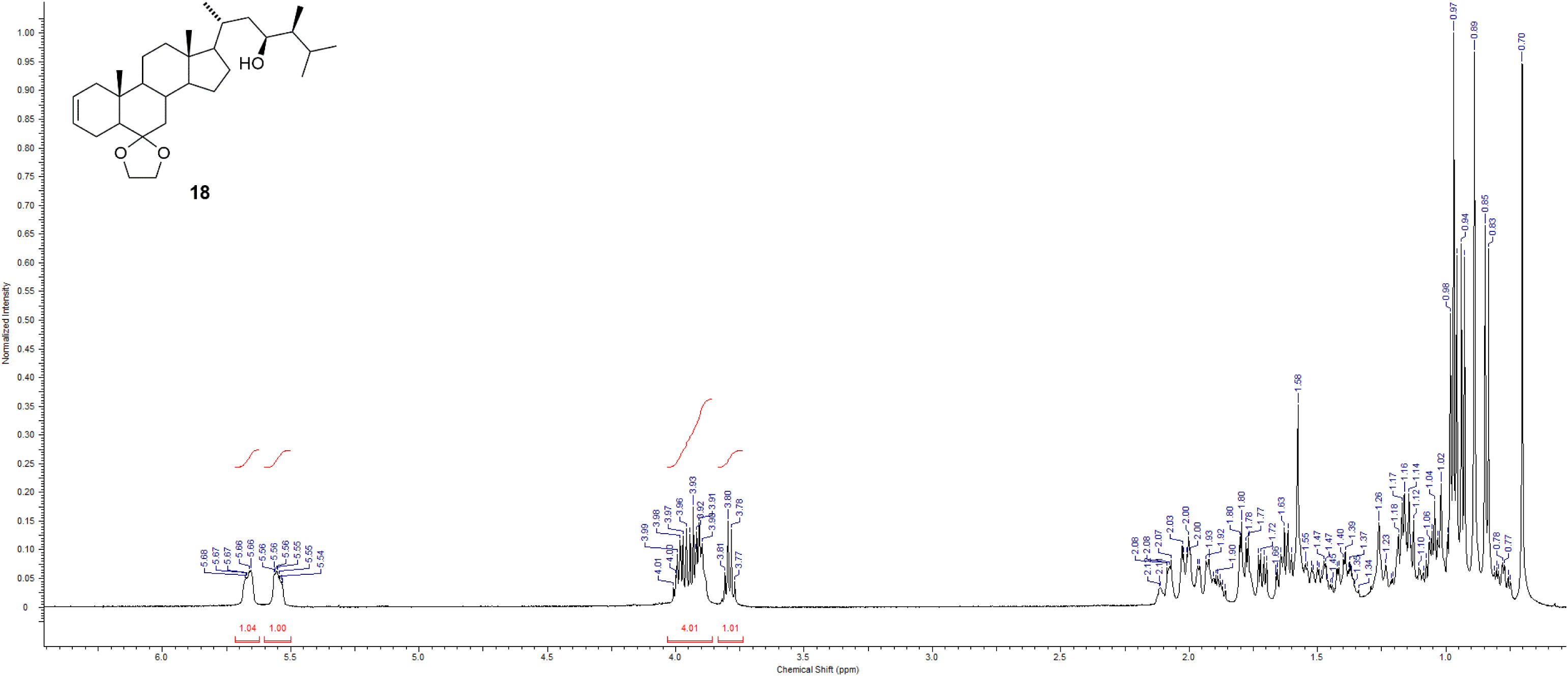
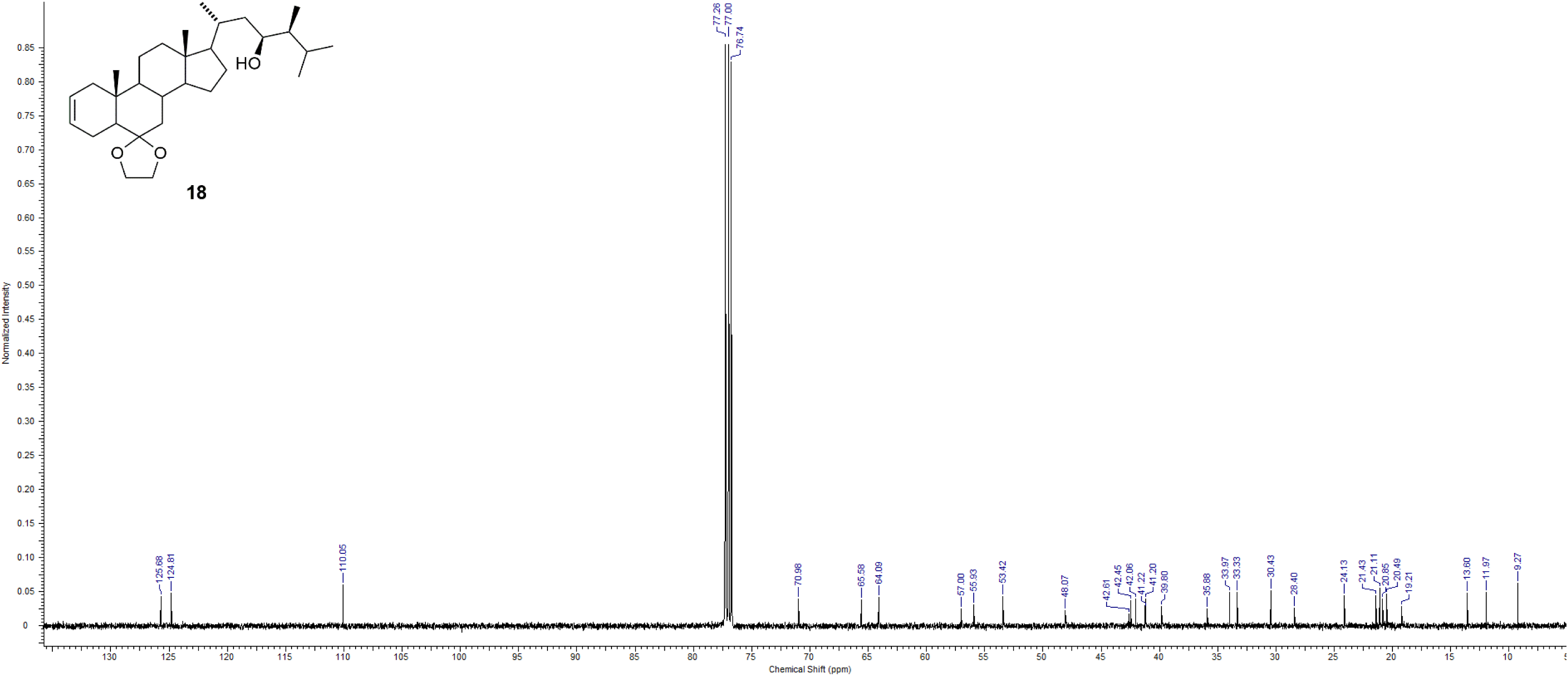
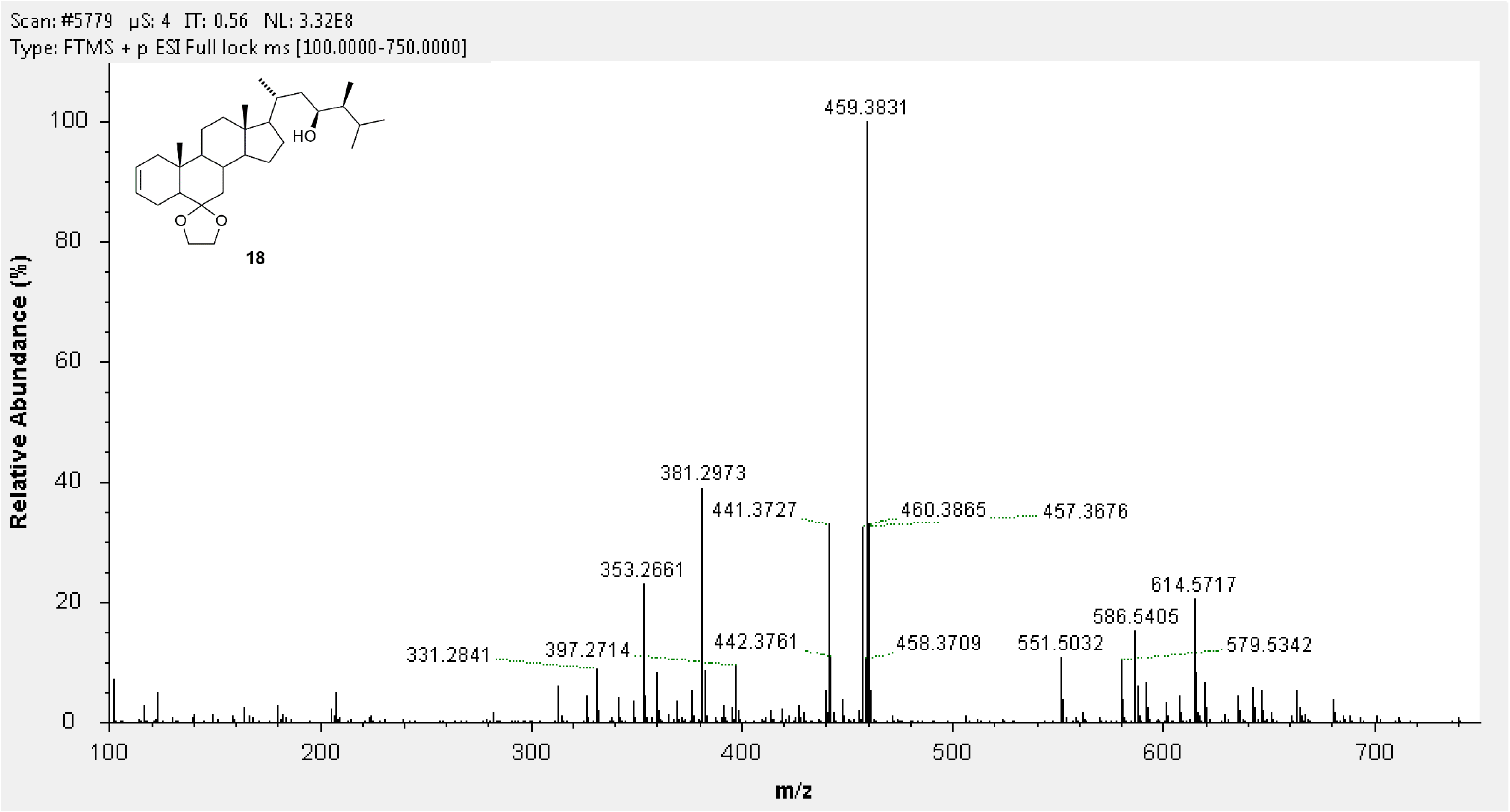
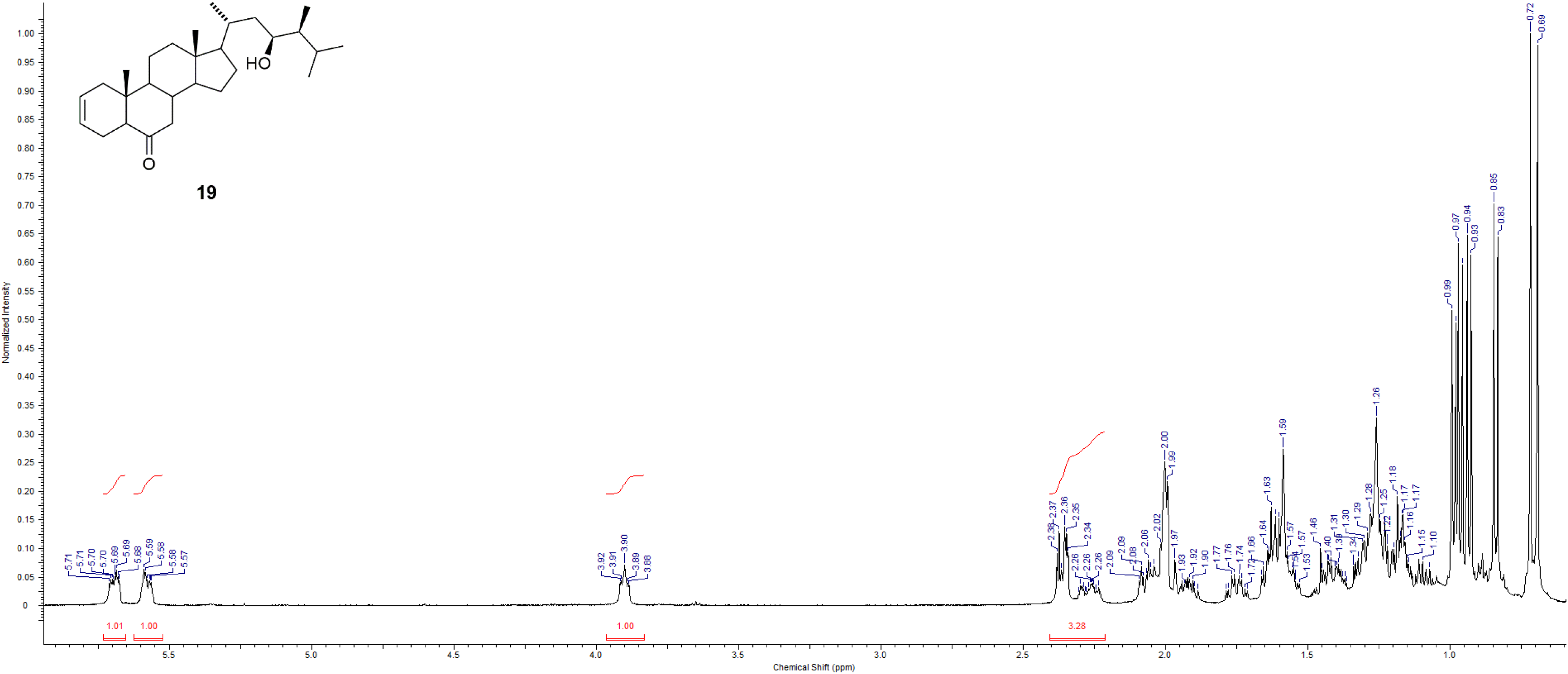
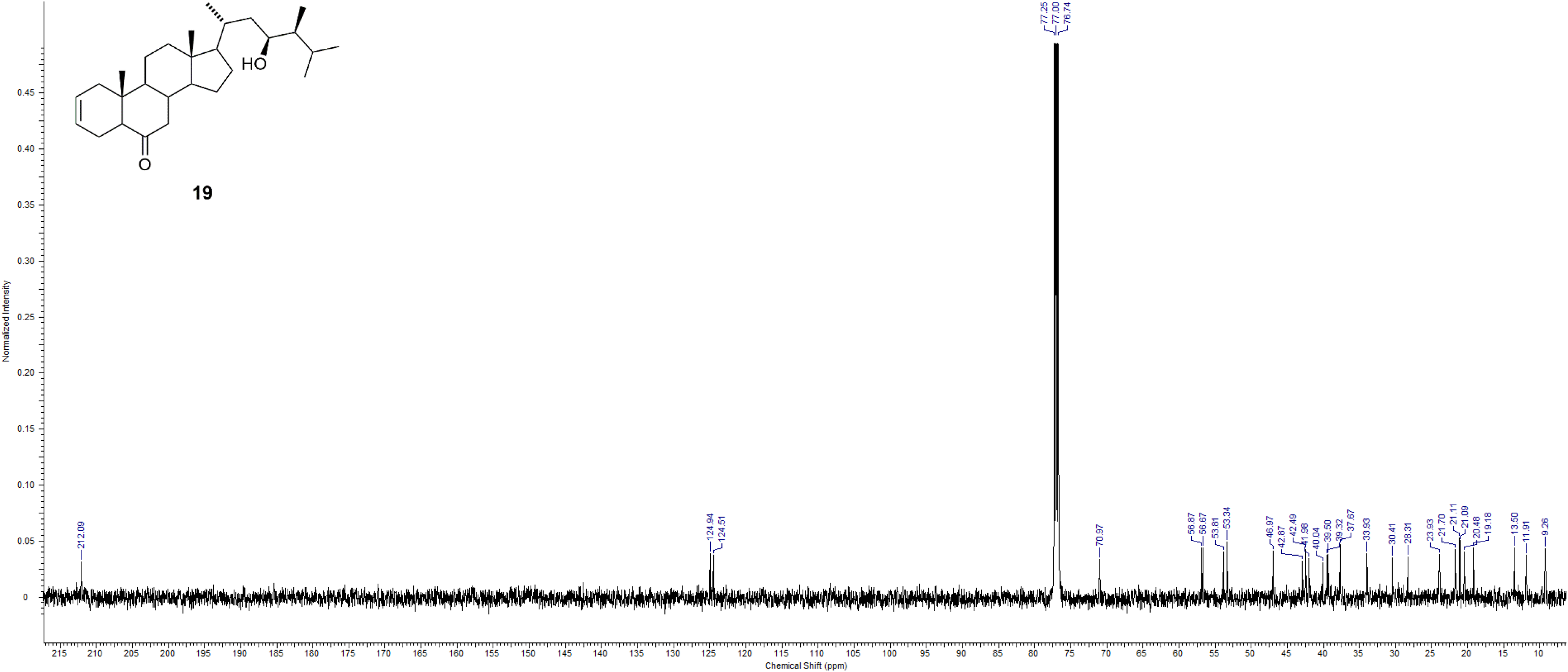
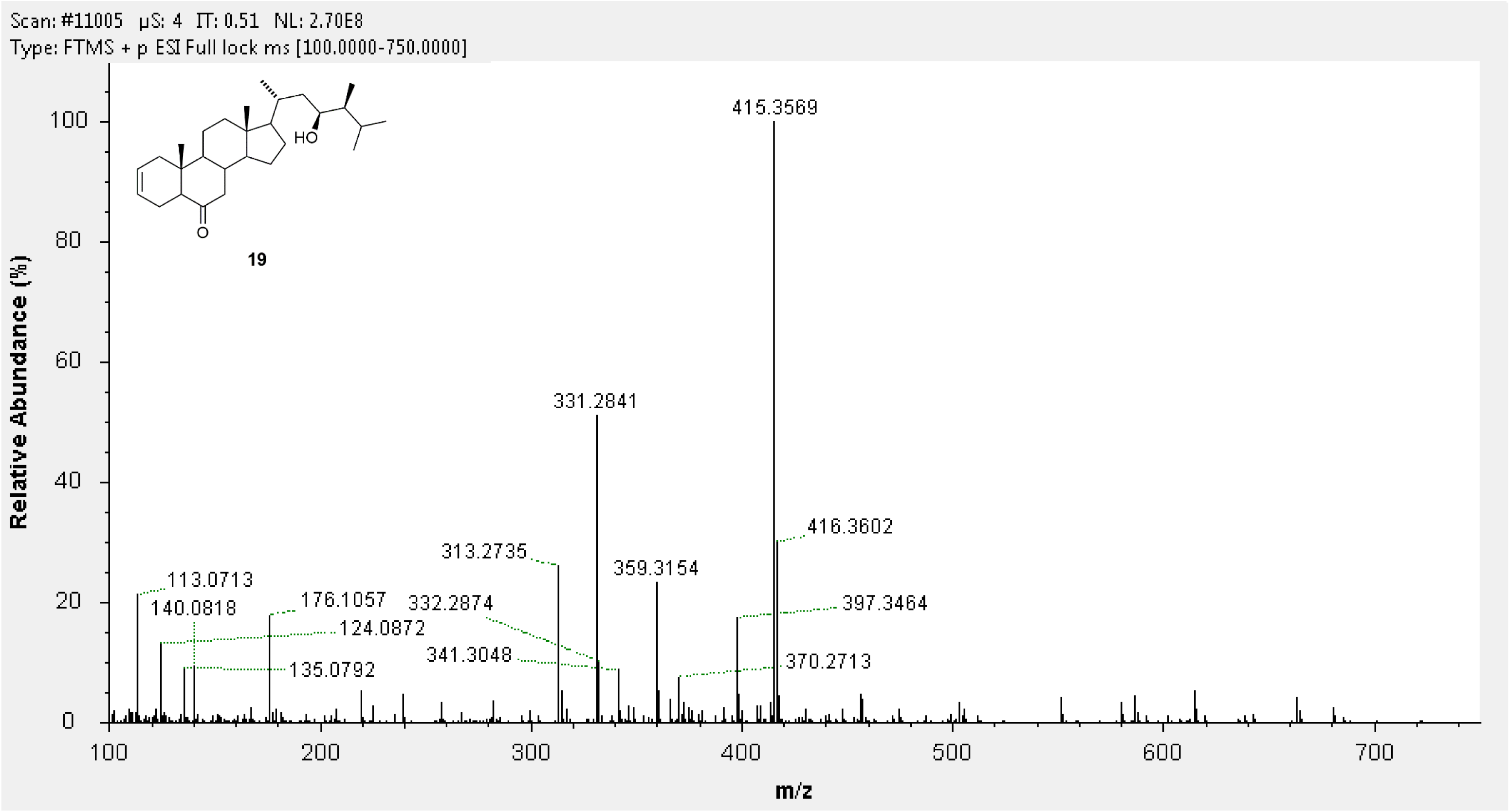
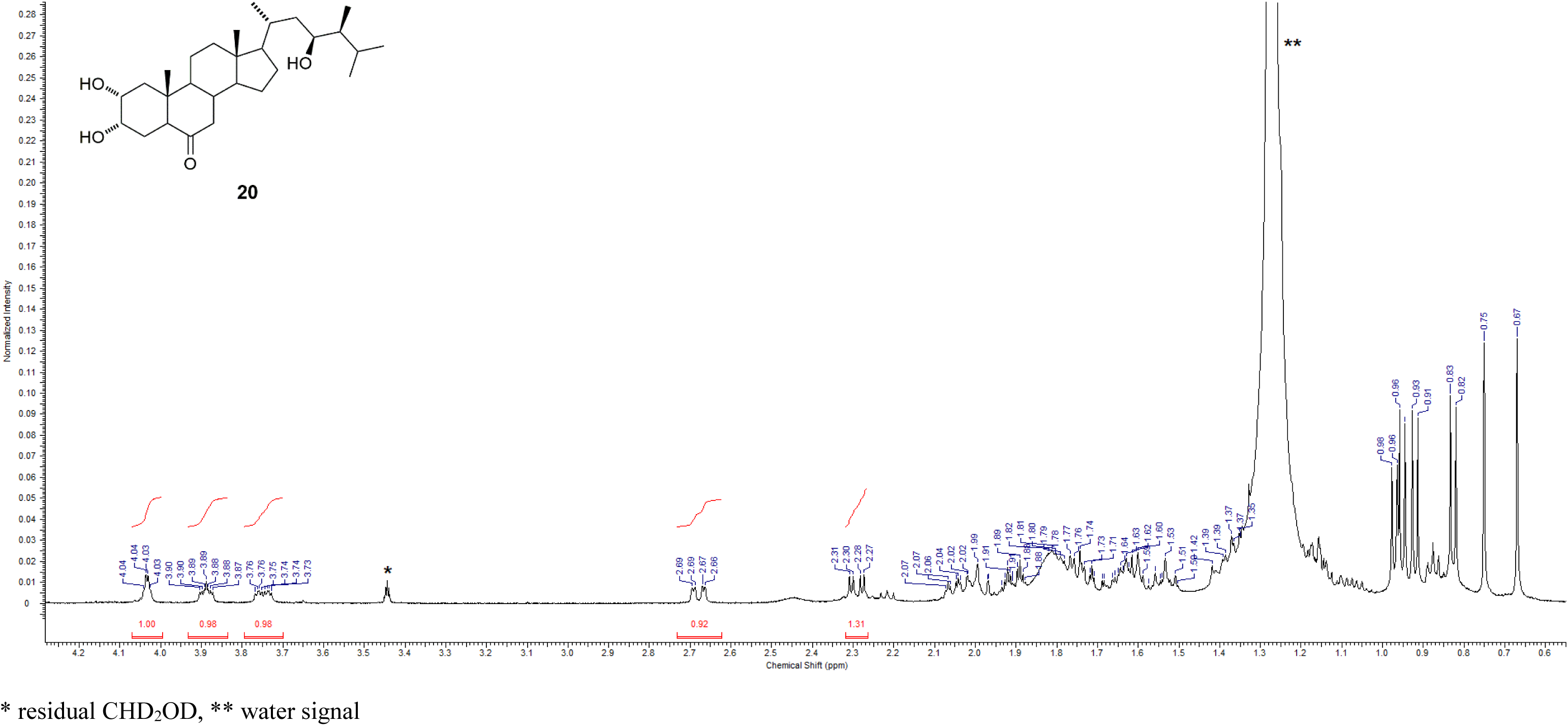
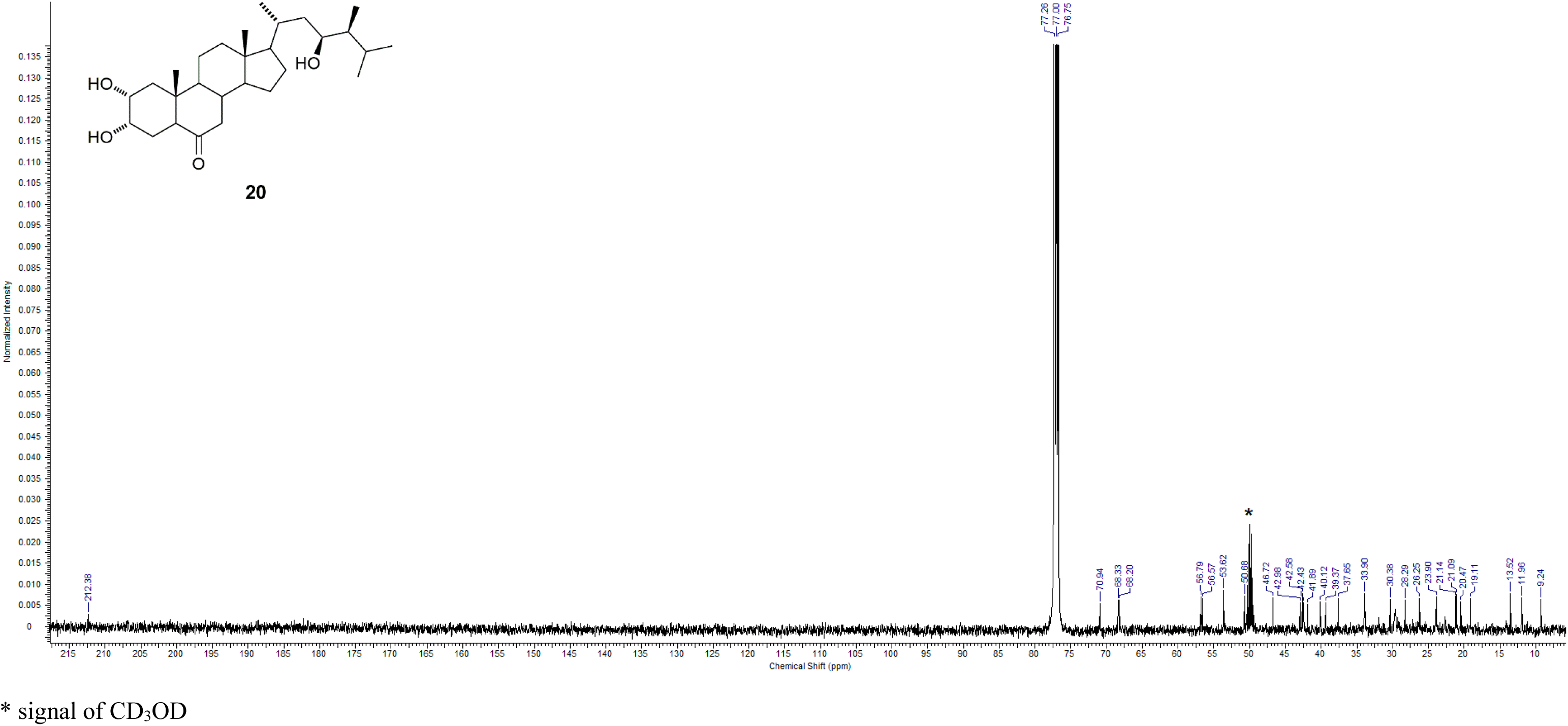
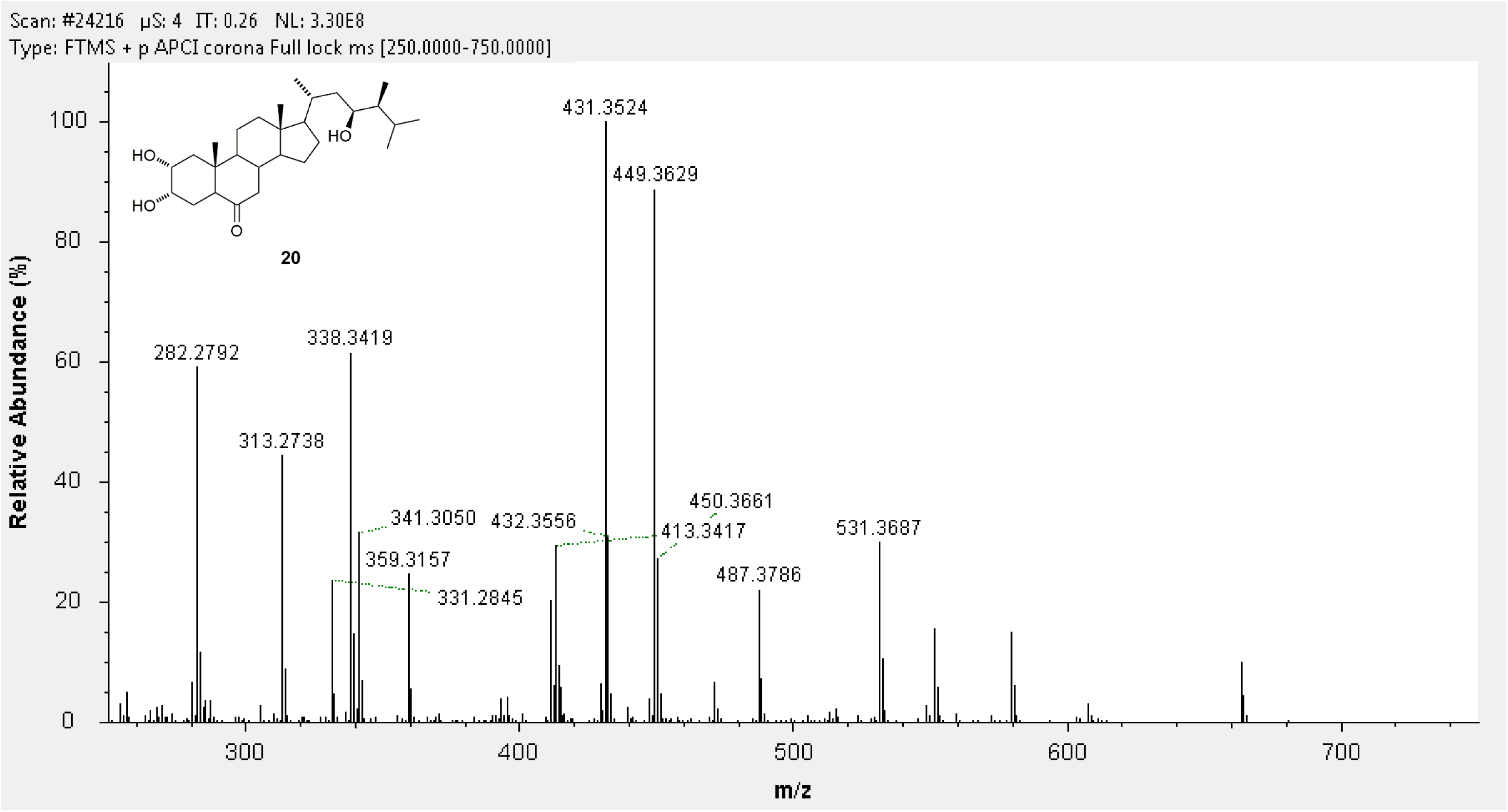

